# CDSeq2 integrates individual-level cell-type expression deconvolution with recoverability-guided prioritization

**DOI:** 10.64898/2026.09.14.751599

**Authors:** Kai Kang, Kaiyu Xie

## Abstract

CDSeq2 combines reference-informed Bayesian deconvolution with gene-type prioritization for individual-level expression analysis from bulk RNA sequencing. Brain benchmarks showed accurate RNA-fraction estimates and expression recovery comparable to BayesPrism in excitatory neurons. Prioritization enriched accurately recovered profiles and supported continuous-phenotype and cell-type-specific expression quantitative trait locus (eQTL) analyses, with the strongest application evidence in excitatory neurons. Forty independently generated synthetic cohorts with local linkage disequilibrium evaluated a frozen scorer. In the principal stronger-effect setting, retaining 40% of candidate profiles increased eQTL precision from 77.6% to 92.6%. CDSeq2 delivers complete expression estimates with optional prioritization to support focused biological follow-up.

## 1 Background

Bulk RNA sequencing measures the combined transcriptional output of multiple cell populations. Large bulk cohorts offer opportunities to study expression variation across individuals, but an association observed in tissue can reflect changes in cellular composition, expression within a cell type, or both. Recovering individual-level cell-type expression can make these cohorts more informative for quantitative-trait and genetic-association studies.

A large deconvolution literature has established effective approaches for estimating cellular composition from bulk expression. CIBERSORT uses support-vector regression to estimate cell fractions, MuSiC incorporates multisubject single-cell information, and Bisque learns transformations between single-cell and bulk measurements [1, 2, 3]. SCDC combines multiple single-cell references, DWLS uses dampened weighting, and Scaden learns composition from simulated mixtures [4, 5, 6]. CDSeq additionally estimates shared cell-type expression profiles without requiring a reference [7]. These developments provide valuable information about cellular abundance, but estimating a cell fraction or a shared reference profile does not recover a separate expression profile for every individual and cell type.

Individual-level expression deconvolution is addressed by a more limited subset of methods. Important examples include CIBERSORTx high-resolution expression imputation, bMIND, BayesPrism, BLADE, TAPE and, more recently, BLUE [8, 9, 10, 11, 12, 13]. These methods demonstrate the feasibility of sample-specific expression recovery using distinct statistical or learning-based assumptions. The opportunity for CDSeq2 is therefore to connect individual-level expression inference to an explicit, phenotype-independent assessment of which gene/type profiles preserve the variation needed for association analysis. This complements the expression-estimation objective by making recoverability a directly evaluated component of the analysis.

This challenge arises because the mixture problem is underdetermined. Even with well-estimated RNA fractions, several configurations of cell-type expression can explain similar bulk counts. Informative reference priors regularize inference, but do not establish likelihood identifiability for every gene and cell type. Genes with overlapping reference expression, limited read support or strong concentration in a small number of donors may have different recovery properties. Accurate mixture reconstruction or average expression therefore does not establish accurate donor-to-donor expression variation.

We developed the CDSeq2 software package to connect individual-level expression inference with an explicit assessment of variation recovery. A reference-informed Bayesian model estimates expression in each sample and cell type. An optional six-feature generalized additive model (GAM6) predicts the recoverability of each gene/type profile using observable features of the bulk data, reference and estimated RNA fractions. Users can retain the complete estimates or prioritize a subset for downstream analysis. The scorer does not use phenotypes, genotypes or association statistics. We evaluate expression inference against known components and assess the additional value of prioritization using directly aggregated human brain counts, synthetic continuous phenotypes, nominated-variant expression quantitative trait locus (eQTL) simulations and a frozen-rule simulation with independent synthetic cohorts and haplotype-based local linkage disequilibrium (LD). The three-workflow comparison separates baseline inference performance from the effect of adding GAM6, with accuracy and discovery coverage reported together. The principal application evidence comes from independent synthetic cohorts evaluated with a frozen rule; additional equal-budget controls assess the value of learned prioritization beyond simple support ranking.

## 2 Results

Supplementary methods and Figures S1–S18 are provided in Additional file 1. Machine-readable data are grouped as Tables S1–S11 (Additional file 2), S12–S19 (Additional file 3), S20–S25 (Additional file 4), S26–S35 (Additional file 5) and S36–S42 (Additional file 6).

### 2.1 A software framework for expression inference and optional prioritization

CDSeq2 takes bulk count data and an annotated single-cell or single-nucleus reference and returns individual-level cell-type expression estimates and RNA fractions. Its optional variation scorer ranks gene/type profiles using six support and variability features (Fig. 1): mean reference-weighted contribution, expected read support, reference specificity, bulk log-counts-per-million (log-CPM) variability, maximum donor read share and correlation between bulk CPM and estimated RNA fraction. A separate generalized additive model is fitted for each cell type to predict across-donor expression recovery. Selection changes which profiles are prioritized for interpretation; it does not refit or alter the underlying expression estimates.

**Figure 1:**
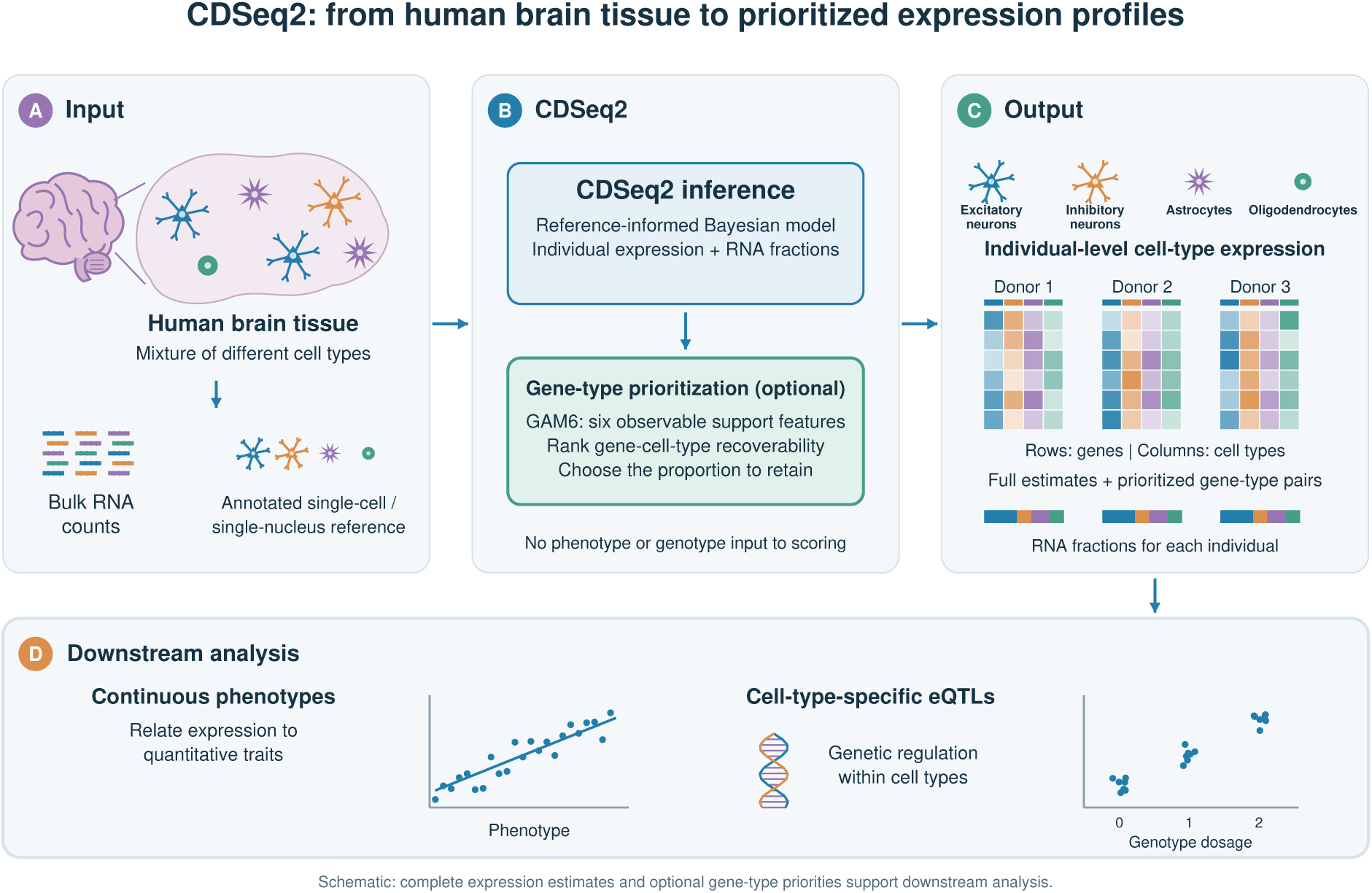
Input, CDSeq2 (with gene-type prioritization) and output for individual-level expression analysis. (A) Human brain tissue contains multiple cell types. Short reads colored by their schematic cell-type origins represent bulk RNA; the annotated reference reuses the neuron and glial shapes shown in the tissue and output. Bulk RNA counts from individual donors and an annotated single-cell or single-nucleus reference provide complementary inputs. (B) Reference-informed CDSeq2 inference estimates individual-level expression and RNA fractions; optional GAM6 scoring uses six observable support features to rank gene/type profiles by predicted across-donor recoverability. No phenotype or genotype is supplied to the scorer. (C) Each donor has a gene-by-cell-type expression matrix. Column colors match the illustrated cell types consistently across tissue, cell icons, expression matrices and RNA-fraction bars; within-column shading is illustrative expression intensity. Complete estimates and optional prioritized gene/type pairs are available without refitting inference. (D) Downstream continuous-phenotype and eQTL analyses assess association accuracy together with discovery coverage. Cartoons and mini-plots are schematic, not measured data. Model development and evaluation are described in Supplementary Section 1.6. Visualization code was prepared with OpenAI Codex assistance. Underlying data are provided in Supplementary Tables S1, S12 and S16–S18 for the illustrated benchmark evaluations.

We distinguish three workflows throughout the comparison: standard BayesPrism with the specified native input preprocessing, CDSeq2 (without gene-type prioritization), and CDSeq2 (with gene-type prioritization). Gene-type prioritization denotes optional selection using the six-feature GAM6 scorer; it does not change the inferred expression values. We use CDSeq2 alone to name the software package and the expanded workflow names to distinguish comparisons. Standard BayesPrism denotes inference without post-inference reliability selection, using the settings recorded in Methods; it does not imply that every package option or tutorial example was used. The first two workflows compare expression inference, while CDSeq2 (without gene-type prioritization) versus CDSeq2 (with gene-type prioritization) measures the incremental value and coverage cost of the package’s prioritization functionality. BayesPrism receives no output reliability selection.

The target is the Pearson correlation between inferred and observed component expression after normalization within each donor and cell type over the modeled gene panel. This target addresses relative expression variation, rather than absolute RNA abundance. Spearman correlation provides a complementary measure of donor ordering. The score is a prediction of profile recovery and is not a probability that a downstream association is correct.

We evaluated 2,000 genes across seven broad cell types using 100 nonoverlapping donors: 20 supplied the reference, 40 supplied training targets and 40 supplied evaluation profiles. The reference contained 39,533 nuclei. Evaluation bulk samples were sums of observed donor/type components, rather than new multinomial simulations. Training and evaluation donors were disjoint. Supplementary Section 1.6 describes model development and the distinction between the earlier evaluation and the separate frozen-rule simulation.

### 2.2 Accurate RNA fractions and competitive expression recovery in excitatory neurons

CDSeq2 accurately recovered known RNA fractions in the direct-aggregation benchmark: across-donor correlations ranged from 0.925 to 0.996 across the seven cell types, with mean absolute errors of 0.15–3.29 percentage points (Supplementary Fig. S3; Table S12). These estimates recover the mixture contributions that support individual-level expression inference.

We compared CDSeq2 (without gene-type prioritization) and standard BayesPrism against known components in nine simulated scenarios (Supplementary Fig. S4A–C; Tables S12–S14). BayesPrism’s specified native preprocessing retained 1,969–1,971 genes per scenario. Both methods and truth were evaluated on this common panel, with CDSeq2 RNA fractions rebased using its inferred expression. Median across-donor expression recovery in excitatory neurons was similar for CDSeq2 and BayesPrism (0.849 and 0.840). This provides a competitive baseline for evaluating the added value of gene-type prioritization; the complete cell-type comparison is provided in the supplement.

RNA-fraction accuracy further supported CDSeq2: averaged across scenarios, it had higher correlations in four of seven cell types and lower mean absolute errors in four. In excitatory neurons, fraction correlation was 0.979 versus 0.781 for BayesPrism, and mean absolute error was 3.58 versus 4.19 percentage points. Together, these results demonstrate accurate mixture estimation and strong individual-level expression recovery in excitatory neurons.

### 2.3 Gene-type prioritization enriches more accurately recovered expression profiles

Across evaluable excitatory-neuron profiles, selecting the top 400 of 2,000 genes using GAM6 increased median Pearson recovery from 0.874 to 0.978 (Fig. 2). Selected-profile medians were 0.758 in oligodendrocytes, 0.631 in inhibitory neurons and 0.551 in astrocytes. The violin plots show enrichment across the recovery distributions, with cell-type-specific counts and medians in Supplementary Table S11.

**Figure 2:**
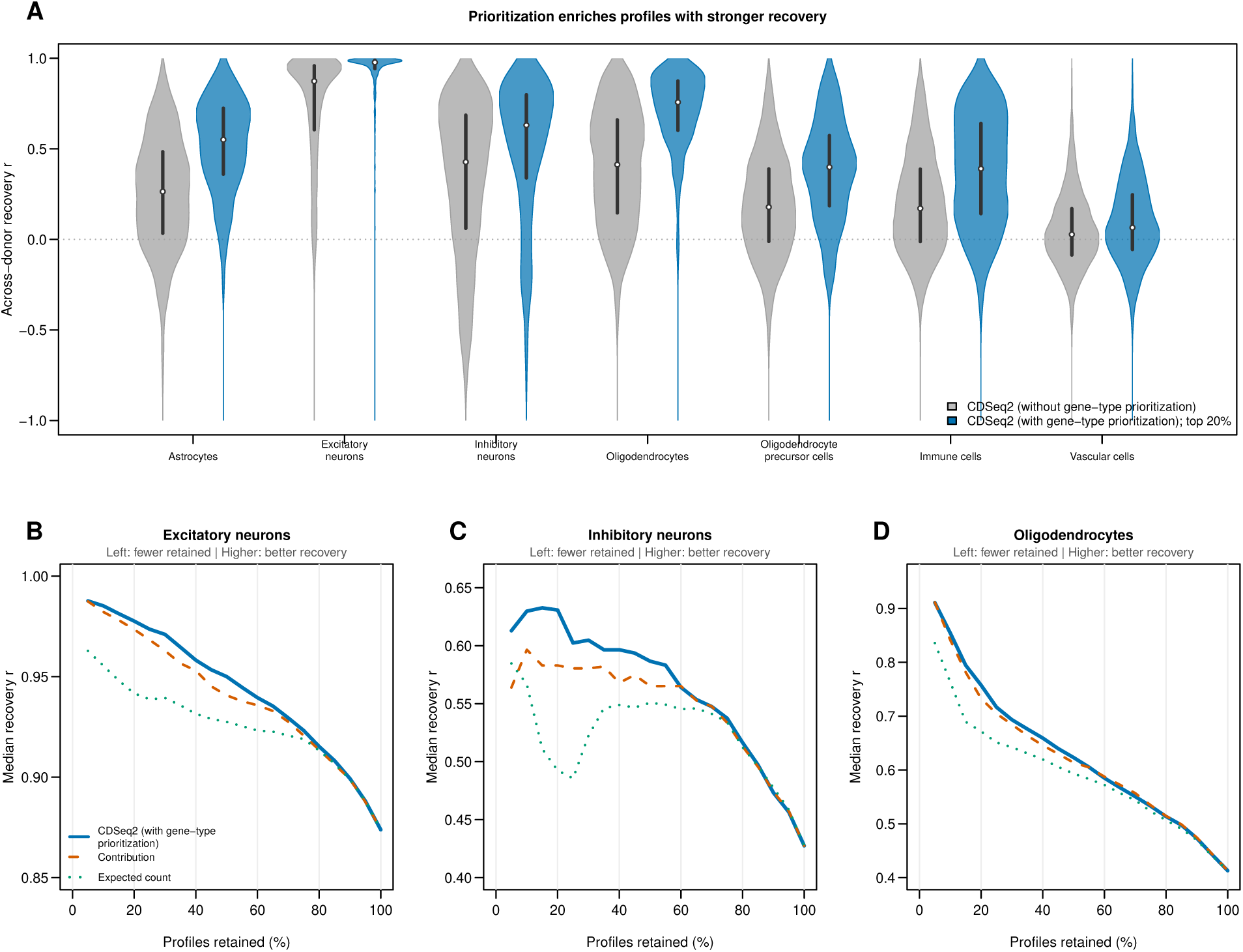
CDSeq2 (with gene-type prioritization) prioritizes more accurately recovered expression profiles. (A) Paired violins compare CDSeq2 (without gene-type prioritization) across all evaluable profiles (gray) with CDSeq2 (with gene-type prioritization) retaining the top 20% in each cell type (blue). The upward shift of the blue distributions indicates enrichment for profiles that more accurately preserve donor-to-donor expression variation. It reflects selection of existing estimates, not a refit or a change to each estimate. White points mark medians and thick lines mark interquartile ranges; equal maximum widths show density, not sample size. There are 2,000 candidate genes and 400 selected genes per type; undefined correlations are omitted. (B–D) Median recovery versus retention for excitatory neurons, inhibitory neurons and oligodendrocytes compares CDSeq2 (with gene-type prioritization) with contribution and expected-count rankings. Moving left retains fewer profiles; a higher curve at the same retention indicates more effective prioritization. At 100% retention, all profiles are retained. Panels use different expanded vertical scales to show differences; Supplementary Fig. S1 provides a common scale. Results use 40 evaluation donors; model development and complete cell-type performance are described in Supplementary Section 1.6. Visualization code was prepared with OpenAI Codex assistance. Underlying data are provided in Supplementary Tables S1, S2 and S11.

The same three-workflow comparison in nine simulated scenarios supported the added value of prioritization (Supplementary Fig. S4C; Tables S23 and S24). GAM6 retained 393–394 of 1,969–1,971 common-panel profiles per scenario and cell type, or 3,545 of 17,730 pooled profiles per type (20.0%). Pooled median excitatory-neuron recovery was 0.840 for BayesPrism, 0.850 for CDSeq2 (without gene-type prioritization) and 0.975 for CDSeq2 (with gene-type prioritization). Selected medians exceeded both unselected medians in all seven cell types. This evaluates the quality of a prioritized subset of existing estimates; the complete expression output remains available.

We also compared GAM6 with contribution ranking, which orders gene-type pairs by the mean reference-weighted fraction of a gene’s bulk expression attributed to that cell type across donors, combining reference expression with estimated RNA fractions. Expected-count ranking instead orders pairs by their corresponding expected read support. At the same 20% retention, GAM6 achieved higher median recovery than contribution ranking in five of seven types: astrocytes, excitatory neurons, inhibitory neurons, oligodendrocytes and oligodendrocyte precursor cells. Selected medians were 0.551 versus 0.507 in astrocytes, 0.631 versus 0.583 in inhibitory neurons and 0.399 versus 0.333 in oligodendrocyte precursor cells. Both approaches selected highly recovered excitatory-neuron profiles, with medians of 0.978 and 0.973. These comparisons demonstrate the value of integrating multiple support features at a shared selection budget (Fig. 2B–D; Supplementary Fig. S1; Table S1). Full cell-type results and interpretation are provided in Supplementary Section 1.6.

### 2.4 Prioritization enriches continuous phenotype association agreement

Cell-type-specific continuous-trait analysis can localize expression programs associated with quantitative disease severity or other graded phenotypes in bulk cohorts. We evaluated whether inferred individual variation preserves these associations using 189 simulated signal phenotypes spanning seven source cell types, three expression programs per type, three signal fractions and three noise repetitions. Rankings were fixed before phenotype generation. The 39-donor sensitivity analysis used the same expression estimates and selections for all phenotypes; cohort selection and full 40-donor comparisons are detailed in Supplementary Section 3.5 and Table S25.

Figure 3 illustrates donor-level expression and association-effect recovery for the first excitatory-neuron program at signal fraction 0.50 and noise repetition 1. The archive identifier trait_031 labels this simulated phenotype. JPT1 was chosen as the highest frozen-GAM6-scoring gene in that program, independently of its association fit. Its CDSeq2 and observed-component slopes were 0.39 and 0.38 per original-cohort phenotype standard deviation. Across all 2,000 excitatory-neuron genes, slope concordance was 0.787; among the fixed 400 prioritized genes, it was 0.992. This example illustrates how prioritization can concentrate profiles that preserve phenotype-related expression variation.

**Figure 3:**
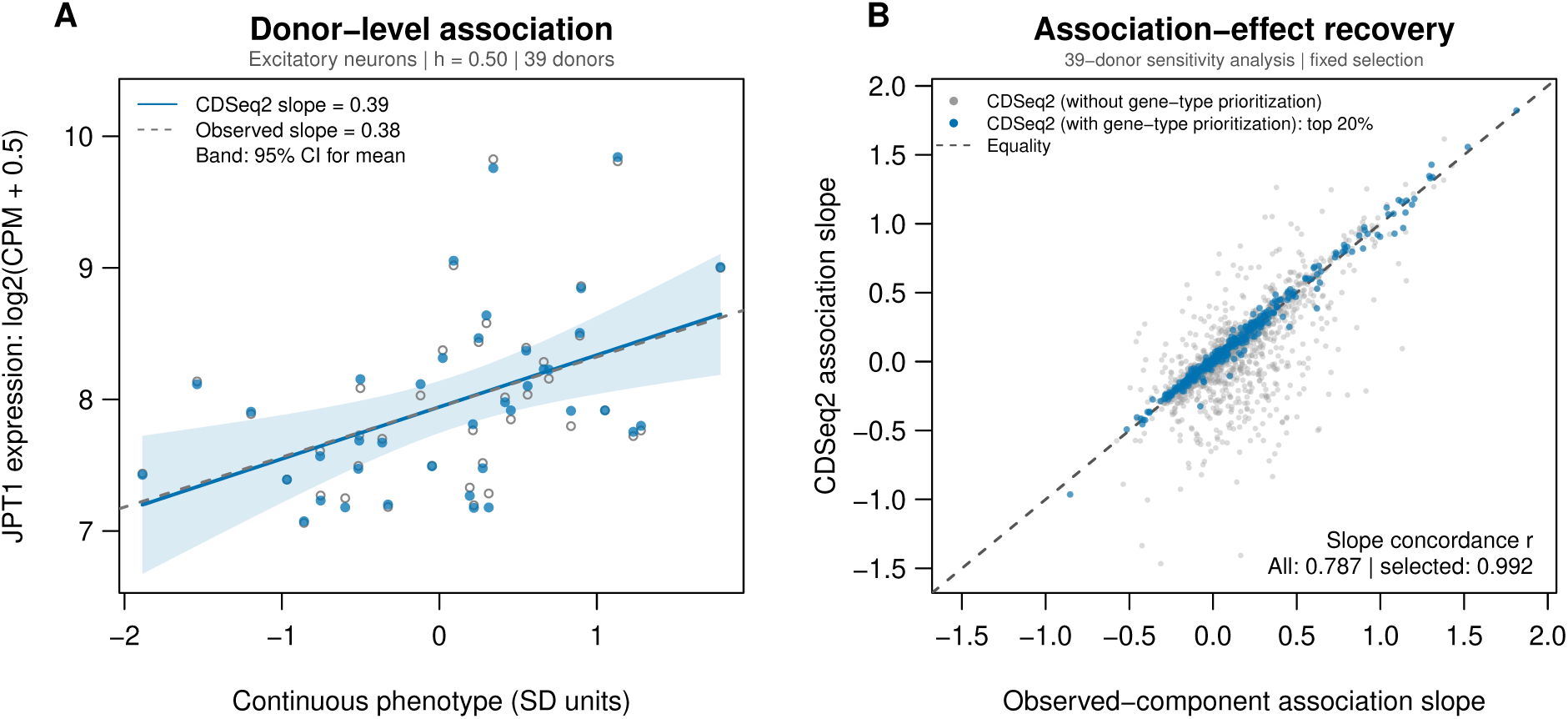
Individual-level continuous-phenotype association recovery. Both panels show the 39-donor sensitivity analysis of the first excitatory-neuron expression program at signal fraction 0.50 and noise repetition 1, archived as `trait_031`. (A) JPT1 expression versus phenotype: blue points and a solid line show CDSeq2 estimates and their regression; open gray points and a dashed line show observed components. The blue band is a pointwise 95% confidence interval for the fitted mean, conditional on estimated expression. JPT1 was chosen by its highest frozen GAM6 score among the program genes, independently of association fit. (B) Estimated versus observed-component association slopes for all 2,000 excitatory-neuron genes (gray), with the fixed top-20% prioritized subset highlighted in blue. The dashed diagonal marks equality; slope concordance is 0.787 overall and 0.992 among prioritized profiles. Phenotype units retain the original 40-donor standardization. Expression inference and scoring are unchanged. Cohort selection and influence diagnostics are detailed in Supplementary Section 3.5; Table S25 contains the full 40-donor result and every single-donor omission. Observed-component slopes are empirical reference estimates. Visualization code was prepared with OpenAI Codex assistance. Underlying data are provided in Supplementary Tables S16, S17 and S25.

At a shared 20% profile budget, GAM6 achieved higher agreement and full-type reference recall than contribution and expected-count ranking for excitatory-neuron programs at signal fractions 0.50 and 0.75 (Supplementary Fig. S7; Table S3). Agreement requires significance in both inferred and observed-component analyses with the same direction. The complete association, coverage and null results, including the interpretation of the observed-component reference, are provided in Supplementary Section 3.9, Figures S2 and S7, and Tables S3, S4 and S9.

### 2.5 Gene-type prioritization supports cell-type-specific eQTL detection

Cell-type-specific eQTL analysis connects inherited regulatory variation to the cellular populations in which it acts, supporting prioritization of genes and mechanisms for follow-up. We constructed nine nominated-variant scenarios from the 40 evaluation donors, including single-type, shared, opposite-direction and other-type effects, together with wholly null genes. The principal injection types were excitatory neurons, inhibitory neurons and oligodendrocytes. Signal ratios of 0, 0.5 and 0.8 were evaluated in three repetitions, with 254 nominated genes and 1,778 gene/type tests per scenario.

For excitatory neurons at signal ratio 0.5, top-40% prioritization retained 79 injected tests across three repetitions and correctly detected 39, giving a selected-set detection rate of 49.4%. At signal ratio 0.8, it correctly detected 75 of 86 retained injected tests, or 87.2% (Supplementary Fig. S11). These results demonstrate detection of genetic effects within the prioritized profiles. Complete discovery counts, full-target recall and attribution outcomes across cell types are reported in Supplementary Tables S5 and S6; the distinction between selected-set detection and overall coverage is detailed in Supplementary Section 3.10.

### 2.6 Expression inference and the added value of gene-type prioritization

We compared standard BayesPrism, CDSeq2 (without gene-type prioritization) and CDSeq2 (with gene-type prioritization) on the same simulated mixtures (Fig. 4A–C). Standard BayesPrism used the specified native ribosomal/mitochondrial cleanup and default bulk-outlier filtering, retaining 250–251 of 254 nominated genes per scenario, with no output reliability selection. CDSeq2 prioritization retained 101 of 254 nominated excitatory-neuron pairs per repetition, using the same expression estimates and association statistics as unprioritized CDSeq2. The comparison therefore evaluates baseline inference and the additional selection functionality within one framework.

**Figure 4:**
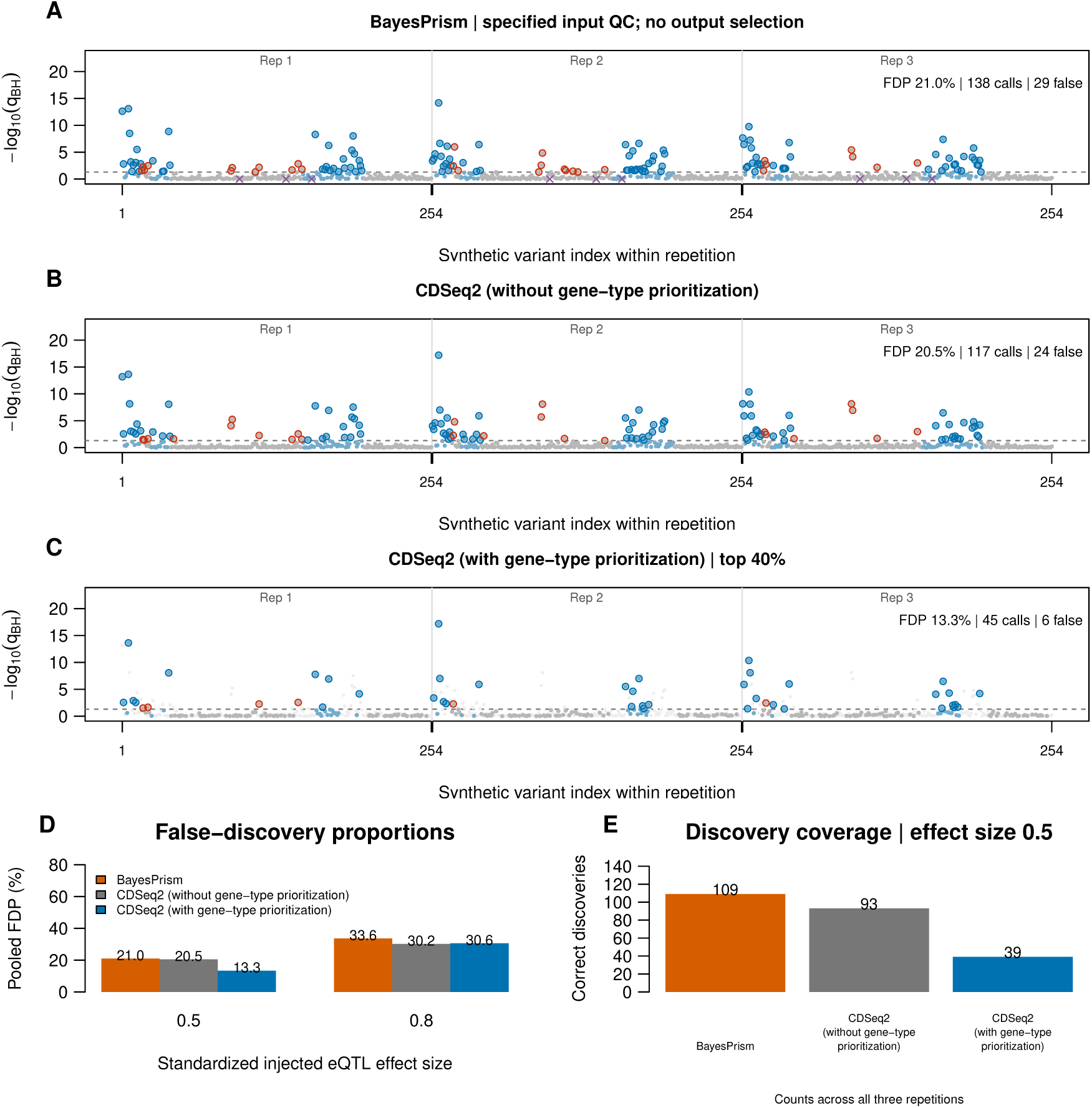
Three-workflow comparison of eQTL inference and added prioritization in excitatory neurons. Signal ratio is 0.5. Indexed Manhattan-style panels show (A) standard BayesPrism with the specified native input preprocessing, (B) CDSeq2 (without gene-type prioritization) and (C) CDSeq2 (with gene-type prioritization) top-40% selection. A and B have no output reliability selection; C retains the same CDSeq2 estimates and association statistics as B. Each panel concatenates three repetitions in the same variant order. Blue filled points indicate injected excitatory-neuron tests and gray points indicate non-injected tests. Blue outlines mark correct-sign significant calls; red outlines mark false or wrong-sign calls. In C, discarded tests are faint background points without call outlines. In A, purple baseline crosses mark preprocessing-excluded genes. The dashed line is BH-adjusted *q* = 0.05 using the original 1,778-test family. (D) Pooled FDP at both signal ratios. (E) Correct discoveries at 0.5 show the coverage cost of prioritization. CDSeq2 (with gene-type prioritization) retains 101 pairs per repetition; BayesPrism tests all 251 genes surviving input QC at ratio 0.5. Repetitions reuse 40 donors, variants lack genomic coordinates, and observed FDP does not establish nominal FDR control. The 0.8 comparison is in Fig. S8. Visualization code was prepared with OpenAI Codex assistance. Underlying data are provided in Supplementary Tables S6, S18 and S19.

At signal ratio 0.5, prioritization increased the precision of CDSeq2’s reported discoveries from 79.5% to 86.7%, compared with 79.0% for standard BayesPrism. At ratio 0.8, the corresponding precisions were 69.8%, 69.4% and 66.4%; prioritization did not improve CDSeq2 precision at that strength. Figure 4 pairs discovery accuracy with correct-discovery counts, making the reporting scope of each workflow explicit; the stronger-effect comparison is in Supplementary Fig. S8. These results support optional prioritization for focused follow-up while showing that its association benefit depends on the setting. Per-test evidence and preprocessing records are provided in Supplementary Tables S8, S18 and S19.

### 2.7 Precision–recall analysis quantifies the accuracy–coverage tradeoff

Precision–recall analysis evaluates the usefulness of a prioritized discovery set across association thresholds (Fig. 5). Correct discoveries require both the injected gene/type target and its direction, and recall uses all 246 injected excitatory-neuron targets across the three repetitions. Figure 5A–D reports precision, full-target recall and F1 together, with prioritization membership held fixed. Full operating-point summaries are provided in Supplementary Section 3.10 and Tables S20–S22.

**Figure 5:**
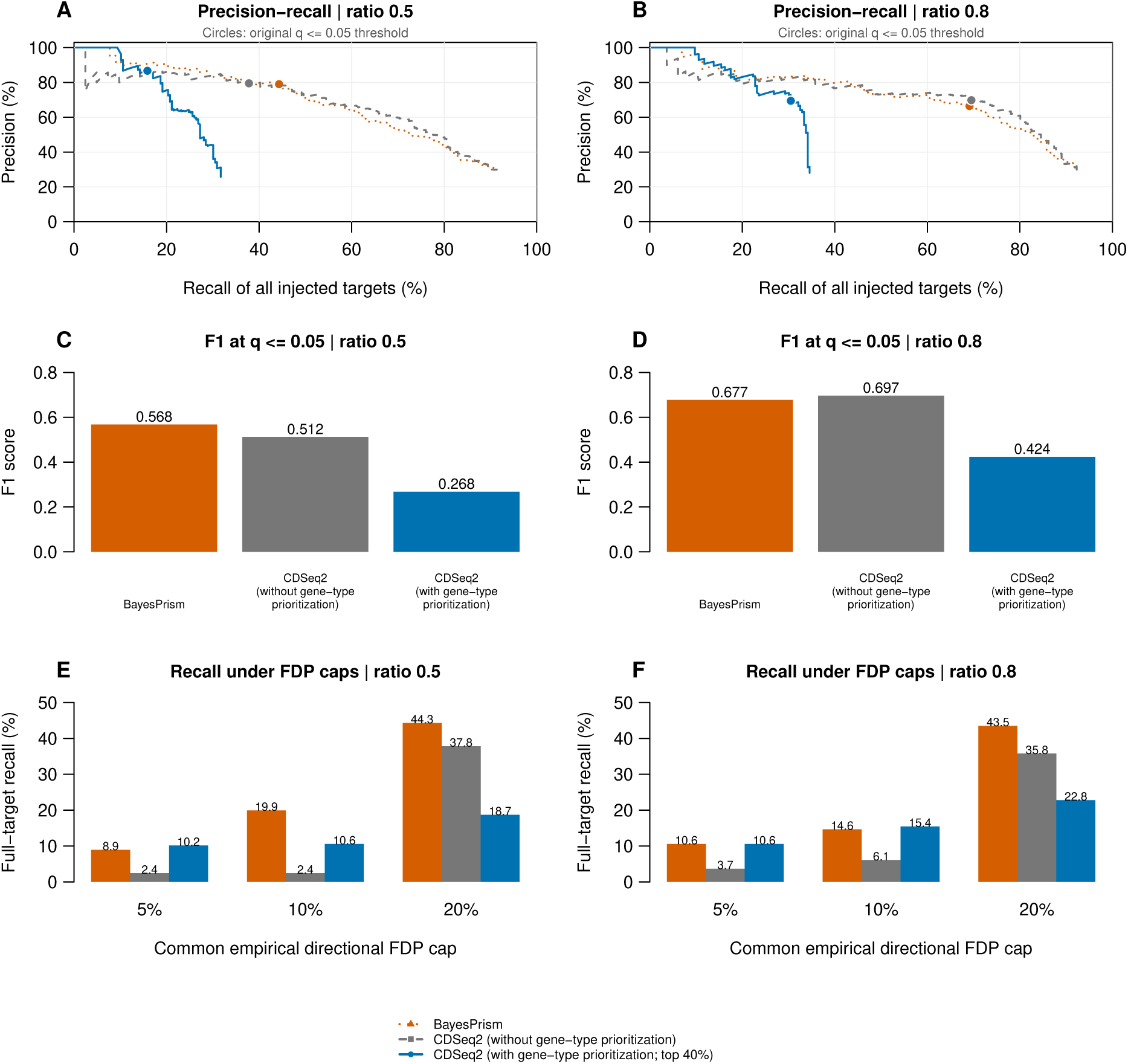
Gene-type prioritization supports precise eQTL discovery in excitatory neurons. Columns show signal ratios 0.5 and 0.8. Orange, gray and blue denote standard BayesPrism, CDSeq2 (without gene-type prioritization) and CDSeq2 (with gene-type prioritization) with fixed top-40% selection. BayesPrism uses the specified native input preprocessing; only CDSeq2 (with gene-type prioritization) applies output reliability selection. (A,B) Precision–recall curves sweep original BH-adjusted thresholds; circles mark *q ≤* 0.05. All tests tied at a threshold enter together; connecting lines guide the eye and do not imply additional attainable operating points. (C,D) F1 at *q ≤* 0.05 combines precision and full-target recall. (E,F) Maximum recall under common empirical directional FDP caps of 5%, 10% and 20% assesses discovery coverage at a shared error budget. Correct calls require the injected target and sign; all other calls contribute to FDP. Recall uses all 246 injected excitatory-neuron targets, including discarded or unavailable targets. Counts pool three repetitions using the same 40 donors. GAM6 membership stays fixed. Matched-cap thresholds use truth retrospectively and may exceed 0.05; achieved FDP can fall below the cap. This is not a prospective FDR-control procedure. Repetition-specific curves are in Fig. S10. Visualization code was prepared with OpenAI Codex assistance. Underlying data are provided in Supplementary Tables S20, S21 and S22.

Under a shared 5% empirical directional false-discovery proportion (FDP) cap, CDSeq2 (with gene-type prioritization) recovered 25 correct effects at signal ratio 0.5 and 26 at ratio 0.8, compared with six and nine for CDSeq2 (without gene-type prioritization), and 22 and 26 for standard BayesPrism. At the 10% cap, prioritization increased CDSeq2 correct discoveries from six to 26 and from 15 to 38. These results show that the scorer can concentrate detectable effects among the most precise candidate sets (Fig. 5E,F). At the 20% cap, however, the unprioritized workflows recovered more effects: 93 and 88 for CDSeq2 and 109 and 107 for BayesPrism, versus 46 and 56 after prioritization. The value of selection therefore depends on the accepted error budget and desired discovery coverage.

The full curves characterize where each workflow is most useful; the complete BayesPrism comparisons and exact achieved FDP values appear in Supplementary Section 3.10. Matched-cap thresholds are selected retrospectively using simulation truth and describe attainable performance, rather than a prospective false-discovery rate (FDR) control procedure. Repetition-specific and all-cell-type results are provided in Supplementary Figures S9 and S10.

### 2.8 Independent synthetic cohorts test prioritization under local LD

We evaluated the fixed scorer in 40 independently generated synthetic cohorts: 30 signal cohorts across six settings and ten global-null expression cohorts (Table S26). Both deconvolution methods were fitted anew. The scorer, top-40% rule and primary *q ≤* 0.05 threshold were fixed before generating evaluation outcomes, and every generated individual was retained. Public phased haplotypes supplied local linkage disequilibrium, and all common variants in each assigned window were tested. These synthetic cohorts assess recovery of directly injected gene/type effects under training-derived expression distributions and local genetic correlation.

At *n* = 120 and nominal pre-normalization genetic variance fraction *h*^2^ = 0.30, standard BayesPrism identified 74 injected excitatory-neuron effects and 25 non-injected calls across five cohorts, giving pooled precision of 74.7%. CDSeq2 (without gene-type prioritization) identified 76 injected effects and 22 non-injected calls (77.6% precision). CDSeq2 (with gene-type prioritization) reported 50 injected effects and four non-injected calls, increasing precision to 92.6%. Retaining 40% of candidate profiles preserved 65.8% of CDSeq2’s injected discoveries while retaining 18.2% of its non-injected calls. Figure 6A–D reports accuracy alongside full-target recall and F1, with all targets retained in the recall denominator.

**Figure 6:**
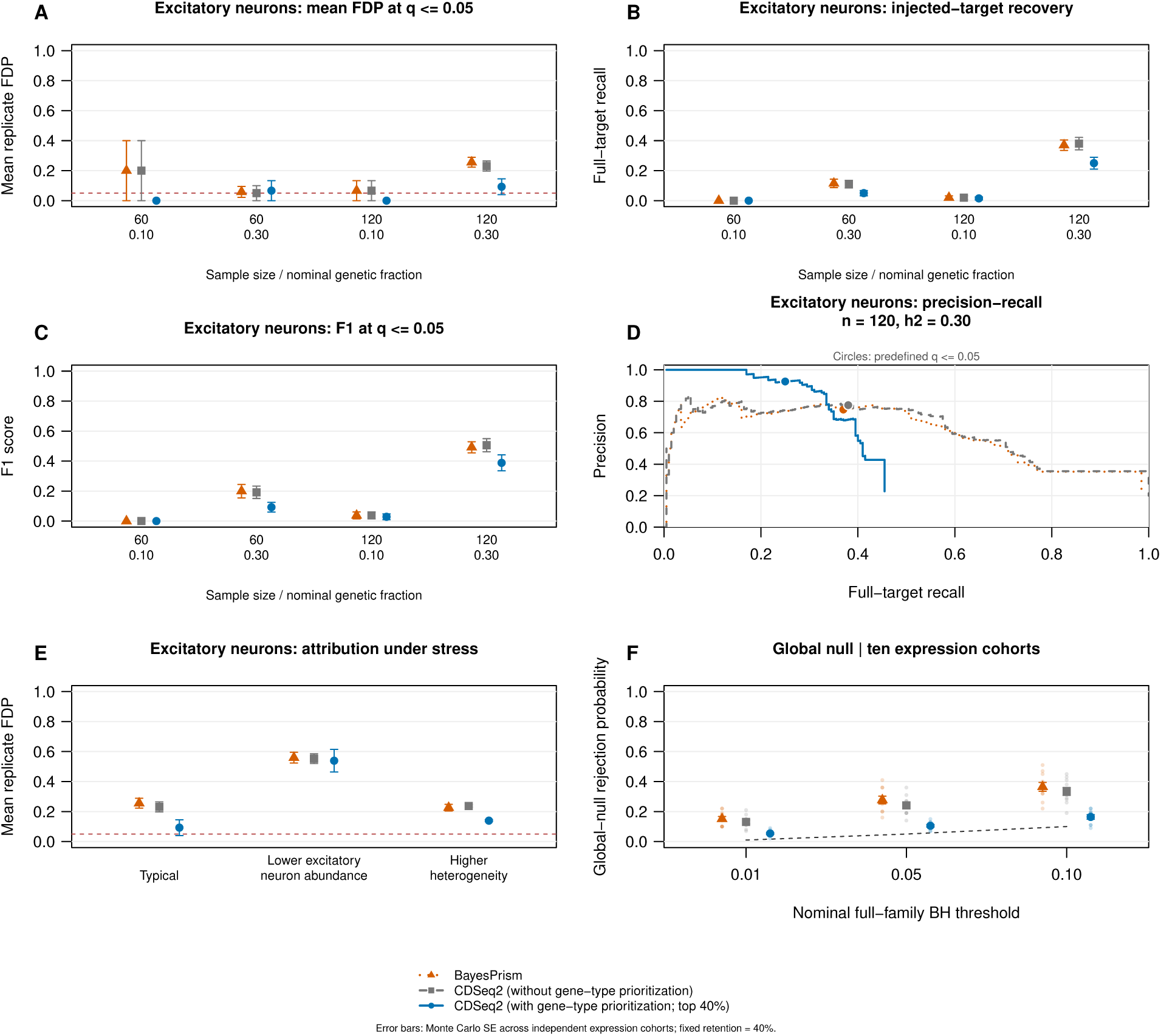
Frozen-rule evaluation in independent synthetic cohorts with local LD. Standard BayesPrism (orange), CDSeq2 (without gene-type prioritization; gray) and CDSeq2 (with gene-type prioritization; blue) use fresh inference in every cohort. The scorer and top-40% rule were fixed before test generation; BayesPrism uses native preprocessing without output selection. Each signal setting has five independent cohorts. (A–C) Mean replicate FDP, full-target recall and F1 for excitatory neurons at *q ≤* 0.05, across sample sizes 60 and 120 and nominal pre-normalization genetic variance fractions *h*^2^ = 0.10 and 0.30. Error bars are Monte Carlo standard errors. (D) Precision–recall curves pool the five *n* = 120, *h*^2^ = 0.30 cohorts at common gene-level *p* thresholds. Circles mark *q* = 0.05, which need not map to one pooled raw-*p* threshold. (E) Mean excitatory-neuron FDP under typical abundance, lower excitatory-neuron abundance and higher heterogeneity at *n* = 120, *h*^2^ = 0.30. Dashed lines in A and E mark 0.05, not guaranteed control. (F) Global-null probability of any full-family rejection at three nominal thresholds: 100 genotype realizations in each of ten independent 60-individual expression cohorts. Error bars use between-cohort standard errors; the dashed line gives nominal thresholds. Discovery denotes a directly injected gene/type effect, not the causal SNP. FDP measures direct-target attribution; normalization-induced marginal effects can fall outside the injected set. All workflows retain the 1,400-pair testing and full-target truth denominators; prioritization retains 80 of 200 candidate profiles per type. Visualization code was prepared with OpenAI Codex assistance. Underlying data are provided in Supplementary Tables S26–S31 and S35.

Expression recovery on these cohorts independently supported the scorer’s prioritization. Median excitatory-neuron correlation was 0.702 for BayesPrism and 0.703 for CDSeq2 (without gene-type prioritization), compared with 0.847 for the prioritized CDSeq2 subset. Selection retained 400 of 1,000 target/cohort profiles across the five replicates. Selected median recovery exceeded both unselected medians in all six settings (Supplementary Fig. S14). In the higher-heterogeneity setting, prioritization increased pooled eQTL precision from 76.2% to 85.9%. The prespecified abundance, effect-strength, retention and complete-null assessments define the performance range of the framework (Fig. 6E,F; Supplementary Sections 4.5–4.7, Figs. S12–S15 and Tables S28–S35).

### 2.9 Score groups and equal-budget controls clarify the prioritization benefit

Score-group diagnostics link predicted recovery to observed performance without selecting a new threshold (Fig. 7A,B; Supplementary Fig. S16). Within each cohort and cell type, profiles were divided into ten groups using the frozen score, and expression recovery and injected-target precision were evaluated separately. These displays distinguish expression predictability from association correctness and show where a high score is informative; they do not interpret the score as a calibrated probability.

**Figure 7:**
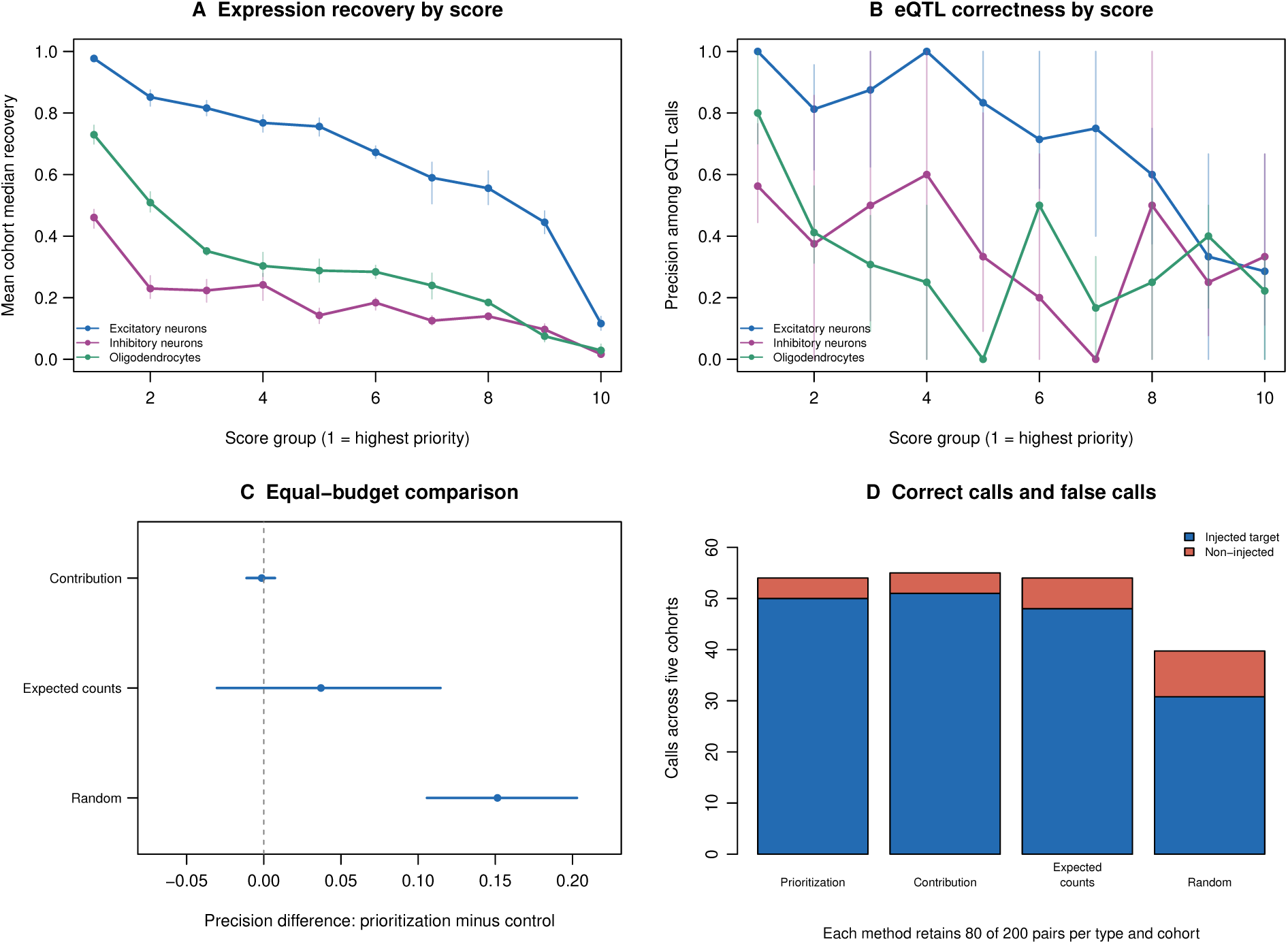
Score groups and equal-budget controls evaluate candidate prioritization. All panels use five independent cohorts at *n* = 120 and *h*^2^ = 0.30. (A) Mean cohort median expression recovery and (B) pooled injected-target precision within ten score groups for the three injection cell types. Group 1 has the highest scores. Each group has 20 candidate profiles per cohort; undefined precision when no calls occur is omitted. Bars show cohort-bootstrap 95% percentile intervals. No group boundary is chosen using truth. (C) Paired precision differences between GAM6 and equal-budget controls for excitatory neurons; intervals resample whole cohorts. (D) Correct and non-injected discovery counts across five cohorts at the unchanged full-family *q ≤* 0.05 threshold. Each control retains 80 of 200 candidate profiles per cohort; random-selection counts average 100 draws within each cohort. Contribution ranking and GAM6 perform similarly in this setting. Five cohorts give limited uncertainty resolution; intervals do not establish cross-tissue generalizability. Visualization code was prepared with OpenAI Codex assistance. Underlying data are provided in Supplementary Tables S36–S38 and S41.

At the primary 40% budget in the *n* = 120, *h*^2^ = 0.30 setting, excitatory-neuron precision was 92.6% for GAM6, 92.7% for contribution ranking, 88.9% for expected-count ranking and 77.5% for repeated random selection. The paired cohort-bootstrap precision difference between GAM6 and random selection was 15.1 percentage points (95% percentile interval, 10.6–20.3), with 50 correct discoveries versus 30.8 on average for random selection. Contribution ranking yielded 51 correct discoveries. Thus the integrated scorer provides useful enrichment beyond random retention, while this setting does not establish an advantage over contribution ranking (Fig. 7C,D). These additional comparisons preserve the original CDSeq2 estimates and full-family adjustment; no output ranking is applied to BayesPrism. All six settings, seven cell types, counts and paired uncertainty estimates are provided in Supplementary Fig. S17 and Tables S36–S38 and S41.

## 3 Discussion

CDSeq2 extends bulk RNA-sequencing analysis by combining individual-level cell-type expression inference with an explicit assessment of profile recoverability. Alongside accurate RNA-fraction estimation, it achieved expression recovery similar to BayesPrism in excitatory neurons in the evaluated benchmarks. Its additional contribution is an integrated prioritization function that identifies better-supported gene/type profiles using observable features, without requiring phenotypes, genotypes or association statistics. The complete estimates and optional priorities support both broad analysis and focused experimental follow-up.

The central application evidence is enrichment in supported cell types and settings, particularly excitatory neurons. The frozen-rule independent-cohort experiment demonstrates higher eQTL precision with lower recall, while continuous-phenotype results demonstrate agreement with observed-component associations rather than known causal correctness. Equal-budget controls show that simple contribution ranking can perform similarly to GAM6. CDSeq2 therefore adds a practical, integrated prioritization capability whose value should be assessed jointly by accuracy, coverage and cell type, rather than by a claim of universal superiority.

Individual-level deconvolution faces a common mathematical challenge: multiple cell-type expression configurations can explain the same bulk mixture. Reference quality, read support and cellular abundance influence the information available to any method. Prioritization makes variation in recoverability explicit; it does not remove non-identifiability or itself calibrate association-test error rates. The continuous-null audit identified rare large rejection bursts dominated by one donor in sparse inferred profiles; these bursts persist in the full cohort and are transformation-sensitive. The 2.5% frequency of any rejection is not itself evidence of excess global-null FDR, but the bursts reveal an association-stability concern. In the independent eQTL null benchmark, full-family rejection remained 10.5% after prioritization at a nominal 5% threshold. Prioritization therefore supports candidate enrichment without establishing nominal error control. Diagnostic details and unchanged primary analyses are provided in Supplementary Section 5. Future work will evaluate frozen scorers in independent biological cohorts and additional tissues, assess transfer across sequencing depths and reference compositions, and integrate deconvolution uncertainty into association models. These directions can extend CDSeq2’s usefulness for resolving cell-type-specific variation in large bulk cohorts.

## 4 Conclusions

CDSeq2 combines individual-level cell-type expression inference with gene-type prioritization in a unified software framework. Brain benchmarks and independent synthetic-cohort simulations demonstrate accurate RNA-fraction estimation, strong excitatory-neuron expression recovery and enrichment of accurately recovered profiles. The optional scorer supports continuous-phenotype analysis and prioritization of cell-type-specific eQTL candidates while preserving complete expression outputs. CDSeq2 makes profile recoverability accessible to users and expands the utility of bulk RNA sequencing for studying individual variation and guiding biological follow-up.

## 5 Methods

### 5.1 Data and direct aggregation benchmark

The benchmark used reconstructed human brain single-nucleus counts from nine processed Religious Orders Study and Memory and Aging Project (ROSMAP) objects accompanying the atlas of Mathys and colleagues [14, 15]. Source identities, preprocessing settings and frozen nucleus-selection manifests are supplied with the reproduction materials. The verified inputs define 20 reference, 40 training and 40 evaluation donors with no overlap. Seven annotations were used: astrocytes, excitatory neurons, inhibitory neurons, oligodendrocytes, oligodendrocyte precursor cells, immune cells and vascular cells. The fixed modeled panel contained 2,000 genes. Raw reference counts from 39,533 nuclei were used for inference. Donor/type component counts were summed to construct training and evaluation bulk counts, preserving the observed component libraries. Independent aggregation from the source files and the new selection manifests exactly recovered the training, evaluation and reference count inputs. Checks also verified panel alignment, donor separation, count nonnegativity and conservation.

### 5.2 Expression inference and reliability scoring

CDSeq2 models each individual’s bulk expression as a mixture of cell-type-specific expression profiles:

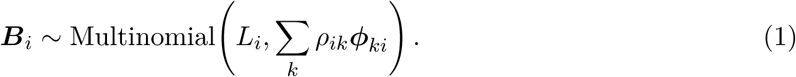

Here *L_i_* is the bulk library size, *ρ_ik_* is the RNA fraction of cell type *k*, and ***ϕ****_ki_* is its individual-specific expression profile. Reference-informed Dirichlet priors guide Bayesian read allocation. Mean allocated counts are normalized within each individual and cell type to obtain relative expression. The evaluated settings used 300 Markov chain Monte Carlo (MCMC) iterations and the last 100 allocation draws; full priors, settings and normalization equations are provided in the Supplementary mathematical methods.

Recoverability is measured by the across-donor correlation between estimated expression and known components. For each cell type, a generalized additive model predicts Fisher-transformed recovery using six observable features: contribution, expected read support, reference specificity, bulk variability, donor concentration and composition correlation. In compact form,

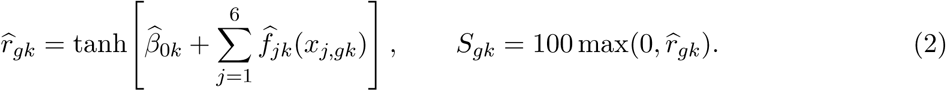

Here *x_j,gk_* denotes the transformed feature used for fitting. Rankings use *S_gk_* without phenotypes, genotypes or association statistics. Models were fitted on separate training donors, with five archived gene folds; model-development history is documented in Supplementary Section 1.6. Detailed features, transformations and fitting rules appear in the supplement. The scorer prioritizes empirically recoverable profiles; it does not eliminate the mathematical non-identifiability of unrestricted individual-level expression (Supplementary mathematical methods).

### 5.3 Continuous-phenotype and eQTL applications

Continuous phenotypes combined a standardized cell-type expression-program score with Gaussian noise at three signal fractions. We analyzed 189 signal phenotypes and 200 global-null phenotypes in a 39-donor sensitivity analysis. Phenotype values, expression estimates and frozen selections were retained from the original 40-donor cohort. Cohort selection, the full-cohort comparison and all single-donor omissions are documented in Supplementary Section 3.5 and Table S25. In each cell type, log-transformed relative expression was regressed on the phenotype. The nominated eQTL simulations instead perturbed cell-type expression according to synthetic genotype dosage, then sampled counts and recombined components into bulk mixtures. Standardized injected effect sizes of 0, 0.5 and 0.8 controlled the added genotype slope relative to baseline expression variability; they are not proportions of variance explained. All nine eQTL scenarios and the expression-recovery benchmarks retained the original 40 evaluation donors.

Association tests used an intercept and no additional covariates. Benjamini–Hochberg (BH) adjustment was performed before reliability selection, over all 14,000 gene/type tests for each continuous phenotype and all 1,778 nominated tests for each eQTL scenario. Selection did not recalculate adjusted values. Continuous-phenotype agreement required significant associations with the same sign in inferred and observed components. In eQTL simulations, correct calls required the injected direction, and false calls were significant non-injected gene/type pairs. We report the observed false-discovery proportion (FDP), rather than claiming control of the expected false-discovery rate. Detailed generation, testing and denominator definitions are in the supplement.

### 5.4 Precision–recall and empirical error-budget evaluation

Precision–recall curves summarize the tradeoff between correct discoveries and recovery of injected effects. A true positive is an injected gene/type effect called with the correct sign; every other call is a false positive. Recall uses all injected targets before filtering, and *F*_1_ = 2TP*/*(2TP + FP +FN). We report F1 at the original *q*_BH_ *≤* 0.05 threshold. These direction-aware definitions include wrong-sign errors, which are tabulated separately in the earlier non-injected-FDP summaries.

Curves sweep distinct original BH-adjusted values with GAM6 membership held fixed. For each common empirical directional FDP cap, we report the largest attainable true-positive count, breaking ties by fewer false positives and then the strictest threshold. Main results pool counts across the three repetitions; the same association cutoff applies to each repetition, whose original BH family remains unchanged. Missing or discarded targets remain in the recall denominator. Exact achieved FDP values may differ because calls are discrete. This retrospective, truth-dependent comparison measures attainable performance rather than prospective FDR control; full equations and tie handling are in the supplement.

### 5.5 BayesPrism and reproducibility

BayesPrism 2.2.3 used the documented pooled-count gene expression profile (GEP) interface with the same reference donors and one state per broad type. Native cleanup excluded ribosomal and mitochondrial gene groups (Rb, Mrp, other_Rb and chrM), with exp.cells = 1 on pooled profiles. The default bulk-outlier settings were outlier.cut = 0.01 and outlier.fraction = 0.1; pseudo.min was 10*^−^*^8^. This specified configuration did not include explicit MALAT1 or sex-chromosome exclusions, or the tutorial example’s five-individual-cell expression requirement. No tumor key or optional marker-gene restriction was used. All nine original scenarios were fitted with update.gibbs = TRUE, chain length 1,000, burn-in 500, thinning two and alpha one. Expression was extracted with get.exp from initial cell-type posterior allocations; final fractions supplied the RNA-fraction benchmark. No GAM6, contribution or other output-ranking filter was applied to BayesPrism. Association testing retained the original 1,778-test multiplicity denominator: unavailable tests entered BH adjustment as *p* = 1 and were subsequently marked unavailable, without calls. The supplement specifies preprocessing, normalization, seeds and exclusions.

The accompanying reproduction workflow separates preparation, direct inference, scorer fitting, continuous associations, eQTL simulation and comparator analysis. All results reported here were rerun using the reconstructed inputs, without historical clustering checkpoints or historical inference caches. This run included 80 direct-aggregation donor fits, 42 refitted GAM6 models, 360 nominated-eQTL donor fits, nine standard BayesPrism scenario fits and both native methods in 40 independently generated LD-aware cohorts. Inputs, settings, scripts and fit caches carry identity checks, and incompatible caches are rejected. The benchmark used the archived CDSeq2 0.0.0.9000 source with variation interface 2.4.0. The accompanying software release is 0.1.0 [16], adding a selection-summary interface, tests, model download and release metadata without changing the inference or scoring functions used in the benchmark. Exact benchmark replay uses the archived source and dependency lock separately from the current package. Numerical checks independently replayed association tests, selection, error metrics and curve operating points. A seeded generator replay and a one-donor fit from an independently installed source archive also passed. The supplementary reproduction record distinguishes the reconstructed benchmark from the earlier archived input version.

### 5.6 Independent synthetic cohorts with haplotype-based local LD

We added a frozen-rule benchmark that generates new individuals independently of the original evaluation profiles. The existing GAM6 models and a top-40% retention rule were fixed before generating the new test cohorts. No test-truth calibration or threshold optimization was used for the primary comparison. Thirty signal cohorts covered six settings with five independent realizations per setting: 60 or 120 individuals and nominal genetic variance fractions of 0.10 or 0.30 under the baseline distribution, plus lower excitatory-neuron abundance and higher-heterogeneity settings at 120 individuals and fraction 0.30. Ten independent 60-individual null-expression cohorts assessed global-null behavior. All cohorts contained 2,000 genes and seven cell types, with 200,000 modeled-panel reads per individual. These are new synthetic individuals conditional on fixed distribution parameters, not newly collected biological donors.

Local genotypes were generated by sampling pairs of whole phased haplotypes from 503 European-reference individuals in the 1000 Genomes phase-3 panel [17, 18]. Forty 25-kb chromosome-22 windows supplied local LD patterns. These windows were assigned to 200 randomly sampled, training-expression-eligible genes, without using scorer ranks; their coordinates are not the actual cis annotations of those genes. Every test-cohort single-nucleotide polymorphism (SNP) with minor allele frequency at least 0.05 in the assigned window was tested. The primary endpoint was discovery of an injected gene–cell-type regulatory effect, rather than identification of the causal SNP. This avoids labeling an LD-associated tag SNP as a false gene-level discovery.

Expression distributions were estimated from the reconstructed training components only, using cell-type-specific log-expression means, three-factor covariance approximations and residual variances. Independent factor scores, residuals, RNA fractions, genotypes and count draws generated each cohort. In signal cohorts, 40 of the 200 target genes per principal cell type (excitatory neurons, inhibitory neurons and oligodendrocytes) received one randomly selected local causal SNP; effects could overlap across cell types. No donors were removed. Native CDSeq2 and BayesPrism inference were rerun for every cohort; the frozen scorer was applied only to CDSeq2.

For each gene/type pair, we combined the two-sided SNP association tests using the Bonferroni-adjusted minimum *p* value, then applied BH across the fixed 1,400-pair family [19]. Missing or constant profiles entered adjustment as *p* = 1 and could not be called. CDSeq2 prioritization retained 80 of 200 pairs per cell type after the full-family adjustment, without changing expression or association statistics. The primary operating point was *q ≤* 0.05. We report mean replicate direct-target FDP, including zero for zero-call replicates, with between-replicate standard errors to describe simulation uncertainty [20]. Under signal settings, this measures attribution to directly injected targets; normalization-induced indirect associations mean that it need not equal the error rate for testing zero marginal association. Precision, full-target recall, F1 and retained coverage are reported separately. Null assessment additionally generated 100 independent genotype realizations per null-expression cohort; uncertainty must account for the ten expression-cohort clusters. Full generation equations, sensitivity rules and provenance are provided in the supplement.

### 5.7 Additional prioritization controls and stability diagnostics

Additional diagnostics were specified after inspecting the primary results and do not change the frozen primary experiment. Score deciles use descending finite scores within cohort and cell type, with gene-name tie breaking. Every equal-budget control retains 80 of 200 nominated profiles per type; random selection uses 100 seeded repetitions per cohort. Uncertainty uses 2,000 resamples of the five independent cohorts, preserving paired methods and all within-cohort profiles. Random repetitions are averaged within each cohort, not treated as biological replicates. Undefined precision is omitted when a resample has no calls; counts and the number of defined resamples accompany the intervals. The small number of independent cohorts limits interval precision.

Continuous-null diagnostics retain the original 200 phenotypes and compare 39- and 40-donor analyses, primary log-CPM with CPM and log-count transformations, standardized-profile principal components, and additional single-donor omissions for the three large bursts. These are sensitivity analyses, not rules for choosing donors or preprocessing from favorable outcomes. The primary cohort, normalization, association tests and selection remain unchanged. Full definitions and results appear in Supplementary Section 5.

### 5.8 Assistance with software implementation and manuscript preparation

OpenAI Codex (OpenAI) assisted with implementing additional simulation and analysis scripts, checking computational outputs, preparing visualization code, and drafting and revising manuscript text. Figure panels were rendered from the supplied numerical data and computational outputs; the workflow schematic was drawn from the described method and author-specified structure. The supplied code and source data make these outputs inspectable. Numerical results were obtained by executing the analysis code, and the design, provenance files and independent checks document the computations. Exact AI model and version records are unavailable. Both authors reviewed and verified the AI-assisted code, analyses, figures and text and take responsibility for the submitted work.

## 6 Availability and requirements

Project name: CDSeq2. Programming language: R with compiled inference code. Project homepage: the CDSeq2 repository [21]. Evaluated platform: macOS Tahoe 26.7.1 on Apple arm64 with R4.6.1. The package requires R *≥* 4.1.0 and a suitable compilation toolchain; the reproduction bundle records the full dependency versions. The full benchmark inference runs were evaluated on macOS. Automated package checks, the synthetic tutorial and scorer-loading checks additionally passed on Linux and macOS through GitHub Actions. License: MIT; no additional restrictions apply to non-academic use. The repository includes reproduction scripts, input checksums and a ROSMAP preprocessing workflow with frozen donor and gene selections. The workflow and its validation scope are described in the supplement.

## Supporting information

Additional file 1: Supplementary methods, Figures S1-S18, and table guide

## Abbreviations

BH: Benjamini–Hochberg
CPM: counts per million
eQTL: expression quantitative trait locus
FDP: false-discovery proportion
FDR: false-discovery rate
GAM6: six-feature generalized additive model
GEP: gene expression profile
LD: linkage disequilibrium
MCMC: Markov chain Monte Carlo
QC: quality control
REML: restricted maximum likelihood
RNA: ribonucleic acid
RMSE: root mean squared error
ROSMAP: Religious Orders Study and Memory and Aging Project
SNP: single-nucleotide polymorphism

## 7 Declarations

### Ethics approval and consent to participate

This study used previously published ROSMAP data downloaded from the source publication. No new participants were recruited and no new human tissue was collected. The investigators consider this secondary analysis of publicly available, deidentified ROSMAP data to be not human subjects research. A formal UNCW IRB determination was not obtained.

### Consent for publication

Not applicable.

### Availability of data and materials

The CDSeq2 package and reproduction workflows are available in the CDSeq2 repository under the MIT license [21]. Tutorials and documentation are provided in the Quarto book [22]. The version 0.1.0 software release and bundled reproduction materials are archived in Zenodo at https://doi.org/10.5281/zenodo.23489824 [16]. The ROSMAP data were downloaded from the AD/Aging Brain Atlas website accompanying Mathys et al. [14, 15]. This resource provides deidentified processed single-nucleus expression data. Complete raw data and participant metadata are available through Synapse (syn52293417 and syn3157322) [23, 24], subject to the access requirements and Data Use Certificate specified by the source repository. The accompanying reconstruction materials identify the source files, donor/gene panels and newly frozen nucleus selections used for the present rerun, together with exact source-aggregation checks. Supplementary Tables S1–S42 provide the figure and analysis data in Additional files 2–6; Additional file 1 contains the supplementary methods, figures and table guide. The independent-cohort benchmark includes its generator, compact derived inputs, public haplotype subset, CDSeq2 source snapshot and analysis scripts in the accompanying reproduction bundle. This submission uses the same reconstructed-input numerical results and additional diagnostics as the archived version 16 bundle. Submission-format changes and updated document metadata do not alter the scientific results. Table checksums match the archived files. The release includes the exact frozen brain scorer, a package-compatible copy with verified prediction replay, and an explicit account of its training context.

### Authors’ contributions

KK (Kai Kang) conceived and developed the CDSeq2 package and designed the simulations. KX (Kaiyu Xie) contributed to benchmarking. Kai Kang drafted the manuscript, and Kaiyu Xie reviewed it. Both authors have read and approved the final manuscript and agree to its submission to Genome Biology.

## Acknowledgements

We thank the ROSMAP participants and study investigators for making the underlying single-nucleus data available.

## Additional files

Additional file 1 supplies the supplementary information. Additional files 2–6 contain all 42 numbered data tables. Each ZIP includes a file list and checksums; the CSV contents are unchanged from the archived release. The source code and complete reproduction bundles are available through the archived software record [16].

**Additional file 1: Supplementary methods and figures.** `Additional_file_1.pdf`; PDF. Mathematical identifiability, detailed models, reconstruction and validation, additional diagnostics, Figures S1–S18 and the guide to Tables S1–S42.

**Additional file 2: Profile recovery and association benchmark data.** `Additional_file_2.zip`; ZIP containing CSV and TXT. Tables S1–S11: profile scores, retention curves, continuous-phenotype summaries, nominated-eQTL results, design and recovery medians.

**Additional file 3: Ground-truth accuracy and displayed application data.** `Additional_file_3.zip`; ZIP containing CSV and TXT. Tables S12–S19: RNA-fraction and expression accuracy, continuous examples, eQTL plot data and BayesPrism preprocessing.

**Additional file 4: Precision, recall and prioritization sensitivity data.** `Additional_fil e_4.zip`; ZIP containing CSV and TXT. Tables S20–S25: precision–recall curves, F1, matched empirical FDP caps, selection membership and donor sensitivity.

**Additional file 5: Independent-cohort simulation benchmark data.** `Additional_file_5.zip`; ZIP containing CSV and TXT. Tables S26–S35: LD-aware design, haplotypes, tests, accuracy, null results, expression recovery, preprocessing and injection design.

**Additional file 6: Prioritization controls and association stability data.** `Additional_file_6.zip`; ZIP containing CSV and TXT. Tables S36–S42: score groups, equal-budget controls, uncertainty, null sensitivity and donor expression support.

