## Additional file 1: Supplementary methods, Figures S1-S18, and table guide for "CDSeq2 integrates individual-level cell-type expression deconvolution with recoverability-guided prioritization"

### CDSeq2 supplementary information

Supplementary methods, eighteen figures and a guide to forty-two accompanying data tables. This document is Additional file 1. The supplementary results use the same benchmark and definitions as the main manuscript. The comparison distinguishes standard inference from the optional GAM6 prioritization functionality in the CDSeq2 package.

### 1 Supplementary methods and interpretation

#### 1.1 Scoring universe and missingness

The six-feature scorer retains the archived eligibility criteria and five gene folds. Comparison tables retain common finite-score universes. Correlation targets can be undefined when component or inferred profiles have no variation; these are excluded from recovery summaries. A high score is relative to the cell-type model and is not an absolute accuracy guarantee.

#### 1.2 Selection policies

Direct-expression and continuous-phenotype figures summarize within-type selection over the modeled 2,000-gene panel. Within-engine CDSeq2 eQTL figures select within the nominated test universe using the matched_tested_budget policy. The three-workflow comparison shows standard BayesPrism, CDSeq2 (without gene-type prioritization) and CDSeq2 (with gene-type prioritization). Standard denotes no post-inference reliability selection, using the specified inference settings and preprocessing, rather than every package default or tutorial option. Adding GAM6 preserves the CDSeq2 estimates and original association statistics; it selects which profiles are prioritized. Full-panel policies and baseline versus scenario rankings are retained in Table S5. Denominators must not be interchanged when quoting retention or detection rates. Table workflow identifiers retain their original names (BayesPrism standard, CDSeq2 before and CDSeq2 GAM6) for reproducibility; displayed workflow names use with or without gene-type prioritization.

#### 1.3 Replicates and error definitions

The 189 continuous signal phenotypes and 200 null phenotypes are reanalyzed on the same 39 donors after post hoc exclusion of original donor index 3. The nine eQTL scenarios and expression-recovery benchmarks retain all 40 evaluation donors. Repetitions vary programs, noise, genotypes and count sampling; they are conditional simulation repetitions, not independent cohorts. Continuous reference agreement is not a causal false discovery rate. In the eQTL tables, non-injected gene/type calls are false under the injection-label definition, while wrong-sign target calls are recorded separately. Normalization can create indirect effects in non-injected components, so the operational definition of error is essential.

#### 1.4 Expected and observed components

The genetic simulation supplies expected components before multinomial sampling and realized observed components after sampling. Both are analyzed separately as oracle references. These differ from the direct-aggregation benchmark, which sums the original observed components without a new multinomial draw.

#### 1.5 Reproducibility record

All numerical results in this manuscript version were regenerated from the reconstructed ROSMAP benchmark inputs. The run completed 80 direct-aggregation CDSeq2 donor fits, 42 GAM6 model fits, 189 signal and 200 null continuous phenotypes, 360 nominated-eQTL CDSeq2 donor fits and nine native BayesPrism scenario fits. Both methods were also fitted anew in forty independently generated LD-aware cohorts comprising 3,600 synthetic individuals. Historical clustering checkpoints and historical native inference caches were not used. New input checksums, frozen cell-selection manifests, computational settings, source identities and installed-package checksums accompany the results. Independent checks replayed BayesPrism regressions, full-family BH adjustment, prioritization membership, continuous-association calculations, precision–recall operating points, F1 and matched-cap optima. The independent-cohort generator reproduced its first seeded cohort exactly. A separately compiled CDSeq2 source archive reproduced a new donor fit exactly; this one-donor source check is distinct from the complete native rerun. The reconstruction record below describes source-level verification and the distinction from the earlier archived input version.

#### 1.6 Model development, comparative scope and performance boundaries

The six-feature specification was developed by removing the composition leave-one-out feature after earlier results from these evaluation donors had been examined. For the present reconstruction, the specification, feature definitions and selection policies were retained, and coefficients and smooths were refitted using reconstructed training components only. Training/evaluation donor separation does not undo the earlier model-development review, and reprocessing the same donors does not create an independent biological validation cohort. The separate 40-cohort simulation freezes the refitted scorer and primary rule before generating new outcomes; it assesses transfer under its defined generator rather than validation in new biological cohorts.

BayesPrism had higher scenario-average median expression recovery in six of seven cell types in the unselected expression benchmark. Similar excitatory-neuron recovery is descriptive and is not a formal equivalence result or a claim of equal accuracy across all types. Contribution ranking remained competitive and exceeded GAM6 in immune and vascular cells. In the direct-aggregation benchmark, selected medians were 0.399 for oligodendrocyte precursor cells, 0.390 for immune cells and 0.065 for vascular cells. Undefined correlations were omitted only from summaries; 399 selected immune-cell and 395 selected vascular-cell profiles had defined correlations. In the nine-scenario common-panel comparison, the prioritized vascular-cell median was 0.077. Relative enrichment therefore does not imply strong absolute recovery in every population.

The new generator uses training-derived distributions and assigned local haplotype windows rather than real gene cis annotations. It does not model long-range linkage disequilibrium, population structure or reference mismatch, and independent factors within each cell type simplify cross-type co-regulation. Five signal cohorts per setting provide limited Monte Carlo precision. The 200,000-read modeled-panel depth is below the original direct-aggregation libraries, and many profiles have at least one scorer feature outside its training range. These settings assess a defined transfer problem; they do not establish robustness across tissues or all sequencing depths. Broader method comparisons should also evaluate matched eligible gene sets, computational cost and inference diagnostics.

### 2 Mathematical identifiability of individual-level expression

#### 2.1 Multiple cell-type profiles can produce exactly the same bulk distribution

Let $\mathbf{\Phi}_i$ be the $G \times K$ matrix whose columns are the cell-type expression probabilities of individual $i$. Even if the RNA fractions $\boldsymbol{\rho}_i$ were known exactly, the bulk data inform only the mixture probability

$$\boldsymbol{p}_i = \mathbf{\Phi}_i \boldsymbol{\rho}_i, \qquad \boldsymbol{B}_i \sim \text{Multinomial}(L_i, \boldsymbol{p}_i). \tag{S1}$$

Each of the $K$ normalized expression profiles has $G-1$ free parameters, whereas the mixture has only $G-1$ independent probabilities. At an interior point with positive fractions, unrestricted individual expression therefore has $(K-1)(G-1)$ locally unidentifiable directions even when fractions are fixed. For $G = 2{,}000$ and $K = 7$, this is 11,994 directions per individual. Unknown fractions add further ambiguity.

A constructive example makes the issue explicit. Choose two types $a, b$ with $\rho_{ia}, \rho_{ib} > 0$ and any nonzero vector $\boldsymbol{u}_i$ satisfying $\mathbf{1}^\top \boldsymbol{u}_i = 0$. Set

$$\boldsymbol{\phi}'_{ai} = \boldsymbol{\phi}_{ai} + t_i \boldsymbol{u}_i, \tag{S2}$$

$$\boldsymbol{\phi}'_{bi} = \boldsymbol{\phi}_{bi} - t_i \frac{\rho_{ia}}{\rho_{ib}} \boldsymbol{u}_i. \tag{S3}$$

For sufficiently small $|t_i|$ around interior profiles, both new profiles remain nonnegative and normalized. Yet their changes cancel exactly:

$$\rho_{ia}\phi'_{ai} + \rho_{ib}\phi'_{bi} = \rho_{ia}\phi_{ai} + \rho_{ib}\phi_{bi}. \tag{S4}$$

Thus the two sets of cell-type expression profiles have the same bulk-data likelihood at every sequencing depth. With two genes and fractions $(0.6, 0.4)$, profiles $(0.8, 0.2)$ and $(0.2, 0.8)$ produce bulk probabilities $(0.56, 0.44)$. The distinct profiles $(0.7, 0.3)$ and $(0.35, 0.65)$ produce the same bulk probabilities. No estimator using only this mixture can uniquely distinguish these alternatives.

#### 2.2 Why accurate proportions and average profiles do not guarantee variation recovery

Estimating fractions with fixed, sufficiently distinct reference profiles is a different inverse problem: the unknowns are the $K$ mixture weights. Allowing all profiles to change independently by individual introduces many additional unknowns. Increasing the number of donors does not by itself remove this ambiguity, because each donor contributes another set of individual-specific profiles. Shared-profile assumptions or additional measurements can reduce the ambiguity, but their validity must be justified.

In the construction above, $t_i$ can vary across individuals, including with phenotype or genotype, without changing any bulk mixture. Therefore different cell-type-specific variation patterns, and different cell-type attributions of an association, can agree equally well with the observed bulk data. Good bulk reconstruction, accurate RNA fractions or accurate average reference expression do not alone establish accurate recovery of every gene/type trajectory.

#### 2.3 Regularization, sampling information and the role of the scorer

Reference-informed priors favor biologically plausible profiles and make estimation practical. They can yield a proper posterior and stable estimates while the bulk likelihood remains non-identifiable. The resulting allocation along ambiguous directions depends on reference information and modeling assumptions. Stronger priors reduce variance but can shrink genuine individual deviations toward the reference. Finite read depth, low type abundance, overlapping expression and reference mismatch further limit accuracy; when a type contributes no RNA, its expression is unobserved in that sample. Normalization also limits the analysis to relative expression over the modeled panel.

The reliability scorer addresses this challenge empirically by learning which gene/type profiles tend to preserve across-donor variation under the evaluated setting. It does not prove identifiability or certify any single profile as correct. Prioritization can enrich accurate estimates and concordant associations while reducing coverage, as shown in the main analyses. These mathematical limits explain why CDSeq2 can perform strongly for supported profiles, particularly in excitatory neurons, without recovering all underlying component expression or guaranteeing cell-type attribution for every association.

### 3 Detailed inference and application models

#### 3.1 Reference informed expression inference

Let $B_{gi}$ be the bulk count for gene $g = 1, \dots, G$ in individual $i = 1, \dots, N$, and let $L_i = \sum_g B_{gi}$. For cell type $k = 1, \dots, K$, $\rho_{ik}$ is the RNA fraction and $\boldsymbol{\phi}_{ki}$ is the individual-specific gene-expression probability vector. With normalized smoothed reference $\boldsymbol{\mu}_k$, the model is

$$\boldsymbol{\phi}_{ki} \sim \text{Dirichlet}(\tau_k \boldsymbol{\mu}_k), \qquad \boldsymbol{\rho}_i \sim \text{Dirichlet}(\boldsymbol{\alpha}), \tag{S5}$$

$$\boldsymbol{B}_i \mid \boldsymbol{\rho}_i, \boldsymbol{\phi}_i \sim \text{Multinomial}\left(L_i, \sum_{k=1}^{K} \rho_{ik} \boldsymbol{\phi}_{ki}\right). \tag{S6}$$

Here $\tau_k$ controls reference precision, $\sum_k \rho_{ik} = 1$, and $\sum_g \phi_{gki} = 1$. Conditional allocation of the observed reads follows

$$(A_{g1i}, \dots, A_{gKi}) \mid B_{gi}, \boldsymbol{\rho}_i, \boldsymbol{\phi}_i \sim \text{Multinomial}\left(B_{gi}, \left\{ \frac{\rho_{ik} \phi_{gki}}{\sum_{\ell=1}^{K} \rho_{i\ell} \phi_{g\ell i}} \right\}_{k=1}^{K} \right). \tag{S7}$$

Fractions describe RNA contributions over the modeled panel, rather than cell-number fractions. Reference priors regularize the underdetermined mixture without guaranteeing unique likelihood-based recovery.

The recorded settings were 300 MCMC iterations, retention of the last 100 allocation draws, reference precision 1,000,000, reference proportion 1, reference smoothing enabled with smoothing alpha 1, prior adjustment disabled and RNG setting 1. Each donor was fitted separately using one computational thread. Allocated counts were averaged across retained draws and checked to reconstruct each bulk gene. The returned estProp supplied RNA fractions. This configuration is the evaluated inference procedure; the saved allocation summaries do not establish multi-chain convergence.

Let $\widehat{A}_{gki}$ be the posterior mean allocated count and $C_{gki}$ the known component count. Relative expression and its across-donor recovery are

$$\widehat{q}_{gki} = \frac{\widehat{A}_{gki}}{\sum_{h=1}^{G} \widehat{A}_{hki}}, \qquad q_{gki} = \frac{C_{gki}}{\sum_{h=1}^{G} C_{hki}}, \qquad r_{gk} = \mathrm{cor}_i(\widehat{q}_{gki}, q_{gki}). \tag{S8}$$

Spearman recovery was calculated on the same profiles. The ratio of mean allocations need not equal the posterior mean expression probability. Undefined correlations were retained as missing rather than set to zero. Known RNA fractions were $\rho_{ik}^{\mathrm{true}} = \sum_g C_{gki}/L_i$; fraction accuracy was summarized by across-donor Pearson correlation, mean absolute error and root mean squared error. Expression RMSE was calculated across donors on the within-type CPM scale. The unselected two-method benchmark used the 1,969–1,971 genes retained by BayesPrism preprocessing in each scenario, without output reliability filtering, and compared both methods against identical expected and sampled components restricted to that panel. CDSeq2 estimates were restricted and renormalized for evaluation; its inference was not refitted. If $a_{ki}$ is the fraction of CDSeq2 allocated counts retained in type $k$, its RNA fractions were rebased as $\widehat{\rho}_{ik}^{R} = \widehat{\rho}_{ik} a_{ki} / \sum_{\ell} \widehat{\rho}_{i\ell} a_{\ell i}$. BayesPrism final fractions already refer to its retained panel. These common-panel accuracy calculations do not change the original CDSeq2 eQTL tests. Scenario summaries reuse the same 40 donors and are descriptive.

#### 3.2 Three-workflow expression recovery on the common benchmark

Supplementary Fig. S4C adds CDSeq2 (with gene-type prioritization) to the same nine-scenario, seven-cell-type benchmark used for standard BayesPrism and CDSeq2 (without gene-type prioritization). Each scenario uses the same retained genes and realized component truth for both methods. We used the archived scenario GAM6 scores, which were computed from the original 2,000-gene bulk panel and inferred fractions without genotype, association statistics or observed recovery as inputs. Scores were not refitted or recalculated after input-QC restriction. Instead, ranking was restricted to the common retained panel within each scenario and cell type. All common-panel profiles had finite scores and satisfied the archived common-rankable flag. We ranked by descending GAM6 score, broke ties by gene name, and selected $\lfloor 0.2M \rfloor$ of $M$ eligible profiles. Observed correlations and training-range flags did not affect selection. This is the top-20% expression-profile policy; the top-40% nominated eQTL policy uses a different universe and is unchanged.

The common panel contained 1,969 genes in one scenario, 1,970 in seven and 1,971 in one. Selection retained 393–394 profiles per scenario and type, or 3,545 of 17,730 pooled gene/scenario profiles per type (19.9944%, reported as 20.0%). Standard BayesPrism and CDSeq2 (without gene-type prioritization) each retained all 17,730 profiles. GAM6 selection did not change inferred expression, its common-panel normalization, the per-profile correlations in Table S14, or RNA-fraction results in Fig. S4A–B. In particular, expression was not renormalized over the selected subset.

For plotting and median calculation, undefined correlations were omitted after selection. Selected counts with defined correlations were 3,545 for astrocytes, excitatory neurons, inhibitory neurons, oligodendrocytes and oligodendrocyte precursor cells, 3,524 for immune cells and 3,459 for vascular cells. Table S24 reports the retained, defined and undefined counts for every workflow, scenario and cell type. Pooled median recovery for BayesPrism / CDSeq2 (without gene-type prioritization) / CDSeq2 (with gene-type prioritization) was 0.305 / 0.219 / 0.518 in astrocytes, 0.840 / 0.850 / 0.975 in excitatory neurons, 0.447 / 0.390 / 0.602 in inhibitory neurons, 0.384 / 0.363 / 0.732 in oligodendrocytes, 0.257 / 0.147 / 0.350 in oligodendrocyte precursor cells, 0.173 / 0.138 / 0.348 in immune cells, and 0.055 / 0.020 / 0.077 in vascular cells. These are descriptive pooled-profile summaries; Table S12 instead supports averages of scenario-specific medians. The CDSeq2 (with gene-type prioritization) set has a higher median than either complete set in every type, while covering approximately one fifth of profiles. This comparison quantifies the prioritization benefit and coverage cost; it is not an equal-coverage comparison or evidence that selection changes inference. Donors and genes recur across scenarios.

Table S23 records every common-panel score, rank, eligibility flag and selection decision. The accompanying scripts reproduce selection from the archived score files or the bundled score table, and plot correlations from the unchanged Table S14. Source hashes and an independent verification record accompany the outputs.

##### 3.3 Observable features and GAM6 training

Using the unsmoothed normalized pooled reference $\mu_{gk}$ and estimated fractions $\widehat{\rho}_{ik}$, define the reference-weighted contribution and bulk CPM as

$$w_{gki} = \frac{\mu_{gk}\widehat{\rho}_{ik}}{\max(\sum_{\ell=1}^{K} \mu_{g\ell}\widehat{\rho}_{i\ell}, 10^{-300})}, \qquad b_{gi} = 10^6 B_{gi}/L_i. \tag{S9}$$

The six gene/type features are

$$x_{1,gk} = N^{-1}\sum_i w_{gki}, \qquad x_{2,gk} = \log_2\left(N^{-1}\sum_i w_{gki}B_{gi} + 0.5\right), \tag{S10}$$

$$x_{3,gk} = \frac{\mu_{gk}}{\sum_\ell \mu_{g\ell}}, \qquad x_{4,gk} = \mathrm{sd}_i[\log_2(b_{gi} + 0.5)], \tag{S11}$$

$$x_{5,gk} = \frac{\max_i B_{gi}}{\max(\sum_i B_{gi}, 1)}, \qquad x_{6,gk} = \mathrm{cor}_i(b_{gi}, \widehat{\rho}_{ik}). \tag{S12}$$

These capture contribution, expected count, reference specificity, bulk variability, donor concentration and composition correlation. The last feature uses linear CPM.

Set $\widetilde{x}_{j,gk} = \log_2(x_{j,gk} + 10^{-6})$ for $j \in \{1, 3, 4, 5\}$ and $\widetilde{x}_{j,gk} = x_{j,gk}$ otherwise. The cell-type-specific regression and reported score are

$$z_{gk} = \mathrm{atanh}[\min\{0.995, \max(-0.995, r_{gk})\}], \tag{S13}$$

$$z_{gk} = \beta_{0k} + \sum_{j=1}^{6} f_{jk}(\widetilde{x}_{j,gk}) + \epsilon_{gk}, \qquad \epsilon_{gk} \sim \mathcal{N}(0, \sigma_k^2), \tag{S14}$$

$$\widehat{r}_{gk} = \tanh\left[\widehat{\beta}_{0k} + \sum_{j=1}^{6} \widehat{f}_{jk}(\widetilde{x}_{j,gk})\right], \qquad S_{gk} = 100\max(0, \widehat{r}_{gk}). \tag{S15}$$

Gaussian generalized additive models were fitted by restricted maximum likelihood (REML) with shrinkage cubic smooths of basis dimension five. Predictors with fewer than five distinct values used linear terms; constant predictors were omitted. Models required at least 100 distinct eligible genes. Feature values outside training ranges were flagged; the archived experiment retained these predictions.

Five gene folds were fixed before fitting the six-feature specification. The archived training eligibility and fold assignments were preserved. Thirty-five fold fits and seven final cell-type models were fitted. Gene-fold diagnostics complement donor-level evaluation. The reported results use this archived training protocol; the final six-feature specification was selected after evaluation inspection.

##### 3.4 Continuous phenotype construction and testing

For each source cell type, ten genes were sampled from the upper half of positive training expression variances to define each of three programs. Let $s_i$ be the standardized first principal-component score from the selected evaluation profiles. We generated

$$y_i = \sqrt{h}\, s_i + \sqrt{1-h}\, \varepsilon_i, \qquad \varepsilon_i \overset{\text{iid}}{\sim} \mathcal{N}(0,1), \qquad h \in \{0.25, 0.50, 0.75\}. \tag{S16}$$

Three repetitions per setting yielded 189 signal phenotypes. The parameter $h$ specifies the construction scale and need not equal the realized variance fraction. An additional 200 independent Gaussian phenotypes supplied global-null tests. This construction originally used all 40 evaluation donors. The earlier manuscript excluded original donor index 3 after inspection of `trait_031`. We retained that same exclusion for this rerun, without choosing a new donor from the reconstructed results. The remaining 39 phenotype values, original principal-component program scores and noise draws were retained; programs and simulated values were not regenerated. Expression estimates, within-donor normalization and all archived selection memberships were also retained. Thus $h$ describes the original simulation construction and is not a newly calibrated variance fraction in the reduced cohort. These results constitute a post hoc cohort restriction and association reanalysis.

For inferred expression $X_{gki} = \log_2(10^6 \widehat{q}_{gki} + 0.5)$, association testing used

$$X_{gki} = a_{gk} + b_{gk} y_i + e_{gki}, \qquad H_0 : b_{gk} = 0. \tag{S17}$$

The same analysis used $q_{gki}$ for observed components. With an intercept and no additional covariates, the two-sided slope test is equivalent to a Pearson correlation test, using 37 residual degrees of freedom for the 39-donor analysis. Benjamini–Hochberg adjusted values were calculated separately for inferred and observed expression over the complete gene/type family, with significance at $q_{\text{BH}} \leq 0.05$. Selection at retention fractions 0.1 through 1.0 did not recompute adjusted values. Same-sign overlap required significance in both analyses. Agreement divided this overlap by inferred calls; full-type reference recall divided it by all observed-component calls in the evaluated type. Ratios were averaged over programs and repetitions, excluding undefined ratios.

#### 3.5 Interpretation and donor sensitivity of the continuous-phenotype example

The identifier `trait_031` is an archive label for the first excitatory-neuron expression program, signal fraction $h = 0.50$ and noise repetition 1. It is a simulated quantitative phenotype, not a disease name or measured clinical variable. The phenotype combines a standardized principal-component score from ten observed-component program genes with independent Gaussian noise. Both panels of main Fig. 3 use 39 donors, excluding original donor index 3. The plotted phenotype retains the original 40-donor standardization so that slope units remain comparable to the archived full-cohort analysis.

In Fig. 3B, the horizontal coordinate is the association slope from observed excitatory-neuron components and the vertical coordinate is the slope from CDSeq2 estimates. Points below equality at positive horizontal values indicate underestimated positive associations; they do not define a separate biological group. In the reconstructed full-cohort result, donor 3 had phenotype $-3.729$ standard deviations on the 40-donor scale. Exclusion substantially reduces that pattern, particularly in the fixed GAM6-selected subset. Remaining off-diagonal points reflect imperfect preservation of phenotype-related expression variation.

Table S25 retains the reconstructed full-cohort result and all 40 single-donor omissions, using original benchmark donor indices. Index 0 denotes all 40 donors; index 3 corresponds to the 39-donor Figure 3. For each omission, we refitted both simple association regressions while retaining expression inference, within-type normalization, the 40-donor phenotype scale and frozen 400-gene GAM6 membership. Excluding donor 3 increased all-gene slope concordance from 0.513 to 0.787 and reduced the median absolute slope difference from 0.135 to 0.034. In the selected subset, concordance increased from 0.761 to 0.992 and median absolute slope difference decreased from 0.049 to 0.021. Table S16 supplies the 39-donor slopes and Table S17 supplies the 39 retained donors, with their original indices. The same historical exclusion was applied to all 389 newly generated continuous phenotypes. This remains a sensitivity analysis: retaining the earlier decision does not make its original choice prespecified or establish that donor 3 is erroneous. The saved data do not identify a unique technical cause for the expression discrepancy.

##### 3.6 Nominated variant simulation and evaluation

Candidates were stratified using baseline GAM6 quintiles in excitatory neurons, inhibitory neurons and oligodendrocytes. The design selected ten single-type targets per quintile/type, eight genes per type pair for each shared or opposite pattern, four other-type control genes per remaining type and 40 wholly null genes. Previously used pilot genes were excluded. Candidate support required nonzero expression in at least 80% of training and evaluation donors, bounded source log-CPM variability and a maximum source panel share of 0.02. These truth-based restrictions construct the benchmark; they are not information available to the operational scorer.

For nominated variant $v$, genotypes $G_{vi} \sim \text{Binomial}(2, 0.30)$ were drawn for the 40 evaluation donors subject to the archived genotype eligibility checks. Define $\widetilde{G}_{vi} = G_{vi} - \overline{G}_v$. For a targeted gene/type pair with baseline component $C^0_{gki}$, the expected count was perturbed as

$$C^*_{gki} = C^0_{gki} 2^{\gamma_{gk} \widetilde{G}_{vi}}. \tag{S18}$$

Writing $X^0_{gki} = \log_2(10^6 C^0_{gki}/L_{ki} + 0.5)$ and analogously $X^*_{gki}$, calibration solved

$$\frac{\left|\text{slope}_{G_v}(X^*_{gk} - X^0_{gk})\right| \text{sd}(G_v)}{\text{sd}(X^0_{gk})} = s, \qquad s \in \{0, 0.5, 0.8\}, \quad |\gamma_{gk}| \leq 8. \tag{S19}$$

The sign followed the injection design. Untested reserve genes were rescaled to preserve each component library $L_{ki}$, while non-target nominated genes were protected. Realized counts and bulk mixtures were generated by

$$\boldsymbol{C}_{ki} \sim \text{Multinomial}(L_{ki}, \boldsymbol{C}^*_{ki}/L_{ki}), \qquad B_{gi} = \sum_k C_{gki}. \tag{S20}$$

The signal ratio is a standardized slope increment, not a heritability or fixed log fold change. Three repetitions produced nine scenarios and 360 CDSeq2 donor fits. Association tests fitted $X_{gki} = a_{gk} + b_{gk}G_{vi} + e_{gki}$ for each nominated variant–gene/type pair. The archived design and calibration outputs specify each effect.

Association testing used the common nominated family of 1,778 gene/type tests, with BH adjustment before selection. Correct detections required a significant injected effect with the correct sign; false calls occurred in non-injected gene/type pairs, and wrong-sign target calls were recorded separately. Pooled FDP was false calls divided by all calls. Detection among selected injected tests and recall of all injected tests used different denominators. Scenario GAM6 ranking was computed from post-injection bulk data and inferred fractions without genotypes or association statistics. Baseline ranking and both full-panel and matched-tested-budget policies were retained in the supplementary results. CDSeq2 (with gene-type prioritization) eQTL results use the scenario GAM6 ranking and top-40% nominated-test budget; BayesPrism receives no output selection, as detailed below.

#### 3.7 Specified BayesPrism preprocessing and three-workflow comparison

BayesPrism 2.2.3 was run using its documented GEP interface with pooled raw counts from the original 20 reference donors and one state per broad type. Its cleanup.genes function excluded groups Rb, Mrp, other_Rb and chrM using the installed human gene annotations; exp.cells = 1 removed genes absent from every pooled reference profile. Nineteen genes were removed by category cleanup, and none by the zero-expression criterion. No explicit MALAT1 or sex-chromosome exclusion was applied. Optional marker selection was not used. The package's documentation recommends input cleanup and describes marker selection as optional (`https://github.com/Danko-Lab/BayesPrism`).

The tutorial example additionally includes MALAT1, chrX and chrY in the cleanup groups and uses exp.cells = 5 on individual cells (`https://github.com/Danko-Lab/BayesPrism/blob/main/tutorial_deconvolution.html`). The current pooled-GEP configuration does not implement that individual-cell prevalence filter; exp.cells = 1 on pooled profiles is a zero-reference-expression check. General guidance makes sex-chromosome exclusion conditional on reference–mixture sex differences. We therefore describe the comparator by its explicit settings and do not claim to have reproduced every tutorial filtering choice. Sensitivity to the broader cleanup configuration has not been evaluated.

The native new.prism settings outlier.cut = 0.01 and outlier.fraction = 0.1 filter genes exceeding 1% of bulk reads in more than 10% of samples. The function aligns the surviving mixture/reference genes and normalizes the reference using pseudo.min = $10^{-8}$. After cleanup and outlier filtering, 1,969–1,971 of the 2,000 genes remained in each scenario. Three nominated genes, MRPL48, PLP1 and RPL7, were excluded in every scenario; LRRC4C was additionally excluded in scenario 3 and NRG1 in scenario 9. This left 250–251 of 254 nominated genes (1,750–1,757 potentially available gene/type tests before checking for constant profiles). Table S19 records gene-level exclusions separately for cleanup and final retention. Preprocessing uses neither genotype, association statistics nor injection labels.

All nine cohorts were freshly fitted with no designated tumor key, update.gibbs = TRUE, chain length 1,000, burn-in 500, thinning two and alpha one. Seeds were $20261013 + j$ for scenario index $j = 1, \ldots, 9$. These are explicitly recorded inference controls, rather than a claim that every package setting was left at its default. Expression was extracted with get.exp (initial cell-type posterior allocated counts) and normalized to within-type $\log_2(\mathrm{CPM} + 0.5)$ over the retained panel. Final RNA fractions were extracted with get.fraction for the accuracy benchmark. Allocated counts were checked to conserve the retained bulk counts.

The eQTL comparison uses the original nominated family of 1,778 tests. Missing or constant-expression tests enter BH adjustment as $p = 1$, then have their reported $p$ and $q$ marked unavailable and receive no call. Thus preprocessing does not reduce the multiplicity denominator. BayesPrism reports every available test, without a gene/type ranking or retention threshold. CDSeq2's original association statistics are unchanged; its scenario GAM6 selection retains 101 of 254 nominated excitatory-neuron pairs per repetition, ranked by descending score with gene-name tie breaking. The CDSeq2 (with gene-type prioritization) results retain their original BH-adjusted values. No CDSeq2 scorer or benchmark-defined support filter is applied to BayesPrism.

The three displayed workflows are standard BayesPrism with the specified preprocessing, CDSeq2 (without gene-type prioritization) and CDSeq2 (with gene-type prioritization). The first two compare inference without output reliability selection. The latter two share inferred expression and original association statistics, isolating the added prioritization step and its effect on reporting coverage. Correct-discovery counts and recall against all injected targets accompany FDP to expose the cost of selection and preprocessing. Raw fraction and expression accuracy are evaluated on a common retained panel as described above. The framework comparison establishes empirical enrichment in the CDSeq2 (with gene-type prioritization) set; it does not imply that selection changes the underlying estimates, that every selected profile is accurate, or that one inference method is uniformly superior.

##### 3.8 Precision–recall, F1 and matched empirical FDP caps

We evaluated standard BayesPrism after the specified input preprocessing, CDSeq2 (without gene-type prioritization) and CDSeq2 (with gene-type prioritization). GAM6 ranks use the original common finite-score eligibility set for the GAM6, contribution and expected-count rankings, followed by descending GAM6 score and gene-name tie breaking. The selected count is $\lfloor 0.4M \rfloor$ for $M$ eligible nominated pairs within a scenario and cell type (101 of 254 in excitatory neurons). These CDSeq2 eligibility features do not filter or rank BayesPrism output. Membership is independent of genotype, association statistics and injection labels and remains unchanged throughout the threshold sweep.

Let $\mathcal{U}$ be the original nominated testing universe and $\mathcal{T} \subseteq \mathcal{U}$ the injected gene/type targets, with $N_+ = |\mathcal{T}|$. For workflow $w$, let $A_w$ be its available and retained tests. Using the original BH-adjusted values, the call set at threshold $\tau$ is

$$\mathcal{C}_w(\tau) = \{j \in A_w : q_{wj} \leq \tau\}. \tag{S21}$$

Writing $d_j \in \{-1, 1\}$ for the injected sign and $\widehat{b}_{wj}$ for the inferred slope, direction-aware counts are

$$\mathrm{TP}_w(\tau) = \sum_{j \in \mathcal{C}_w(\tau)} \mathbf{1}\{j \in \mathcal{T},\ \mathrm{sign}(\widehat{b}_{wj}) = d_j\}, \tag{S22}$$

$$\mathrm{FP}_w(\tau) = |\mathcal{C}_w(\tau)| - \mathrm{TP}_w(\tau), \qquad \mathrm{FN}_w(\tau) = N_+ - \mathrm{TP}_w(\tau). \tag{S23}$$

Thus a wrong-sign call at an injected target contributes both an incorrect reported effect and a missed correct effect. False positives comprise non-injected calls plus wrong-sign target calls. This definition differs from the earlier non-injected-FDP summaries, which record wrong-sign calls separately; for the pooled excitatory-neuron results at $q \le 0.05$ there are no wrong-sign calls, so both FDP definitions coincide. Both components are retained in Tables S20–S22.

$$\mathrm{Precision} = \frac{\mathrm{TP}}{\mathrm{TP} + \mathrm{FP}}, \qquad \mathrm{Recall} = \frac{\mathrm{TP}}{N_+}, \tag{S24}$$

$$F_1 = \frac{2\mathrm{TP}}{2\mathrm{TP} + \mathrm{FP} + \mathrm{FN}}, \qquad \mathrm{FDP}_{\mathrm{dir}} = \frac{\mathrm{FP}}{\max(|\mathcal{C}|, 1)}. \tag{S25}$$

With no calls, precision is undefined, directional FDP is defined as zero, and recall and F1 are zero when injected targets exist. Targets discarded by GAM6, removed during BayesPrism input preprocessing or unavailable because of constant expression remain in $N_+$. Recall is never renormalized to the selected target set. This measures recovery of the original injected effects, rather than a method-specific subset.

Precision–recall curves enumerate every distinct finite original $q$ value, up to one; all ties enter simultaneously. The no-call state is included in the tables and omitted from the curves because its precision is undefined. We do not interpolate operating points or recompute BH adjustment after selection. Each scenario retains its original 1,778-test BH family. For the main curves, counts are pooled across the three repetitions at each nonzero signal ratio and the same $q$ cutoff is applied to all three repetitions. These are pooled count ratios rather than averages of repetition-specific ratios; $N_+ = 246$ for excitatory neurons. Supplementary Fig. S10 and the tables provide each repetition separately. F1 is reported at the original $q \le 0.05$ operating point, without selecting a threshold to maximize F1.

For an empirical directional FDP cap $\alpha$, the matched-cap comparison selects an attainable threshold according to

$$\tau_w^*(\alpha) \in \underset{\tau:\,\mathrm{FDP}_{\mathrm{dir},w}(\tau)\le\alpha}{\arg\max} \mathrm{TP}_w(\tau). \tag{S26}$$

Among thresholds with the same maximum true-positive count, we choose fewer false positives and then the strictest threshold. The no-call state is always feasible. Every observed cutoff is considered because empirical FDP need not be monotone in the cutoff. A common cap need not yield exactly equal achieved FDP; discrete calls and ties can leave a workflow below the cap. Main Fig. 5 shows 5%, 10% and 20% caps, and Table S22 includes caps from zero to 50% in one-percentage-point increments plus 100%. The table records each selected threshold, achieved FDP, call count, precision, recall, F1 and availability denominator.

This threshold choice uses known injection labels and is an oracle, retrospective summary of attainable performance. It is not an operational filtering rule, a prospective threshold-calibration method or evidence of FDR control. The GAM6 retention fraction remains fixed at 40% and is not optimized against injection truth. In particular, the threshold for CDSeq2 (with gene-type prioritization) at ratio 0.5 under the 20% cap is approximately 0.0838; the matched-cap analysis is therefore distinct from the fixed $q \le 0.05$ comparison. Repetitions share the same 40 biological donors, so the curves are descriptive and no independent-cohort confidence interval or significance claim is implied. The evaluated unit is a nominated variant–gene/type effect, not a genome-wide independent locus.

#### 3.9 Continuous-phenotype agreement, coverage and null results

For excitatory-neuron programs at signal fraction 0.50, mean agreement among CDSeq2 calls increased from 86.7% across all profiles to 91.5% in the GAM6 top 20% (Supplementary Fig. S7). This enrichment came with lower coverage: mean full-type reference recall declined from 77.2% to 15.4%. At signal fraction 0.75, the corresponding agreement increased from 75.4% to 78.5%, while full-type reference recall declined from 77.0% to 16.2%. These values average three programs and three noise repetitions and are not estimates from independent biological cohorts.

The comparison with contribution and expected-count ranking shows that selection should be assessed jointly by agreement and coverage, rather than by agreement alone. Results across all source cell types and signal fractions are provided in Supplementary Fig. S2 and Tables S3 and S9. Observed-component significance is a noisy reference and does not identify a set of known causal phenotype effects; disagreement with that reference is therefore not equivalent to a false biological association.

In 200 independent Gaussian null phenotypes, any rejection across the full family occurred in 2.5% of CDSeq2 analyses and 3.0% of observed-component analyses. Mean numbers of rejections were 65.915 and 0.030, respectively. Thus a low proportion of null traits with any call can coexist with a large mean number of calls in the inferred-expression analysis; the two summaries answer different questions and should both be reported. These results describe global-null behavior conditional on the fixed expression matrices; they do not establish error control for cell-type attribution under structured genetic effects. The accompanying continuous-null replicate table records individual-trait call counts.

At the same 20% retention budget, GAM6 achieved agreement/full-type reference recall of 91.5%/15.4% at signal fraction 0.50, compared with 89.2%/9.5% for contribution ranking and 83.6%/9.5% for expected-count ranking. At signal fraction 0.75, the corresponding values were 78.5%/16.2%, 74.2%/11.6% and 72.5%/10.2%. These same-budget comparisons support the ranking's utility for prioritizing a subset; they do not imply that restricting to 20% improves comprehensive discovery coverage (Tables S3 and S9).

#### 3.10 Nominated-variant attribution, coverage and matched-error comparisons

Cell-type attribution remained imperfect. There were six false calls among 45 selected calls at signal ratio 0.5 and 33 among 108 at ratio 0.8, corresponding to pooled false discovery proportions of 13.3% and 30.6%. All selected false calls at both strengths arose from genes perturbed in other cell types. This pattern illustrates the distinction between detecting a tissue-level genetic signal and assigning it to the correct cellular source (Supplementary Table S5). Nominal BH significance did not ensure 5% empirical cell-type attribution error in these simulations.

At the original $q_{\mathrm{BH}} \leq 0.05$ threshold and signal ratio 0.5, precision was 79.0% for BayesPrism, 79.5% for CDSeq2 (without gene-type prioritization) and 86.7% for CDSeq2 (with gene-type prioritization).

Full-target recall was 44.3%, 37.8% and 15.9%, respectively, with F1 scores of 0.568, 0.512 and 0.268 (Fig. 5C). At ratio 0.8, precision was 66.4%, 69.8% and 69.4%, recall was 69.1%, 69.5% and 30.5%, and F1 was 0.677, 0.697 and 0.424 in the same workflow order (Fig. 5D). Adding GAM6 improved precision at ratio 0.5 but not at ratio 0.8; its loss of recall lowered F1 at both strengths. The scorer targets expression recoverability, not eQTL F1, and discarded profiles can contain detectable true effects.

Each cap is a shared upper bound, rather than a requirement for exactly equal achieved FDP: at the 20% cap, CDSeq2 (with gene-type prioritization) and BayesPrism achieved 16.4% and 19.3% FDP at ratio 0.5, and 16.4% and 19.5% at ratio 0.8. Thresholds were chosen retrospectively using simulation truth to maximize correct discoveries under each cap. This describes attainable performance and does not provide a deployable threshold or establish FDR control. All-type fixed-threshold results and repetition-specific curves are in Supplementary Figs. S9 and S10; Tables S20–S22 provide every threshold, denominator and achieved FDP.

At a common 5% directional FDP cap, prioritization increased CDSeq2 correct-discovery counts from six to 25 at signal ratio 0.5 and from nine to 26 at ratio 0.8; BayesPrism recovered 22 and 26. At the 10% cap, CDSeq2 recovered six and 15 without selection and 26 and 38 with selection, compared with 49 and 36 for BayesPrism. At the 20% cap, the prioritized workflow recovered fewer effects: 46 and 56, compared with 93 and 88 for unprioritized CDSeq2 and 109 and 107 for BayesPrism. Thus the prioritization advantage is strongest for stringent error budgets in this benchmark and does not extend to every cap. BayesPrism receives only its specified input preprocessing and association threshold, without gene/type output ranking. Complete outcomes for all cell types remain in Figures S9 and S11 and Tables S5, S20–S22.

### 4 Frozen-rule, LD-aware simulation benchmark

#### 4.1 Design freeze and independence

Before generating any new test outcomes, we recorded the scorer identity, settings, seeds, source-file checksums, package versions, primary retention (40%) and significance threshold ($q \leq 0.05$). The six signal settings each have five independent simulation realizations, and ten additional cohorts have no injected effects. Original evaluation-donor profiles never enter the generator. GAM6 was refitted on the reconstructed training components, then frozen before LD-aware cohort generation. Its form was chosen after reviewing earlier evaluation results, but the new simulated outcomes were not used to select its form or retention rule. There is no calibration split because no parameter is tuned to the new outcomes. Synthetic cohorts are conditionally independent given the training-derived distribution parameters and the fixed haplotype library. Reusing a reference library does not turn them into independent biological validation cohorts.

The modeled panel has 2,000 genes and seven cell types. We sampled 200 target genes uniformly, with seed 2026101401, from genes with mean training-component CPM between 1 and 10,000 in each of excitatory neurons, inhibitory neurons and oligodendrocytes. Neither scores nor expression-recovery accuracy entered gene sampling. Each target receives one of 40 LD windows, with five genes per window. The fixed family has 1,400 gene/type hypotheses. Sample sizes are 60 and 120, and nominal genetic variance fractions are 0.10 and 0.30. Additional settings use mean excitatory-neuron RNA fraction 0.20 or a 1.5-fold multiplier on log-expression heterogeneity, both at $n = 120$ and fraction 0.30. No post hoc donor exclusion is applied. The design comprises 3,600 independently generated synthetic individuals across 40 cohorts (Table S26).

#### 4.2 Haplotype sampling and local variant coverage

We used the publicly accessible phased phase-3 chromosome-22 VCF on GRCh37:

`https://ftp.1000genomes.ebi.ac.uk/vol1/ftp/release/20130502/ALL.chr22.phase3_shapeit2_mvncall_integrated_v5b.20130502.genotypes.vcf.gz`.

The accompanying population panel identifies 503 European-reference individuals (1,006 phased haplotypes). Candidate windows start at position 20,000,000 and then every 50,000 bases, each spanning 25,000 bases. We retained the first 40 among 80 candidate windows with at least five PASS, biallelic, fully phased SNPs having reference minor allele frequency at least 0.05. Window eligibility uses only reference genotypes, not association outcomes. For each synthetic individual and window, two whole haplotypes are sampled with replacement and summed. This preserves empirical within-window haplotype structure in expectation while generating new genotype combinations. Windows are sampled independently, so long-range LD and population structure are not modeled. Test-cohort minor allele frequency must be at least 0.05; all eligible SNPs in the assigned window enter the test. Window coordinates, SNP counts and LD summaries are provided in Table S27. The block field indexes the 40 retained windows, while candidate_window records the original candidate-window number before sparse windows were omitted.

The random assignment of windows to genes is a controlled simulation device. It does not represent the genes' true cis locations or a whole-genome cis scan, and the new benchmark makes no causal fine-mapping claim. A discovery is a correct gene/type discovery when a direct regulatory effect was injected in that pair, regardless of whether the most significant SNP is causal or tags it through LD. This endpoint differs from the correct-sign nominated-variant endpoint of the earlier experiment; their numerical FDP values should not be interpreted as interchangeable.

#### 4.3 Independent expression and count generation

Let $Y_{gki}$ be training-donor $\log_2(\mathrm{CPM}+0.5)$ for gene $g$, cell type $k$ and donor $i$. We estimate a mean $m_{gk}$ and obtain three loading columns $L_k$ from the singular value decomposition of centered training expression. The loading scale is singular value divided by $\sqrt{39}$. Residual standard deviations satisfy

$$\sigma_{gk}^2 = \max\left(0.15^2, \widehat{\mathrm{Var}}(Y_{gk}) - \sum_{r=1}^{3} L_{gkr}^2\right).$$

For each new individual, independent factor scores $F_{kir} \sim N(0,1)$ and residuals $\epsilon_{gki} \sim N(0,1)$ produce

$$\eta_{gki} = m_{gk} + a\left(\sum_{r=1}^{3} L_{gkr}F_{kir} + \sigma_{gk}\epsilon_{gki}\right) + \beta_{gk}Z_{gki},$$

where $a = 1$ or 1.5 and $Z_{gki}$ is the sample-standardized dosage of the causal SNP chosen for that gene/type pair. For excitatory neurons, inhibitory neurons and oligodendrocytes, 40 distinct target genes are independently sampled for injection; all other pairs have $\beta_{gk} = 0$. The coefficient has an independently sampled sign and magnitude

$$|\beta_{gk}| = a\sqrt{\sum_{r=1}^{3} L_{gkr}^2 + \sigma_{gk}^2}\sqrt{\frac{h^2}{1-h^2}}, \qquad h^2 \in \{0.10, 0.30\}.$$

Thus $h^2$ is a nominal pre-normalization genetic variance fraction under the generator, not the realized association $R^2$ after normalization, count sampling and deconvolution. It is distinct from the earlier simulation's standardized effect ratio of 0.5 or 0.8. All injected coefficients, causal variants and test-cohort allele frequencies are archived in Table S35.

Within-type expression probabilities are $P_{gki} = 2^{\eta_{gki}} / \sum_{g'} 2^{\eta_{g'ki}}$. RNA fractions follow a Dirichlet distribution with concentration 35 and a mean derived from training-component RNA fractions, with a minimum mean of 0.005 followed by renormalization. The typical excitatory-neuron mean is approximately 0.610. In the low excitatory-neuron setting, the excitatory-neuron mean is set to 0.20 and other means are rescaled proportionally to sum to 0.80. Each individual has 200,000 reads over the modeled panel: type totals are multinomial draws from its RNA fractions, and gene counts within each type are multinomial draws from $P_{gki}$. Summing the realized type components gives bulk counts. The modest-depth setting is deliberately distinct from the original high-depth directly aggregated data. The scorer's out-of-training-range flags are retained and reported; a high flag rate limits interpretation of its extrapolation. Normalization can propagate small relative-expression changes, so the endpoint is recovery of direct injected regulatory targets, not the complete set of nonzero marginal effects on normalized expression. Consequently, direct-target FDP under non-null settings is an attribution metric and need not equal the false-discovery proportion for the statistical null of zero marginal association. Under the complete global null, no genetic effect is injected anywhere, allowing a separate check of the association pipeline's nominal calibration.

#### 4.4 Native fitting and fixed association procedure

Every cohort is fitted anew. CDSeq2 uses 300 MCMC iterations, the last 100 draws, reference precision $10^6$, pooled reference counts, aggregate smoothing with pseudocount one, and no prior adjustment. Expression is the posterior mean allocated count, identical to the estimator used in the earlier benchmark. The pooled reference has one count profile per cell type and preserves the original reference aggregate; the original individual reference nuclei are not reintroduced as test individuals. The six-feature scorer is applied to observable features from the new bulk and estimated RNA fractions. Predictions use the archived model's transformations and retain finite out-of-range values with explicit flags.

BayesPrism 2.2.3 uses the same reference aggregate, the GEP interface, Rb/Mrp/other_Rb/chrM cleanup, exp.cells = 1, default outlier.cut = 0.01 and outlier.fraction = 0.1, and pseudo.min = $10^{-8}$. It has no tumor key, marker restriction or output reliability filter. Gibbs sampling uses 1,000 iterations, burn-in 500, thinning two and alpha one, with reference updating enabled. Both methods conserve the observed bulk counts over their respective modeled panels. Gene preprocessing is reported for every cohort in Table S33. Each method uses its own retained panel for within-type CPM normalization, as in the earlier application comparison; missing target genes remain in the full multiplicity and recall denominators.

For every eligible SNP $v$ in the window $\mathcal{V}_g$, ordinary least squares fits $\log_2(\mathrm{CPM}_{gki} + 0.5) = b_{0,gkv} + b_{1,gkv}G_{vi} + e_{gkvi}$, with an intercept and no additional covariates. Genotypes are generated independently of expression noise and composition, so this experiment does not test confounding adjustment. Gene-level evidence is

$$p_{gk}^{\mathrm{gene}} = \min\left\{1, |\mathcal{V}_g| \min_{v\in\mathcal{V}_g} p_{gkv}\right\}, \qquad q_{gk} = \mathrm{BH}_{1400}(p_{gk}^{\mathrm{gene}}).$$

The within-gene Bonferroni rule accommodates arbitrary LD but can be conservative. Missing or constant profiles receive $p = 1$ during adjustment and never make a call. Only CDSeq2 (with gene-type prioritization) then restricts reporting to the top 80 of 200 pairs per cell type, using descending score and gene-name tie breaking. Full-family $q$ values remain unchanged. The fixed primary threshold is 0.05; nominal thresholds 0.01 and 0.10 and retentions 20%, 60% and 80% are prespecified sensitivities, not optimized replacements. Per-test evidence, selection, truth labels and lead variants are in Table S28. Score, rank and range fields are populated only for CDSeq2; they do not select BayesPrism or observed-component results. Selected-candidate counts describe output membership before availability checks, while the availability flags and preprocessing tables document which tests can be evaluated.

##### 4.5 Accuracy, uncertainty and null assessment

For each replicate, false calls are called gene/type pairs without a direct injected effect. We calculate $\mathrm{FDP} = \mathrm{FP}/\max(1, \mathrm{TP} + \mathrm{FP})$, full-target recall $\mathrm{TP}/N_{\text{injected}}$, and $F_1 = 2\mathrm{TP}/(\mathrm{TP} + \mathrm{FP} + N_{\text{injected}})$. Undefined precision for zero calls is left undefined, while FDP is zero. The mean of the five independent replicate FDPs estimates the expected direct-target attribution FDP under each signal setting; it is not automatically the FDR for zero-marginal-association tests. Monte Carlo standard errors use the between-replicate standard deviation divided by $\sqrt{5}$; five replicates give limited precision and do not establish universal error control. Pooled precision and counts are additionally reported but are not called mean FDR. The complete fixed-threshold results are in Table S29.

Precision–recall curves sweep distinct unadjusted gene-level $p$ thresholds, retaining all ties and fixed prioritization membership (Table S30). Within one cohort these give the same ordering as global BH values. They are descriptive curves; the predefined $q = 0.05$ operating point, rather than a truth-optimized point on the curve, supplies the primary result. A secondary, explicitly retrospective diagnostic chooses the attainable point with maximum true positives under each truth-measured FDP cap of 5%, 10% and 20% (Table S34). These cap-based thresholds do not constitute a deployable FDR procedure.

Ten independently generated 60-individual no-effect expression cohorts at typical cell-type abundance are each deconvolved once. For each cohort, 100 independent whole-haplotype genotype realizations are then tested using unchanged inferred expression and scorer ranks. Under the global null, FDP equals the indicator of any rejection. All 1,000 genotype/cohort results are retained (Table S31), but uncertainty is assessed across the ten independent expression cohorts, not by treating the 1,000 tests as independent expression datasets. This global-null analysis cannot establish cell-type attribution control when true effects in other cell types can be misassigned. No null or signal result is used to retune the scorer, retained proportion, preprocessing or significance threshold. Secondary expression-recovery results on the same target universe are in Table S32: Pearson correlations of within-type CPM match the scorer-training target, and log-CPM correlations describe the association scale. Undefined correlations and selection flags are retained.

#### 4.6 What the additional benchmark establishes

The new experiment separates three questions. Expression-recovery comparisons evaluate whether the frozen scorer enriches profiles that preserve known across-individual expression variation. Direct-target association metrics assess recovery of injected gene/type effects, allowing a tagged local variant to support discovery without requiring causal fine mapping. Global-null testing evaluates nominal association calibration when no genetic effects are present anywhere. Better expression recovery, lower direct-target FDP and global-null calibration are related but distinct properties; none is substituted for the others.

The primary rule was specified before evaluation. Secondary retention curves and truth-matched caps are descriptive comparisons, not alternative rules selected to optimize the reported result. Reduced full-family null rejection after filtering can partly reflect the smaller reporting set. Evidence that the ranking prioritizes more accurately recovered expression comes instead from the selected-versus-complete recovery distributions (Fig. S14). Five signal cohorts per setting limit Monte Carlo precision, and the repeated gene panel and reference constrain the scope of generalization.

#### 4.7 Abundance sensitivity and null calibration

At $n = 120$, $h^2 = 0.30$ under typical abundance, prioritization increased pooled excitatory-neuron precision from 77.6% to 92.6%, while full-target recall decreased from 38.0% to 25.0%. Mean replicate FDP decreased from 23.1% to 9.3%, and mean F1 from 0.506 to 0.388; BayesPrism's mean F1 was 0.492. A higher selected-set precision therefore does not imply better overall F1. The fixed top-40% rule retained 50 of the 76 injected discoveries made by unprioritized CDSeq2 (Table S29).

In the higher-heterogeneity setting, prioritization increased pooled excitatory-neuron precision from 76.2% to 85.9%, with recall decreasing from 49.5% to 27.5% and mean F1 from 0.598 to 0.406. Mean direct-target FDP was 13.9% after selection, compared with 23.7% without selection (Fig. 6E). At the weaker fraction $h^2 = 0.10$, no workflow recovered an injected excitatory-neuron target at $n = 60$; at $n = 120$, recall was 2.0% for each unprioritized method and 1.5% after prioritization. Selection therefore does not compensate for limited association power. At $n = 60$, $h^2 = 0.30$, pooled precision was already high (92.0% for BayesPrism and 91.7% for unprioritized CDSeq2) and did not improve with selection (90.9%). All-type results and prespecified retention sensitivities are reported in Figures S13 and S15, with full precision–recall curves in Figure S12.

Cell-type abundance materially affected the result. In the low excitatory-neuron setting ($n = 120$, $h^2 = 0.30$, mean excitatory-neuron RNA fraction 0.20 rather than the typical 0.610), pooled excitatory-neuron discovery counts were 36 injected and 43 non-injected calls for BayesPrism, 36 and 42 for CDSeq2 (without gene-type prioritization), and 15 and 15 for CDSeq2 (with gene-type prioritization). Pooled precision increased from 46.2% to 50.0% with selection, while full-target recall decreased from 18.0% to 7.5%. Mean replicate direct-target FDP remained 53.9% after selection. Median CPM-profile recovery was 0.242, 0.239 and 0.371 for the three workflows, respectively. Relative enrichment therefore coexisted with weak absolute recovery. Selection retained exactly 400 of 1,000 target/cohort profiles in every signal setting; BayesPrism had 990 defined correlations after preprocessing, compared with 1,000 for unselected CDSeq2. Across the six settings, 77.3–93.9% of unselected CDSeq2 excitatory-neuron target profiles and 72.5–95.0% of selected profiles had at least one feature outside its training range (Tables S28 and S32). These flags identify extrapolation; they were not used to discard profiles or retune the frozen rule.

The independent global-null assessment separated prioritization from nominal error control. At nominal $q \leq 0.05$, the probability of any rejection across all seven cell types was 27.7% for BayesPrism, 24.1% for CDSeq2 (without gene-type prioritization) and 10.5% for CDSeq2 (with gene-type prioritization); between-expression-cohort Monte Carlo standard errors were 2.54, 2.08 and 0.83 percentage points, respectively (main Fig. 6F; Table S31). Corresponding excitatory-neuron-specific probabilities, while retaining global multiplicity adjustment, were 2.6%, 2.2% and 0.9%. Observed components also showed full-family inflation (38.2%), demonstrating that the failure cannot be attributed solely to deconvolution. An independent regression audit replayed 135 nominal 5% calls across 100 genotype realizations in null cohort C07_R10, with all stored decisions agreeing. These findings support reporting calibration by cell type and show that restricting reported profiles can reduce null discoveries; they do not establish a 5% FDR guarantee for the prioritized workflow or isolate the scorer's benefit from the effect of reporting fewer targets. Sparse, heterogeneous profiles and the fixed parametric association model remain limitations in this modest-depth simulation.

#### ROSMAP reconstruction, versioning and source-level verification

The present results use a newly frozen reconstruction of the ROSMAP benchmark, not the historical prepared-input version. The source files are nine processed Seurat objects and deidentified participant metadata downloaded from `https://compbio.mit.edu/ad_aging_brain/`. The original clustering and primary-preparation workflow is available at `https://github.com/kkang7/CDSeq2/tree/main/reproduction/brain/preprocessing`. Frozen manifests identify the 2,000 genes and disjoint 40 training, 20 reference and 40 evaluation donors. Training/reference preparation selects the top-ranked donor-shared cluster at resolution 0.01 in each source component, using qualifying donor coverage, total nuclei, median qualifying donor count and cluster identifier. Reference sampling uses the recorded seed and cell caps. Evaluation counts aggregate all source nuclei for the evaluation donors, without the training/reference cluster filter or count resampling.

The source-data reconstruction ran on an Apple arm64 workstation with 128 GiB RAM, using R 4.6.1, Seurat 5.5.1, SeuratObject 5.4.0, Harmony 2.0.5 and Matrix 1.7-6. All nine source RDS files (approximately 174 GiB) matched the recorded download identities. Legacy Seurat object updates were checked to preserve raw counts and cell metadata. All nine clustering components completed and the four analysis inputs were assembled. New training-nucleus and ordered reference-nucleus manifests freeze this version's selections, so users can reconstruct counts without rerunning version-sensitive clustering.

An initial comparison with the historical input version reproduced all 560,000 evaluation counts and all 280 evaluation nucleus counts exactly, but found differences in 132,289 of 560,000 training counts and 164 of 280 training nucleus counts. The new reference contained 39,533 nuclei rather than 39,692; 29,108 nuclei were shared and all 58,216,000 shared-nucleus count entries agreed. Different reference membership changed 11,929 of 14,000 pooled reference entries. These differences concern upstream cell selection, not a claim that particular inferred genes were incorrect. The historical clustering environment was not fully documented, and dependency-version effects were not established as the cause. Historical identity checks were retained rather than changed to accept the new inputs.

The reconstructed inputs were registered as a separate benchmark version before native fitting.

A second, independent source-only aggregation used the newly frozen nucleus manifests and the original source RDS files, without reading any clustering checkpoints. It exactly reproduced all 560,000 training counts, all 560,000 evaluation counts, the 280 nucleus counts in each cohort, and every selected reference count block in its recorded order. Input MD5/SHA-256 identities, selection manifests, source checks and software versions are supplied in `reconstruction_and_verification/`. The source-level check establishes exact regeneration of this version's numerical inputs; it does not claim exact recovery of the historical cell selection or guarantee identical future clustering under different dependencies.

Every downstream analysis and both inference methods were then rerun from the reconstructed inputs. The six-feature specification, retention rules, association thresholds and random-seed policies were retained; model coefficients, scenario scores and native estimates were freshly computed. The 39-donor continuous sensitivity rule preserves the earlier donor-index-3 exclusion; all other benchmarks retain their specified cohorts. Expression programs and nominated gene lists were regenerated using the same selection algorithms and seed policies with the new scores and inputs; their identities are therefore not assumed to match the earlier simulation. New training-derived distribution parameters and a refitted scorer were frozen before generating the forty independent LD-aware cohorts. The primary figures and Tables S1–S35 report the reconstructed-input run; Tables S36–S42 provide the additional prioritization and stability diagnostics. The earlier manuscript and its numerical tables remain a separate archive.

The complete pipeline passed count-conservation, donor-separation, checksum, genotype, selection and statistical checks. LD-reference QC metadata were regenerated directly from the frozen haplotypes after an export-stage missing-file error; this did not change any simulation or fit. All newly computed caches and their input identities remain in the run directory. The accompanying compact independent-cohort bundle supports fresh native reruns, while the source and cell-selection manifests support regeneration of the brain inputs. Raw source files are not redistributed here.

### 5 Additional prioritization and null-stability diagnostics

#### 5.1 Score groups, controls and uncertainty

These diagnostics were formulated after reviewing the primary results. They neither retrain the scorer nor choose a new primary retention rule. All 30 signal cohorts contribute cell-type-specific score groups and equal-budget controls. Ten score groups are constructed within each cohort and type, with group 1 highest. Expression summaries average cohort medians; precision pools correct and false calls. Each group contains 20 candidates per cohort. The same genes can recur across cohorts and are never treated as independent biological replicates.

The controls retain 80 of 200 target pairs per type: the frozen GAM6 score, reference-weighted contribution, expected read support and 100 repeated random subsets. All share the original inferred expression, association statistics, testing family and $q \leq 0.05$ threshold. Scores and support features are computed without association outcomes. Random draws use seed 20261020. Cohort-bootstrap intervals use 2,000 draws, seeds 20261021 for controls and 20261022 for score groups. Methods are paired within cohort; random results are first averaged within cohort. Percentile intervals describe conditional simulation uncertainty with only five independent cohorts per setting. Undefined precision is not replaced by zero, and decile tables report defined bootstrap counts.

In the principal excitatory-neuron setting, GAM6, contribution, expected-count and random selection have pooled precision 92.6%, 92.7%, 88.9% and 77.5%, respectively. Corresponding correct-discovery counts are 50, 51, 48 and 30.8 (random average). Mean cohort median expression recovery is 0.844, 0.872, 0.838 and 0.702; these averages differ from medians pooled over all profiles. The paired GAM6-minus-contribution precision difference is $-0.13$ percentage points (95% interval $-1.11$ to 0.73), and the expression-recovery difference favors contribution ranking. In the lower-abundance setting, contribution precision is 64.3% versus 50.0% for GAM6. Thus learned prioritization is useful relative to random retention and the complete output in supported settings, but this audit does not establish uniform superiority over simple support ranking. Main Figure 7 and Figures S16–S17 report the additional comparisons; Tables S36–S38 and S41 provide all settings and cell types.

#### 5.2 Continuous-null rejection bursts and donor influence

Independent replay had already verified the regression and adjustment calculations. We therefore examined expression structure and sensitivity rather than treating the call bursts as a software arithmetic error. Null phenotypes 10, 55 and 112 produced 4,011, 4,702 and 4,467 calls in the 39-donor primary analysis. Across their called profiles, 96.4% had the greatest centered-expression contribution from original donor index 38, and the median maximum donor share of centered sum of squares was 89.6%. This donor is distinct from the earlier index-3 omission.

The first principal component of row-centered, unit-norm inferred log-CPM profiles explained 38.2% of profile variation, compared with 14.4% for observed components. The median absolute profile alignment with this component was 0.956 among calls from the three bursts. Those null phenotypes correlated with the component by 0.452, 0.474 and −0.465, respectively. The dominant profile pattern and chance phenotype alignment explain why many tests can reject together, without identifying a biological association. This is a conditional diagnostic association, not proof that one particular inference step caused the pattern.

Donor 38 has very low inferred modeled-panel library support in several types: 4.80 allocated reads for inhibitory neurons, 15.16 for oligodendrocyte precursor cells and 14.09 for vascular cells, with 1,744, 1,536 and 1,508 zero inferred genes out of 2,000, respectively. Observed component libraries for those types are 2,054, 32,166 and 3,048 reads. Such sparse inferred profiles, followed by log transformation with an offset, provide a plausible source of donor-dominated shared patterns. Original counts and zeros are tabulated in Table S42.

Omitting donor 38 additionally eliminates all calls for these three phenotypes in the resulting 38-donor diagnostic. This is not used to redefine the study cohort. The full 40-donor log-CPM analysis still has mean 55.655 calls per null phenotype, versus 65.915 with 39 donors. On the same 39 donors, CPM gives mean 0.080 calls while log-counts give 159.700. Transformations change the response scale and interpretation; these observations do not validate choosing the most favorable scale after viewing outcomes. The original log-CPM analysis is retained.

Any rejection occurs in 2.5% of the 200 primary null phenotypes. Under a complete null this is the empirical probability of a nonempty false discovery set, so the rare large bursts alone do not establish excess global-null FDR. They do identify sensitivity to sparse profiles and influential donors. The independent LD benchmark is a separate concern: full-family null rejection remains 10.5% after prioritization at a nominal 5% threshold. Neither the diagnostic donor omission nor a normalization change is promoted as a validated correction. Figure S18 and Tables S39–S40 and S42 report all sensitivities; primary outcomes remain unchanged.

The version 0.1.0 code and bundled reproduction materials are archived at `https://doi.org/10.5281/zenodo.23489824`. The tagged source commit is `1fa8e88f61b6da2dd58669f6ae12a5a9d41564ee`. Automated Linux and macOS checks cover package build/check, the synthetic tutorial and scorer replay; they do not claim a fresh full ROSMAP or LD benchmark run on those CI hosts.

### 6 Supplementary table guide

All tables are supplied as separate comma-separated values (CSV) files in Additional files 2–6: Tables S1–S11 in Additional file 2; S12–S19 in Additional file 3; S20–S25 in Additional file 4; S26–S35 in Additional file 5; and S36–S42 in Additional file 6. Original table filenames and numerical contents are preserved. The separate additional-file index maps every table to its archive.

S1. Gene/type features, observed recovery, predictions and range diagnostics.

S2. Training out-of-fold and evaluation retention curves for GAM6 and support rankings.

S3. Continuous-phenotype summaries across source types, strengths, evaluated types and retention, using 39 donors after excluding original donor index 3.

S4. Conditional global-null summaries for inferred and observed components, using the same 39-donor subset.

S5. Nominated eQTL detection, recall and error summaries across policies.

S6. Pooled excitatory-neuron eQTL results for the three workflows, including available tests, FDP, correct discoveries and full-target recall.

S7. Source-to-recipient counts by genetic scenario and selection policy.

S8. Per-test genetic association results for the compared engines.

S9. Continuous phenotype results before program and repetition averaging, using 39 donors after excluding original donor index 3.

S10. Injected gene/type design and effect labels.

S11. All-profile and GAM6 top-20% evaluation medians.

S12. Ground-truth fraction and expression accuracy, by cell type, engine, scenario and expected/observed component definition. Direct-aggregation results are included separately.

S13. Paired true and estimated RNA fractions; donors use within-benchmark indices.

S14. Per-gene across-donor expression correlations and CPM RMSE for the ground-truth comparisons.

S15. Same-sign eQTL agreement with observed components, for all nominated tests, by scenario, cell type and engine.

S16. Observed-component and CDSeq2 association slopes for all excitatory-neuron genes in trait_031, refitted on 39 donors with frozen top-20% selection.

S17. Donor-level phenotype and expression values for the JPT1 example; the 39 retained donors preserve their original benchmark indices, excluding index 3.

S18. Nominated excitatory-neuron eQTL tests and availability, adjusted values and gene-type prioritization membership; BayesPrism has no output selection.

S19. Native BayesPrism preprocessing availability for all 2,000 candidate genes in each scenario, including cleanup, final retention and nominated-gene status.

S20. Exhaustive precision–recall operating points for the three workflows, both nonzero signal ratios, each cell type and all types pooled. Repetition zero denotes counts pooled over repetitions 1–3; each repetition is also provided. Includes original $q$ cutoffs, directional and non-injected error counts, full-target denominators and available/retained test counts.

S21. Precision, full-target recall, F1 and directional FDP at the original $q \leq 0.05$ threshold, with counts and coverage denominators for every workflow, signal ratio, repetition and cell type.

S22. Maximum attainable recall under each empirical directional FDP cap, with the selected original $q$ threshold, achieved FDP and complete counts. Caps are upper bounds, not exactly equal achieved FDP. Thresholds are chosen retrospectively using known injection truth.

S23. Common-panel scenario GAM6 scores, range and eligibility flags, within-scenario/type ranks, candidate counts and top-20% membership used in Fig. S4C. Selection precedes omission of undefined recovery correlations.

S24. Three-workflow expression-recovery counts and summaries for Fig. S4C, by scenario and pooled (ALL): common-panel and score-eligible counts, retained proportion, retained/defined/undefined counts, median and quartiles.

S25. Figure 3 slope concordance and errors for all genes and the fixed GAM6 subset. Index 0 denotes all 40 original donors; indices 1–40 identify the omitted donor. Index 3 matches Figure 3 and 39-donor Table S17.

S26. Independent LD-aware cohort manifest, seeds, sample sizes, nominal genetic variance fractions, abundance and heterogeneity settings.

S27. Public haplotype-window coordinates, eligible SNP counts, allele frequencies and LD summaries.

S28. New-cohort gene/type association evidence, full-family adjusted values, availability, truth, scorer ranks and range flags for both methods and observed components.

S29. New-cohort fixed-threshold metrics and coverage, by replicate, workflow, cell type, nominal threshold and retention. Mean replicate FDP summarizes direct-target attribution error; pooled precision is a separate quantity.

S30. New-cohort precision–recall operating points with fixed membership and all threshold ties retained.

S31. Global-null results for 100 genotype realizations within each of ten independent null-expression cohorts. Expression-cohort clustering must be retained for uncertainty estimates.

S32. New-cohort target-expression recovery on both CPM and log-CPM scales, selection flags and scorer range diagnostics.

S33. Native BayesPrism cleanup and final gene availability for every new cohort.

S34. Secondary retrospective matched-FDP operating points for the LD-aware benchmark; thresholds use test truth and are not prospective rules.

S35. New-cohort injection design, effect coefficients, causal variants, allele frequencies and assigned LD windows for each target gene/type pair.

S36. Per-cohort score-group recovery, discovery counts and precision.

S37. Equal-budget ranking controls, including all random draws.

S38. Paired cohort-bootstrap uncertainty for equal-budget controls.

S39. Null phenotype, expression-structure and donor-omission diagnostics.

S40. Per-profile diagnostics for the three large null rejection bursts.

S41. Score-group summaries, counts and cohort-bootstrap intervals.

S42. Per-donor modeled expression support and zero-gene counts.

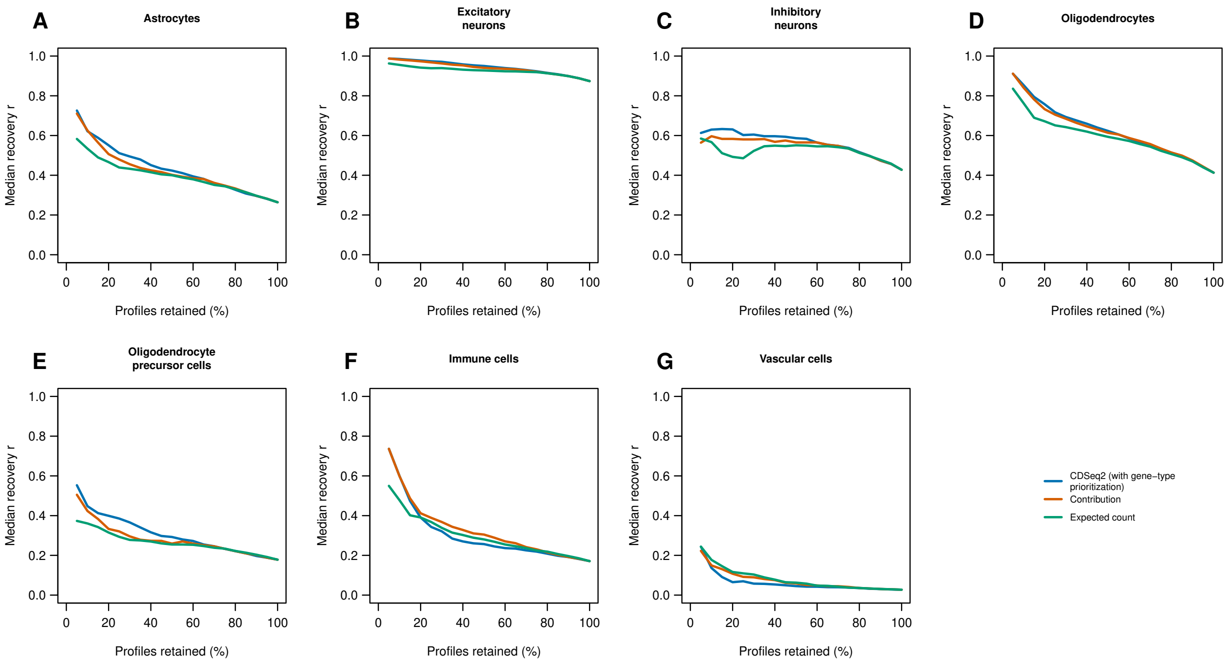


**Figure S1: Recovery curves in all seven cell types.** Median Pearson recovery for each selection method across retention fractions, using the same 40 evaluation donors and common eligible scoring universe as the six-feature scorer evaluation. Poor recovery in vascular profiles persists under restrictive selection. Undefined correlations are omitted. No confidence intervals are implied by the smooth connections between retention points. Visualization code was prepared with OpenAI Codex assistance. Underlying data are provided in Supplementary Table S2.

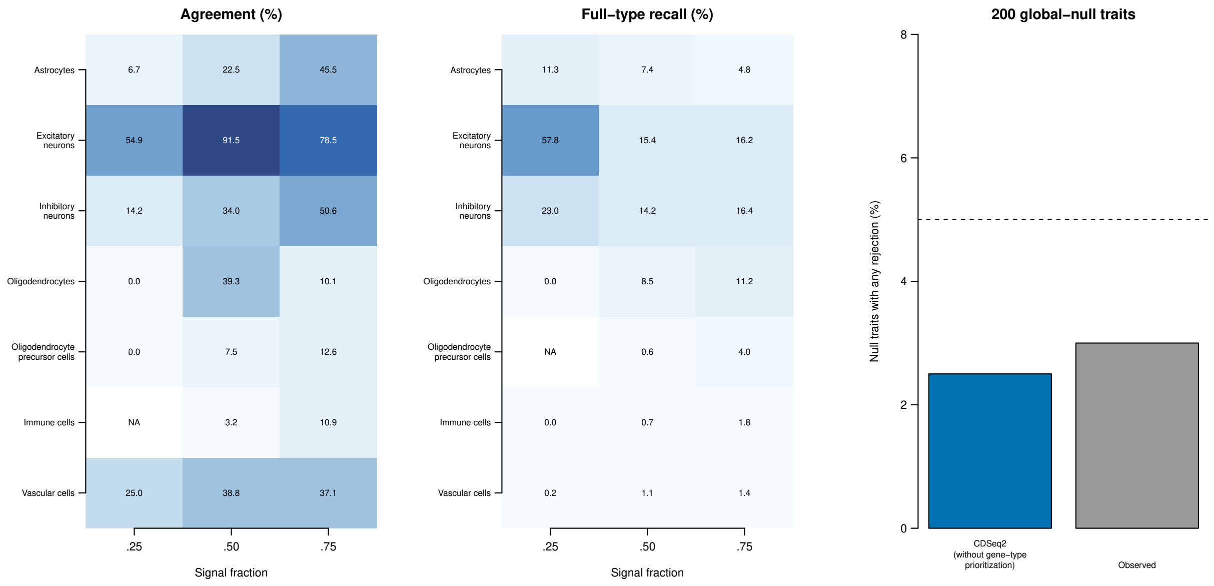


**Figure S2: Continuous associations and global nulls.** All panels use the same 39 donors after post hoc exclusion of original donor index 3, retaining the original phenotype values and frozen rankings. The two matrices summarize GAM6 top-20% selection when the phenotype source and evaluated cell type coincide. Values are mean agreement among calls and mean full-type reference recall, expressed as percentages over three programs and three noise repetitions. NA denotes an undefined ratio rather than zero performance. The right panel gives the proportion of 200 independent Gaussian null traits with any rejection in the complete testing family. The dashed line marks 5%; these conditional global-null results do not imply cell-type attribution control under alternatives. Visualization code was prepared with OpenAI Codex assistance. Underlying data are provided in Supplementary Tables S3 and S4.

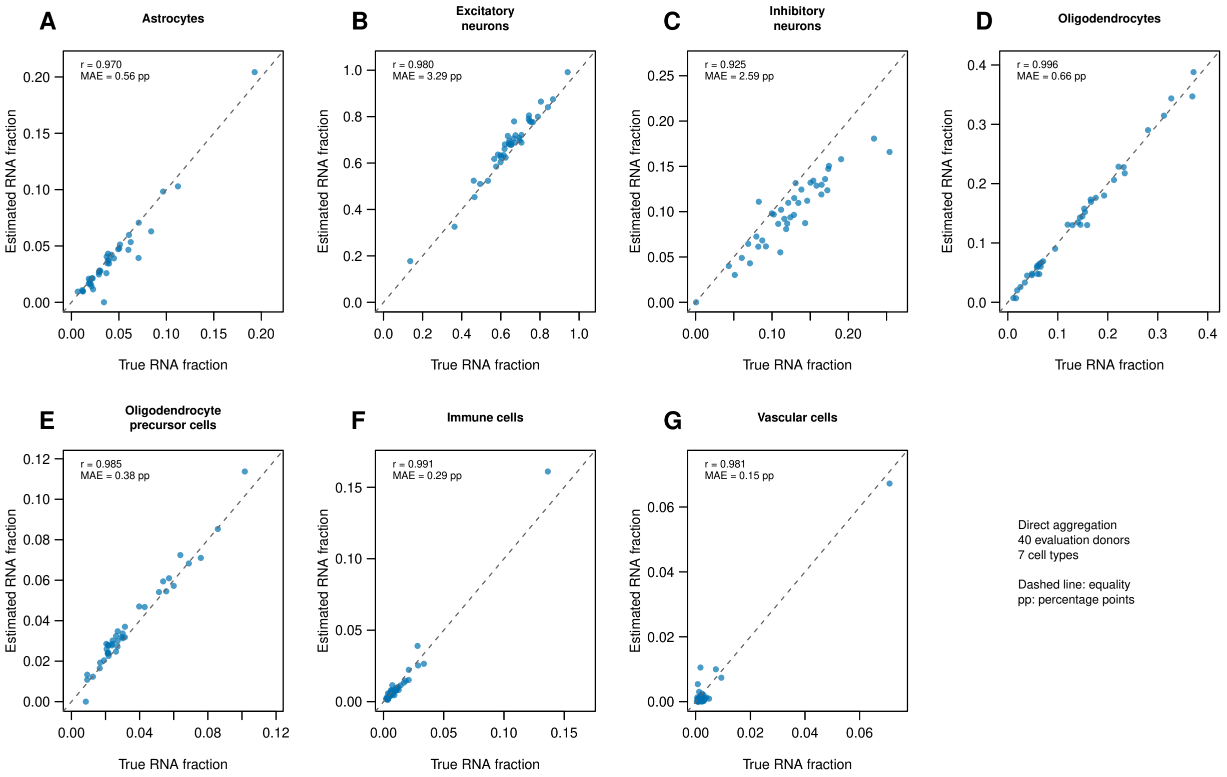


**Figure S3: Direct ground-truth RNA-fraction recovery.** Each panel shows one cell type; each point is one of 40 evaluation donors. Both axes are fractions of RNA over the 2,000-gene panel. The dashed line marks equality, and annotations give across-donor Pearson correlation and mean absolute error in percentage points. Ground truth is the component library divided by the bulk library. These are direct-aggregation data, with no multinomial resampling. Individual-expression recovery on these donors is shown in main Fig. 2; unfiltered metrics are in Table S12. Visualization code was prepared with OpenAI Codex assistance. Underlying data are provided in Supplementary Tables S12 and S13.

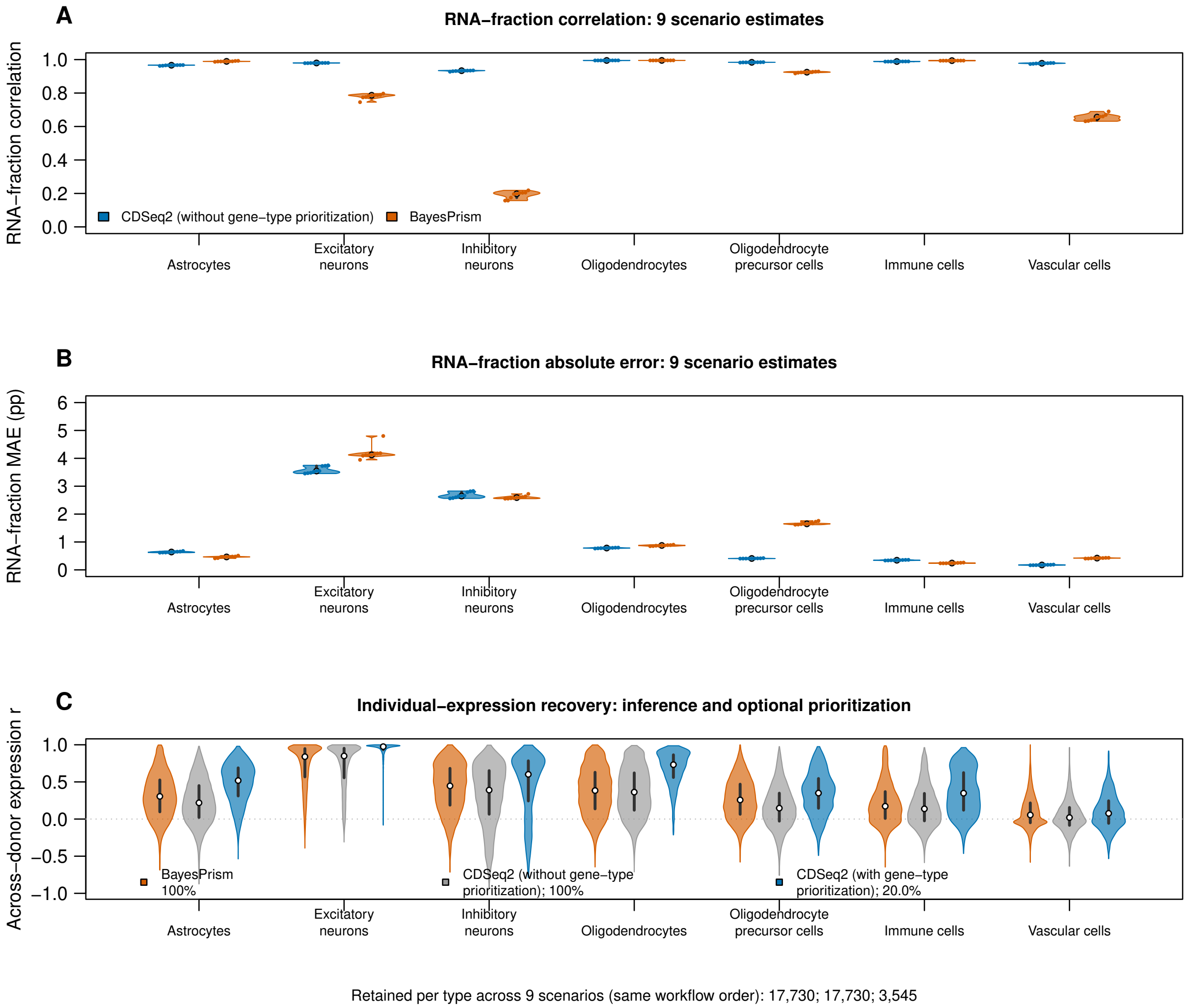


**Figure S4: Ground-truth recovery and the added value of GAM6 prioritization.** All panels use the same nine simulated scenarios and realized component truth on the common gene panel retained by BayesPrism input preprocessing. CDSeq2 estimates and truth are restricted to that panel; CDSeq2 RNA fractions are rebased using inferred expression. (A) RNA-fraction correlations and (B) mean absolute errors in percentage points compare CDSeq2 (without gene-type prioritization) (blue) and BayesPrism (orange); each violin contains nine scenario values, also shown as points. (C) Expression-recovery violins compare standard BayesPrism (orange), CDSeq2 (without gene-type prioritization) (gray) and CDSeq2 (with gene-type prioritization) (blue). Both unselected workflows retain 17,730 gene/scenario profiles per type. GAM6 retains 393–394 of 1,969–1,971 common-panel profiles per scenario/type, or 3,545 pooled profiles per type (20.0%; exact proportion 19.9944%). Selection ranks archived scenario scores, with gene-name tie breaking, without using recovery truth. Expression and its common-panel normalization remain unchanged. Undefined correlations are omitted only from plotting: selected plotted counts are 3,545 for astrocytes, excitatory neurons, inhibitory neurons, oligodendrocytes and oligodendrocyte precursor cells, 3,524 for immune cells and 3,459 for vascular cells. Table S24 provides all workflow-specific counts and medians. White dots mark medians; dark segments mark interquartile ranges. Equal maximum violin widths show within-group density, not sample size. The same 40 donors and genes recur across scenarios. Higher correlation and lower error indicate better accuracy; selected distributions describe a smaller reporting subset. Visualization code was prepared with OpenAI Codex assistance. Underlying data are provided in Supplementary Tables S12, S14, S23 and S24.

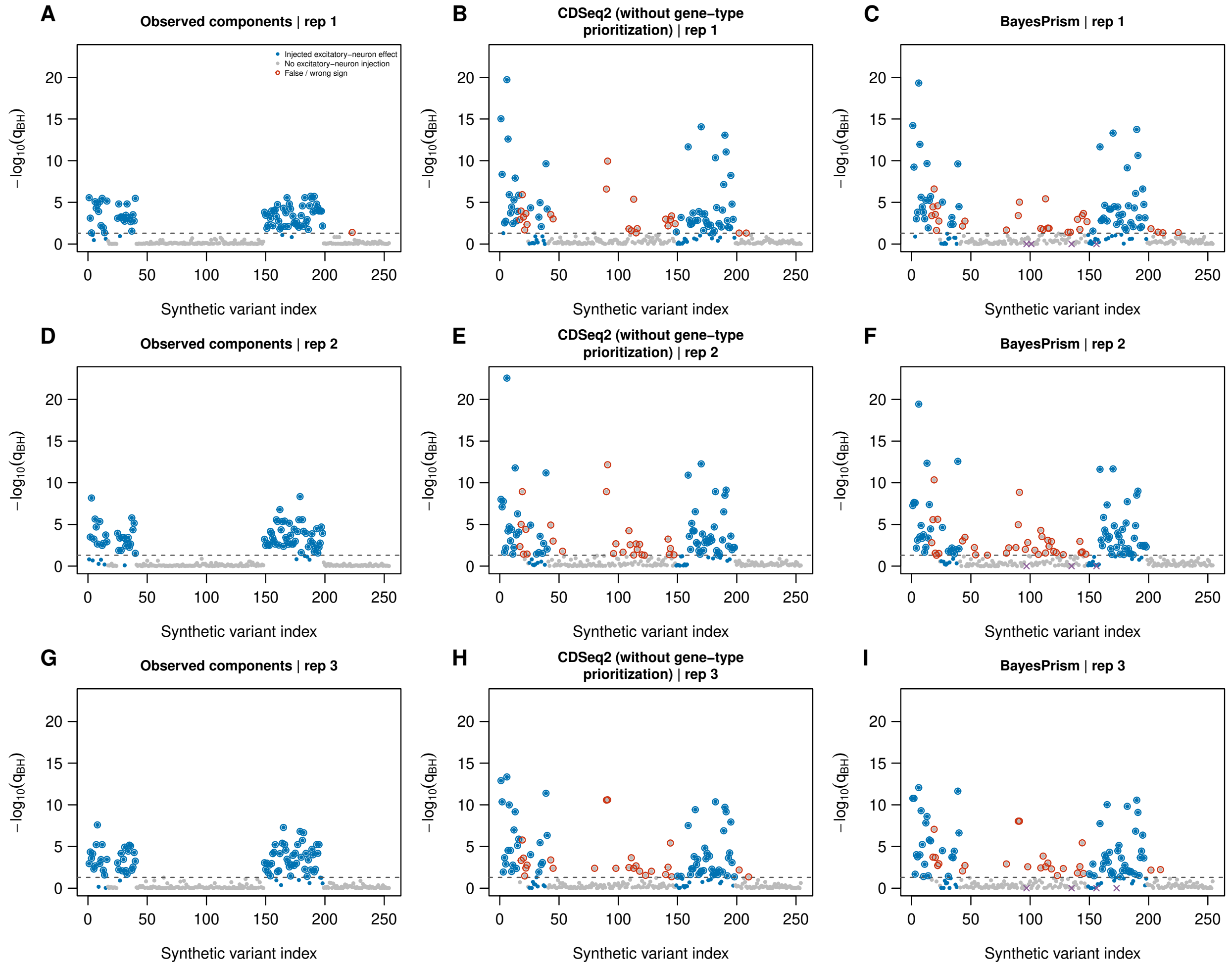


**Figure S5: Aligned Manhattan-style eQTL profiles at signal ratio 0.8.** Columns show observed components, CDSeq2 (without gene-type prioritization) and BayesPrism; rows show all three repetitions. Each point is one of 254 nominated excitatory-neuron variant–gene tests, ordered by synthetic variant identifier identically in every panel. The vertical axis is $-\log_{10}(q_{\text{BH}})$, with adjustment over all 1,778 gene/type tests before filtering. The dashed line is $q_{\text{BH}} = 0.05$. Blue points have an injected excitatory-neuron effect and gray points do not; circles around significant points denote calls with the injected sign (blue) or false/wrong-sign calls (red). All available tests are displayed without output reliability selection. BayesPrism uses native input preprocessing; purple crosses at baseline mark unavailable tests, not evidence of nonsignificance. Shared peaks show agreement in association evidence; differences show method-specific detection or attribution. Synthetic variants have no chromosome coordinates, so these are indexed simulation profiles, not genome-wide Manhattan plots. Visualization code was prepared with OpenAI Codex assistance. Underlying data are provided in Supplementary Table S8.

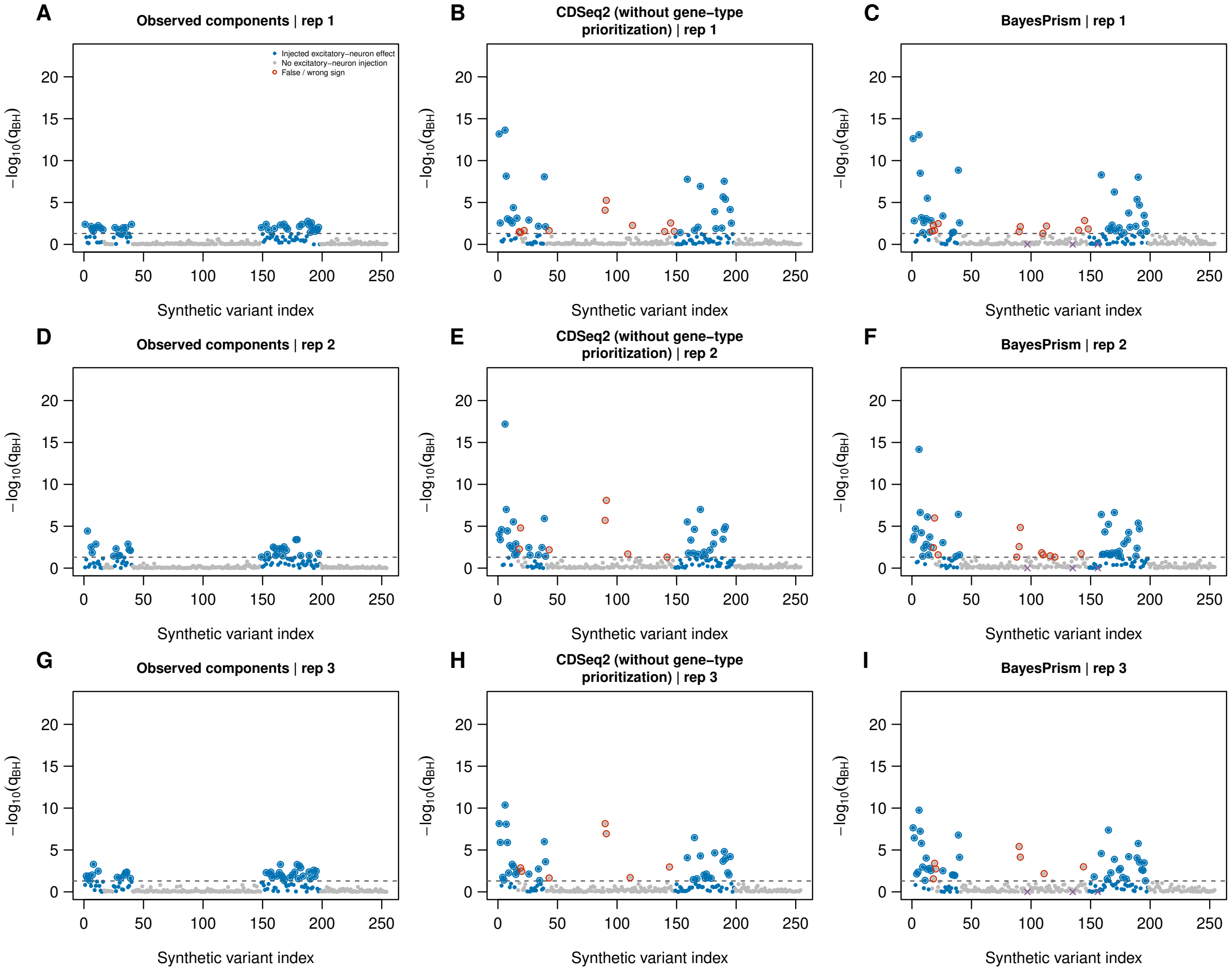


**Figure S6: Aligned Manhattan-style eQTL profiles at signal ratio 0.5.** The layout, test order, colors and thresholds match Supplementary Fig. S5. All three repetitions are shown at the moderate signal strength, using the same vertical scale as Fig. S5. Together, the two figures display agreement and missed or additional signals across both strengths without selecting a favorable repetition. Visualization code was prepared with OpenAI Codex assistance. Underlying data are provided in Supplementary Table S8.

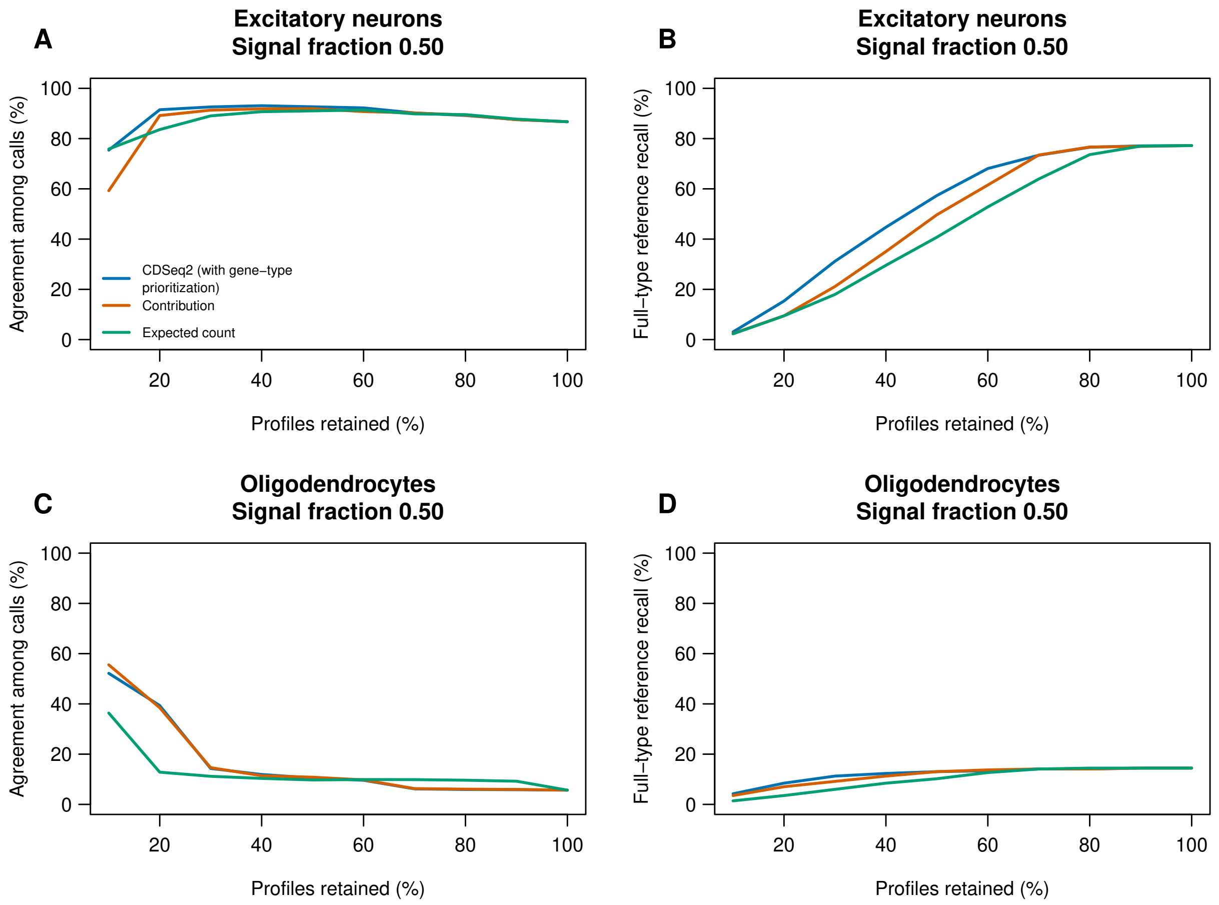


**Figure S7: Continuous-phenotype association agreement and coverage.** All associations were recomputed on 39 donors after post hoc exclusion of original donor index 3; original phenotype values and frozen rankings were retained. Rows show phenotypes generated from excitatory neurons (A,B) or oligodendrocytes (C,D) expression programs at signal fraction 0.50. The horizontal axis is the percentage of gene/type profiles retained; moving left applies more restrictive selection. Blue, orange and green curves rank profiles by GAM6, reference-weighted contribution and expected read support. (A,C) Agreement: among significant CDSeq2 associations retained, the percentage also significant in the observed-component analysis with the same sign. Higher values indicate greater concordance among reported candidates. (B,D) Coverage: that same concordant count divided by all significant observed-component associations in the corresponding cell type before selection. Higher values indicate that more reference associations are retained. Reading the panels together shows the agreement–coverage tradeoff: prioritization can enrich concordant candidates while discarding some associations. Each point averages nine program/noise combinations; undefined ratios are excluded. BH adjustment over the full gene/type family precedes selection and uses a 5% threshold. The observed-component analysis is a noisy reference, not causal truth. Visualization code was prepared with OpenAI Codex assistance. Underlying data are provided in Supplementary Tables S3 and S9.

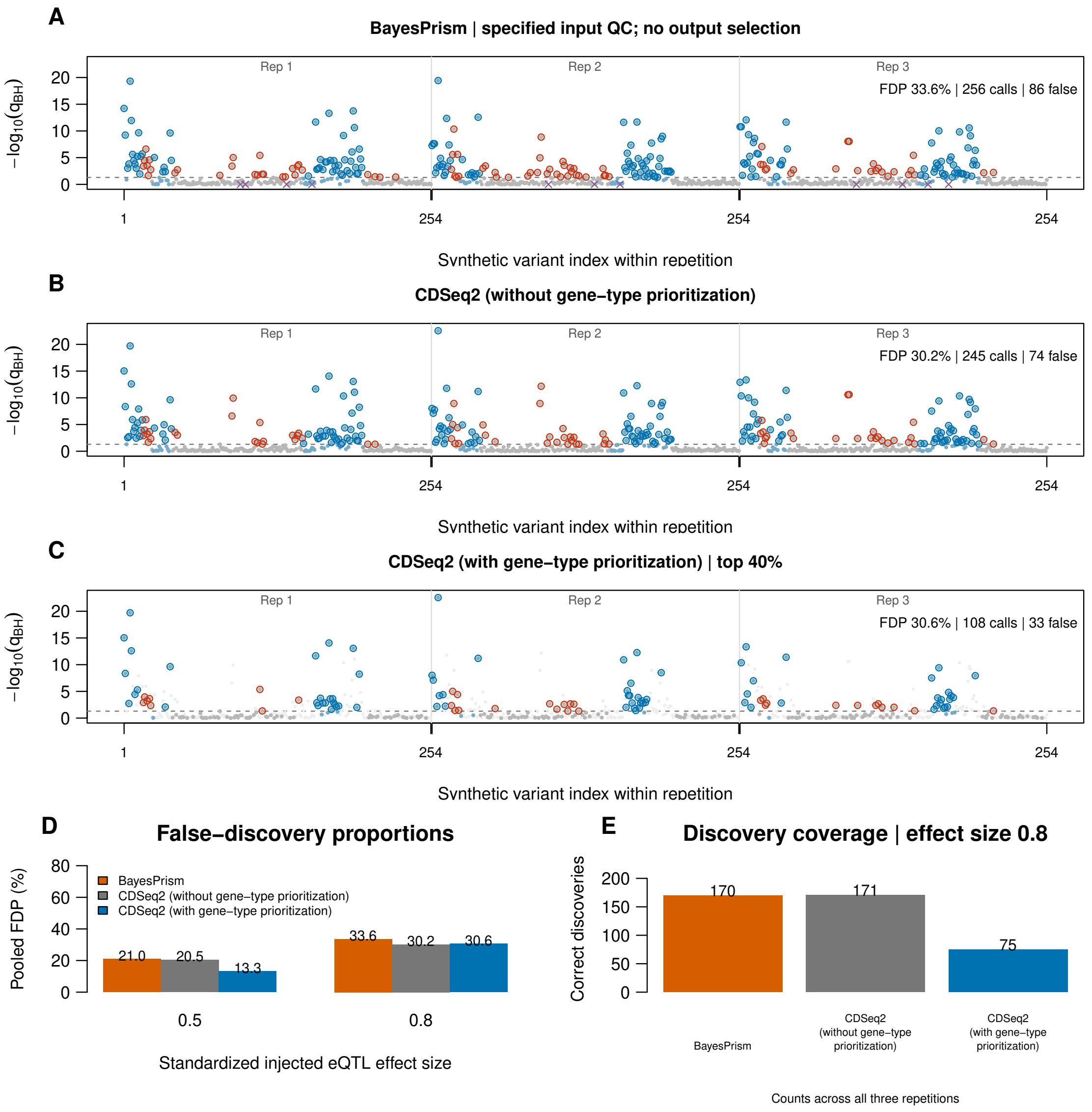


**Figure S8: Three-workflow eQTL comparison at signal ratio 0.8.** Panels A–C follow main Fig. 4: standard BayesPrism with the specified input preprocessing, CDSeq2 (without gene-type prioritization) and CDSeq2 (with gene-type prioritization) top-40% selection, each across all three repetitions. Only C applies output reliability selection. Purple baseline crosses in A mark preprocessing-excluded tests; discarded tests in C are faint background points. Panel D summarizes pooled FDP at both strengths; panel E shows correct-discovery counts at 0.8. CDSeq2 (with gene-type prioritization) retains the same inferred expression and BH-adjusted values as CDSeq2 (without gene-type prioritization). BayesPrism BH adjustment preserves the original 1,778-test denominator by treating unavailable tests as $p = 1$ during adjustment. Reading error and coverage together shows the incremental effect of prioritization. Visualization code was prepared with OpenAI Codex assistance. Underlying data are provided in Supplementary Tables S6, S18 and S19.

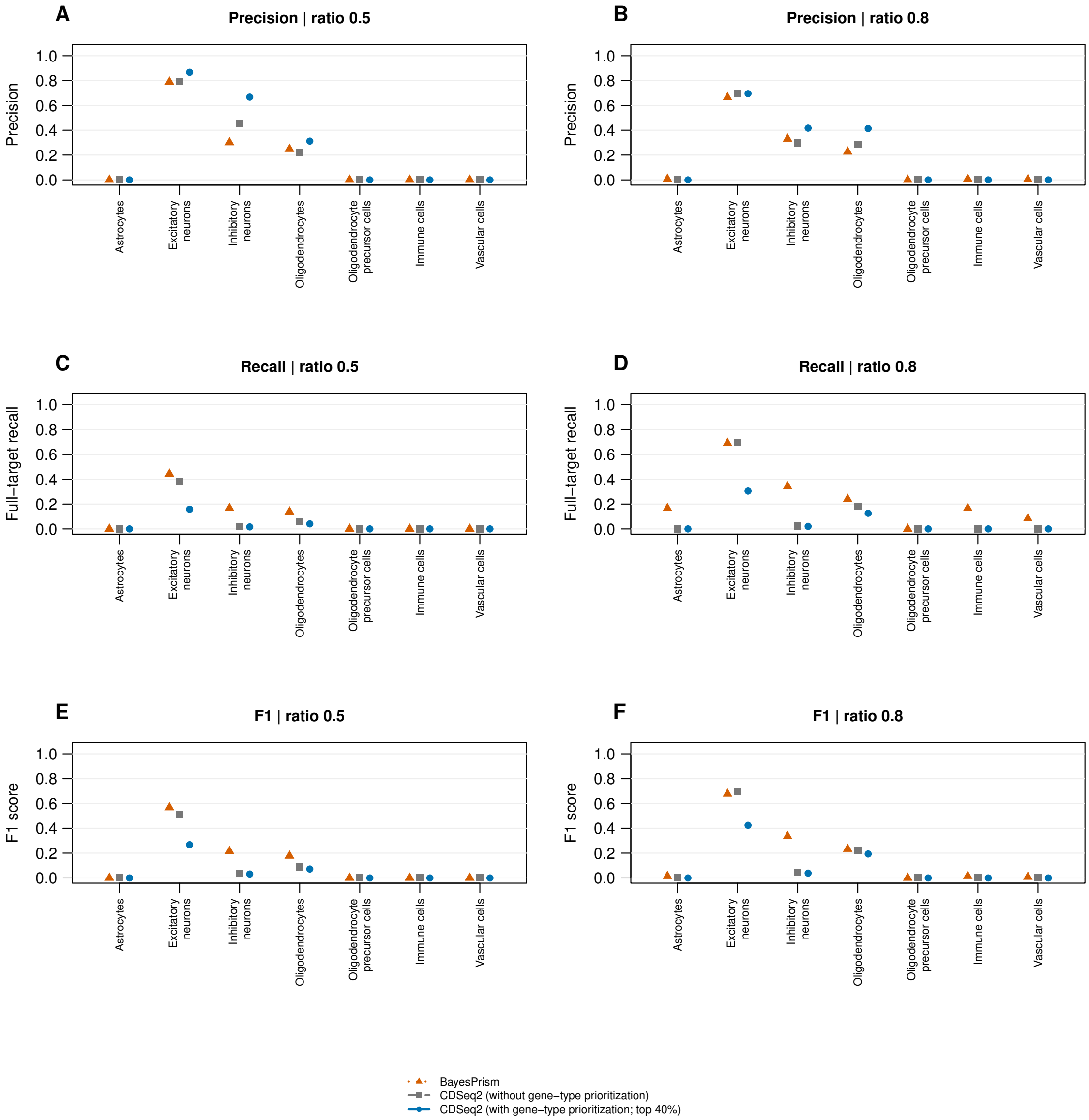


**Figure S9: Precision, recall and F1 across all seven cell types.** Columns show signal ratios 0.5 and 0.8. Rows show precision (A,B), recall of all injected targets (C,D) and F1 (E,F), each at the original BH-adjusted $q \leq 0.05$ threshold. Gray squares, blue circles and orange triangles denote CDSeq2 (without gene-type prioritization), CDSeq2 (with gene-type prioritization) and standard BayesPrism without output selection. Counts pool the three repetitions. Correct discovery requires the injected target and direction; all other calls are errors. Recall includes discarded and unavailable injected targets in its denominator. Undefined precision for a workflow with no calls is omitted; zero recall and F1 are retained. These panels show cell-type variation in performance and avoid interpreting precision without coverage. Repetitions share 40 donors. Visualization code was prepared with OpenAI Codex assistance. Underlying data are provided in Supplementary Table S21.

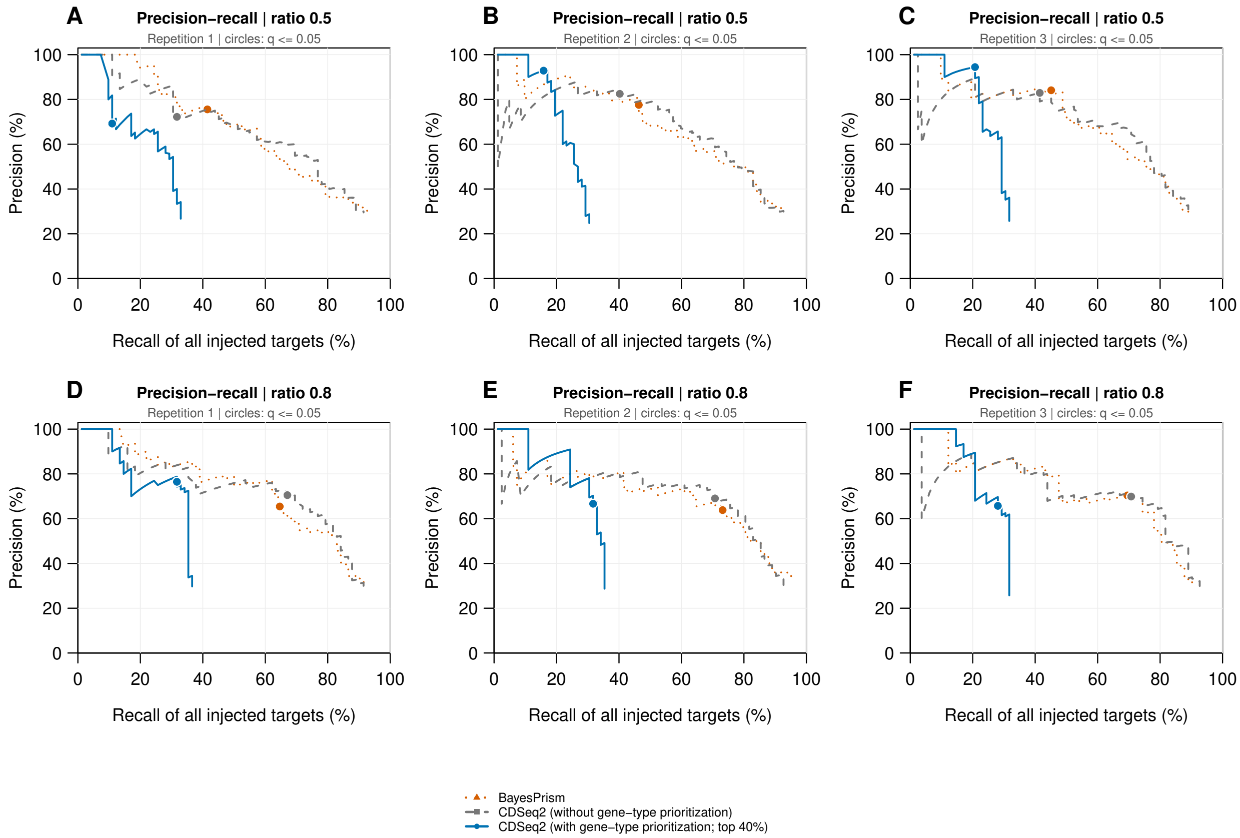


**Figure S10: Excitatory-neuron precision–recall curves in each simulation repetition.** Rows show signal ratios 0.5 (A–C) and 0.8 (D–F); columns show repetitions 1–3. Gray dashed, blue solid and orange dotted curves denote CDSeq2 (without gene-type prioritization), CDSeq2 (with gene-type prioritization) and standard BayesPrism without output selection. Circles mark the original $q \leq 0.05$ threshold. Each curve sweeps distinct original BH-adjusted values, with all ties entering together and no recomputation after selection; connecting lines are visual guides. Recall uses all 82 injected excitatory-neuron targets in that repetition, including discarded or unavailable targets. Correct calls require the target and injected sign. The full testing family contains 1,778 gene/type tests per scenario. All repetitions use the same 40 donors, so these panels describe simulation variability conditional on the cohort. Visualization code was prepared with OpenAI Codex assistance. Underlying data are provided in Supplementary Tables S20 and S21.

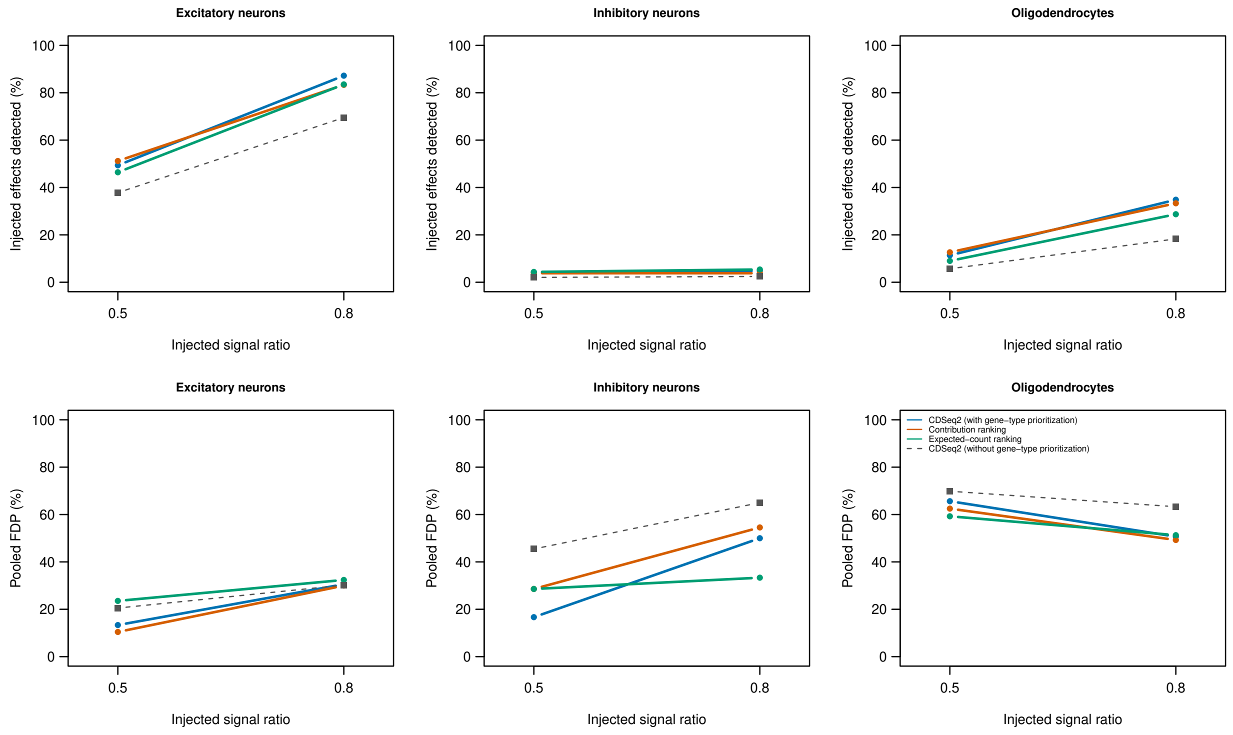


**Figure S11: Detection and attribution in nominated eQTL simulations.** Columns show excitatory neurons, inhibitory neurons and oligodendrocytes. Top, correct-sign detections divided by selected injected tests. Bottom, non-injected gene/type calls divided by all selected calls. Colored curves use post-injection rankings and a matched top-40% nominated-test budget; the dashed curve is CDSeq2 (without gene-type prioritization). Counts pool three repetitions at each signal ratio, using the same 40 donors. Selection changes denominators and does not recompute BH-adjusted values. A high selected-set detection rate is not equivalent to high recall of all injected tests. Visualization code was prepared with OpenAI Codex assistance. Underlying data are provided in Supplementary Table S5.

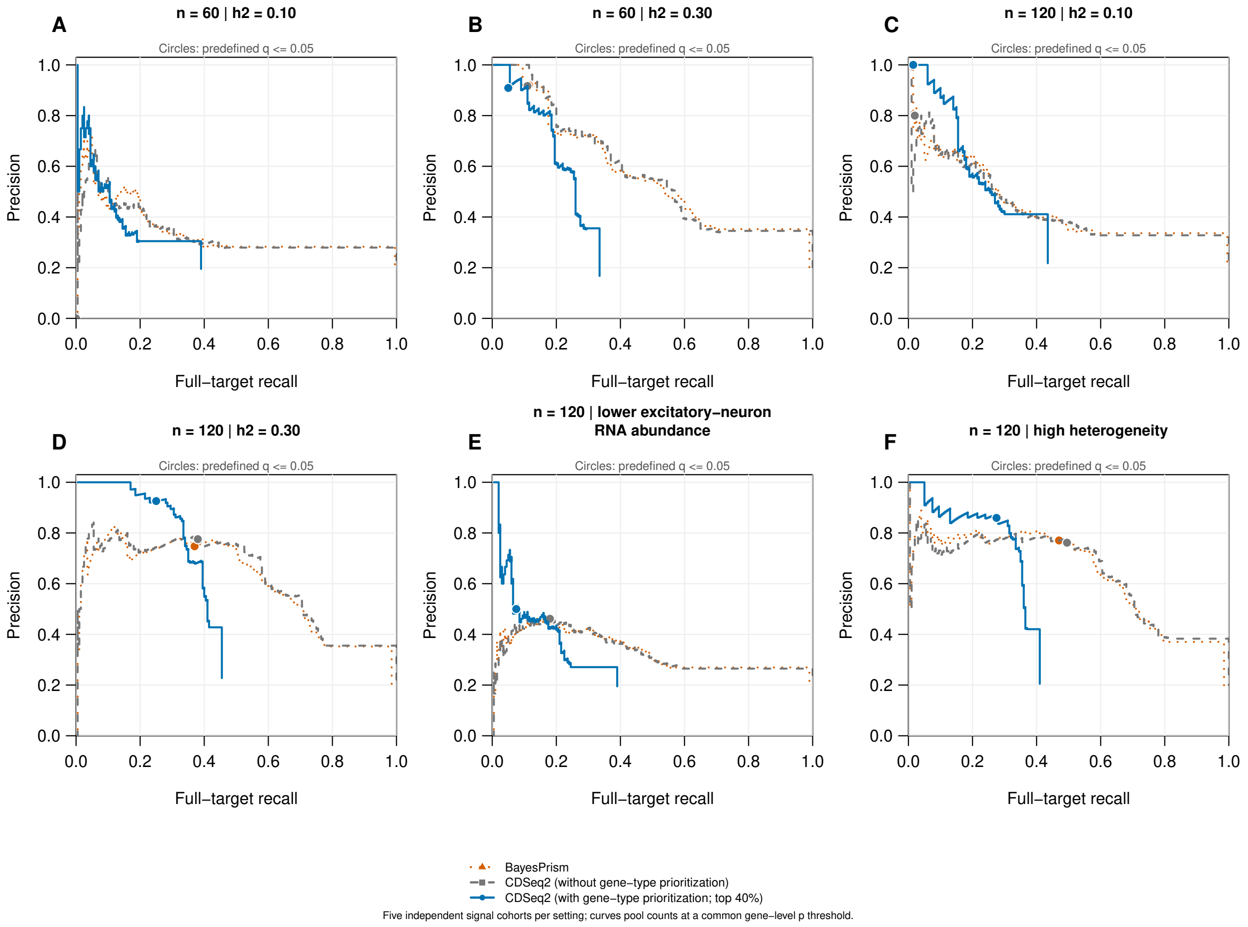


**Figure S12: Excitatory-neuron precision–recall in all six LD-aware simulation settings.** Each panel pools counts from five independent synthetic cohorts at common gene-level $p$ thresholds. The six settings vary sample size, nominal pre-normalization genetic variance fraction, excitatory-neuron abundance and expression heterogeneity. Orange, gray and blue represent standard BayesPrism, CDSeq2 (without gene-type prioritization) and CDSeq2 (with gene-type prioritization), respectively. Circles show the fixed $q \leq 0.05$ operating points; cohort-specific BH thresholds mean these circles need not lie exactly on a curve defined by one common unadjusted threshold. All ties enter together, and steps describe attainable pooled decisions. The blue workflow retains 80 of 200 targets per type without using genotype or test truth to choose membership. Recall always includes all 200 injected excitatory-neuron targets across the five cohorts, including discarded or unavailable pairs. Curves are descriptive; the primary threshold and retention are frozen. Visualization code was prepared with OpenAI Codex assistance. Underlying data are provided in Supplementary Tables S26, S28–S30 and S35.

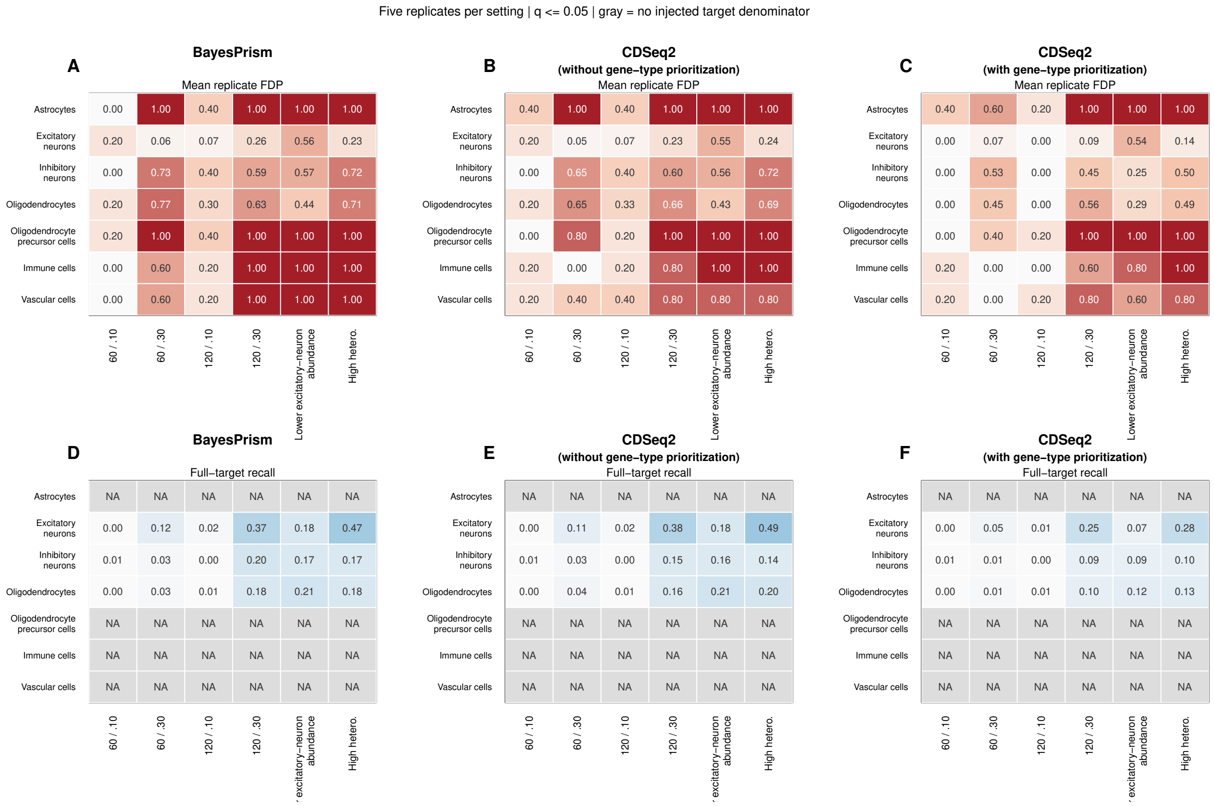


**Figure S13: Cell-type dependence of LD-aware association performance.** Columns compare standard BayesPrism, CDSeq2 (without gene-type prioritization) and CDSeq2 (with gene-type prioritization). Rows show (A–C) mean replicate FDP and (D–F) mean full-target recall at $q \leq 0.05$. Within each heat map, rows are cell types and columns are the six simulation settings. Values average five independent synthetic cohorts; color scales span zero to one, with darker red indicating larger FDP and darker blue indicating greater recall. Only excitatory neurons, inhibitory neurons and oligodendrocytes receive injected effects. Recall is therefore undefined, rather than zero, in the other types (gray cells). Their significant calls count as false direct-target assignments. A zero FDP can reflect no discoveries and must be interpreted with the call counts in Table S29. The full 1,400-pair family is retained before optional prioritization. Visualization code was prepared with OpenAI Codex assistance. Underlying data are provided in Supplementary Tables S26, S28, S29 and S35.

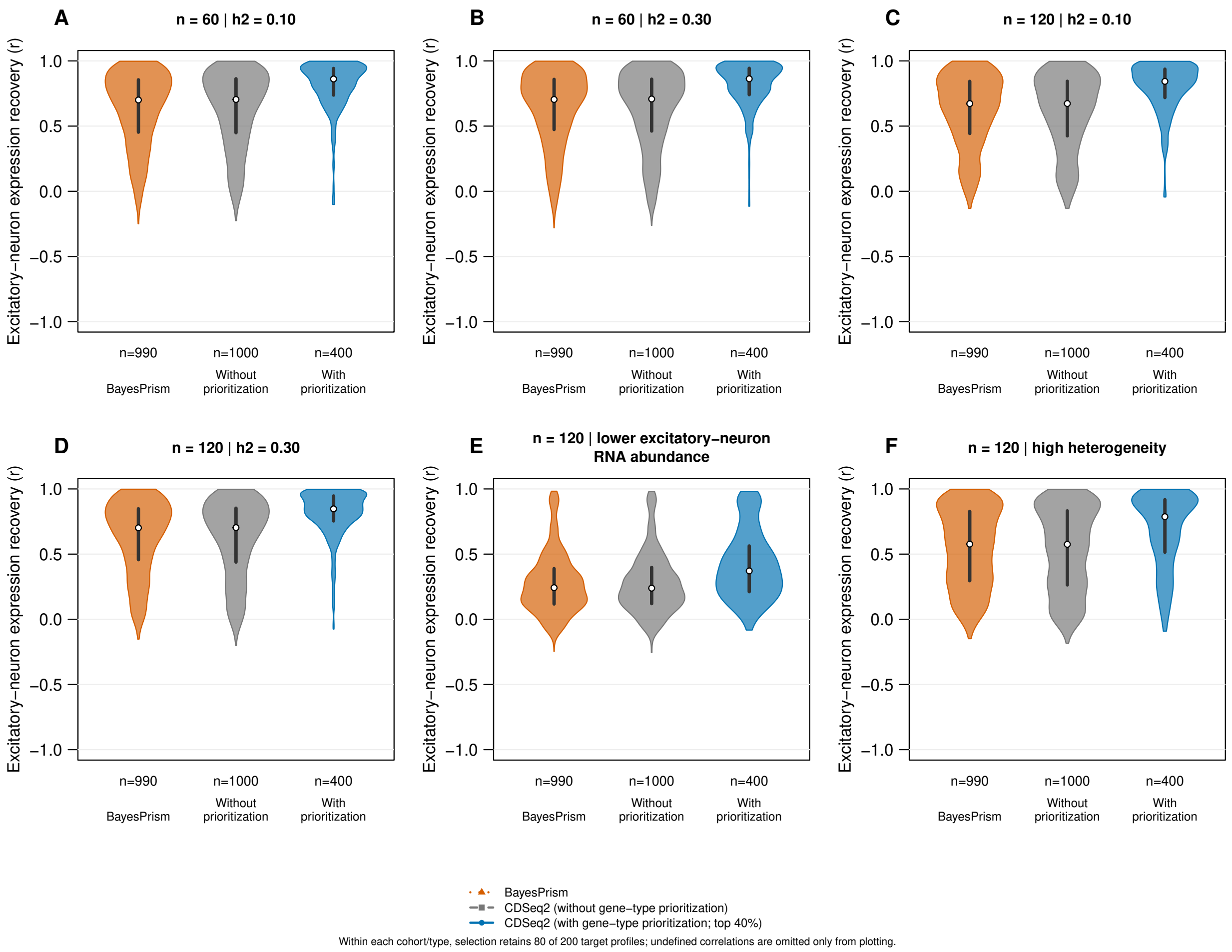


**Figure S14: Expression recovery and prioritization in the new LD-aware cohorts.** Each violin displays across-donor Pearson correlations between inferred and realized observed-component CPM for excitatory-neuron targets, pooled over five independent cohorts within a setting. The relative-expression target matches the original scorer-training definition. Table S32 additionally reports correlations on the log-expression scale used for association testing. Orange and gray include every available BayesPrism and CDSeq2 target profile; blue shows CDSeq2's frozen top-40% subset. Standard workflows have 1,000 candidate target/cohort profiles per setting, and prioritization retains 400. Displayed $n$ values count defined correlations, after method-specific availability and zero-library checks. Selection precedes removal of undefined correlations and never uses observed-component truth. White dots indicate medians and dark segments indicate interquartile ranges; equal maximum violin widths show distributional shape, not sample size. Densities are truncated at observed correlation ranges. The same target genes recur across cohorts; profiles within a cohort share individuals. These are target-universe results, not an assessment of all 2,000 modeled genes. Visualization code was prepared with OpenAI Codex assistance. Underlying data are provided in Supplementary Tables S26, S28, S32 and S33.

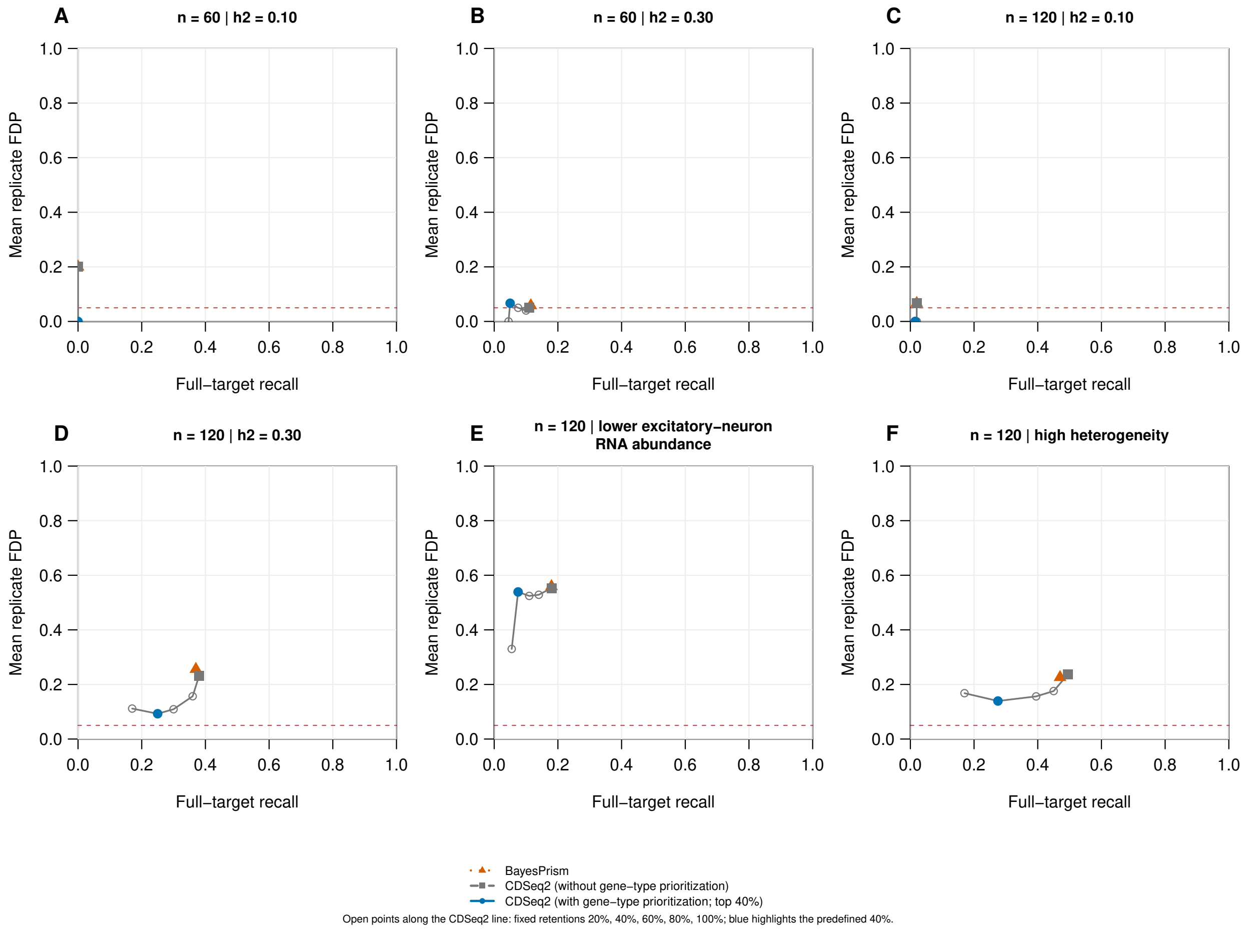


**Figure S15: Predefined retention sensitivity at the fixed significance threshold.** Each panel shows excitatory-neuron full-target recall against mean replicate FDP at $q \leq 0.05$, averaging five independent cohorts. Open points connected by a gray line show CDSeq2 retention levels 20%, 40%, 60%, 80% and 100%; the filled blue point highlights the frozen primary 40% rule, and the gray square marks CDSeq2 without prioritization. The orange triangle shows standard BayesPrism with native input preprocessing and no output selection. Selection never changes CDSeq2 expression, association tests or full-family BH adjustment. Dashed horizontal lines mark 0.05 as a reference. Lower FDP with lower recall represents an accuracy–coverage tradeoff, and a zero FDP may reflect no calls. These sensitivities were specified before evaluation and do not select a new operating point from test truth. Visualization code was prepared with OpenAI Codex assistance. Underlying data are provided in Supplementary Tables S26, S28, S29 and S35.

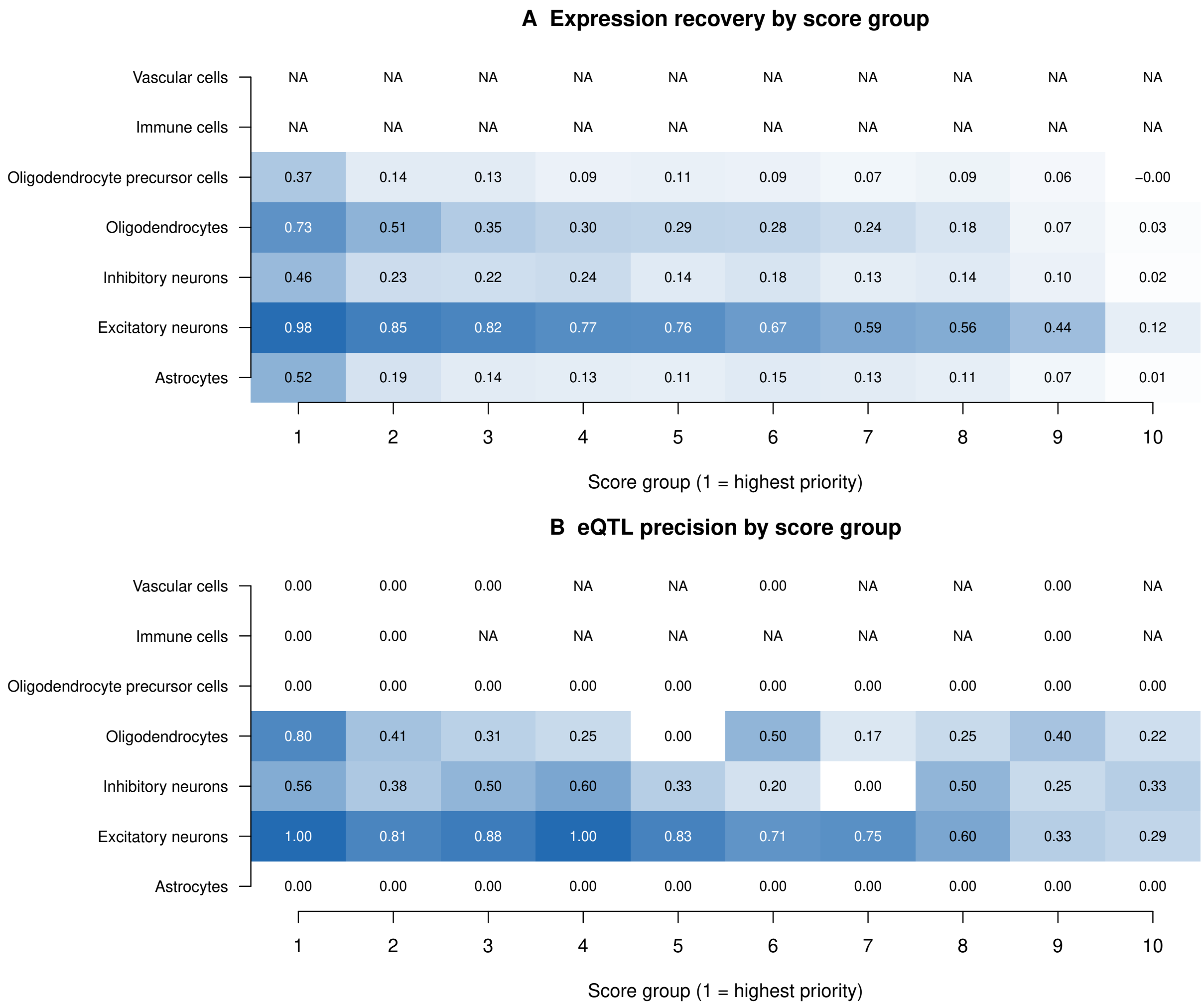


**Figure S16: Score groups across all seven cell types.** Panels show mean cohort median expression recovery and pooled injected-target precision at the fixed full-family significance threshold, for the principal independent-cohort setting. Rows are cell types and columns are score groups; group 1 has highest priority. Each group contains 20 candidate profiles per cohort. Numeric values are the estimates; cohort-bootstrap intervals are provided in Table S41. Only three types have injected effects; precision is undefined for zero calls and zero for nonempty call sets with no injected targets. The displays do not imply a calibrated probability or monotonic improvement in every type. Visualization code was prepared with OpenAI Codex assistance. Underlying data are provided in Supplementary Tables S36 and S41.

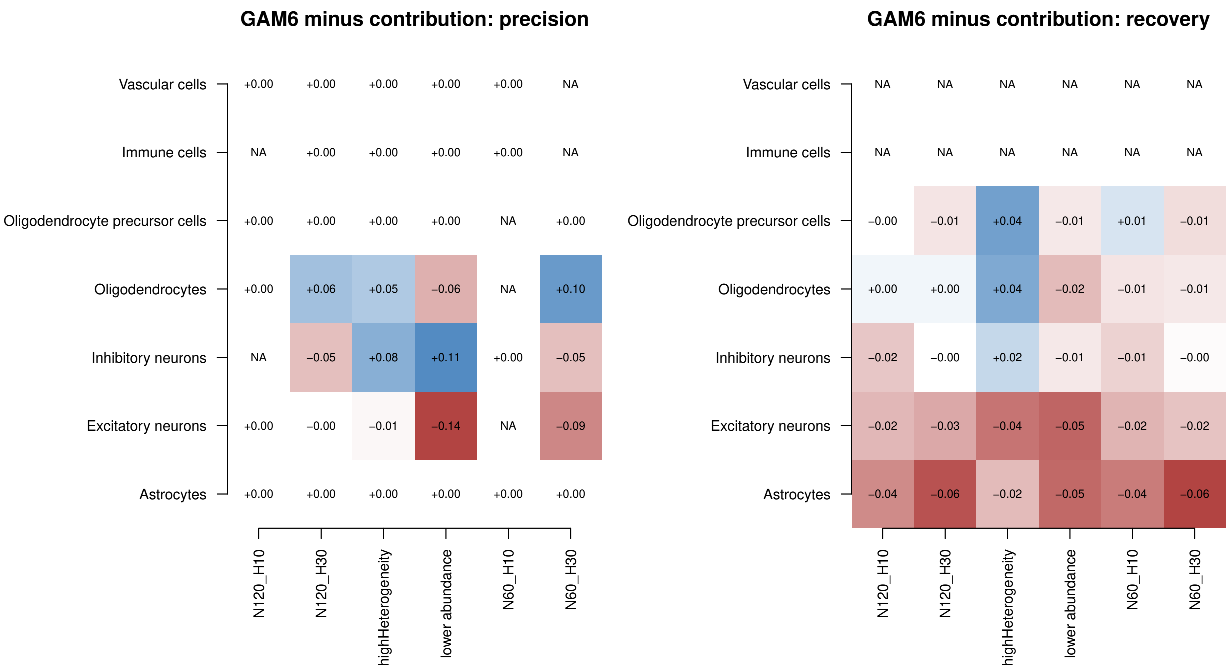


**Figure S17: Learned and contribution ranking across simulation settings.** Each cell reports GAM6 minus contribution ranking at the same 40% retention. Blue is positive, red negative and NA undefined. Left shows pooled injected-target precision; right shows mean cohort median expression recovery. Numeric values are unstandardized differences; each panel uses its own symmetric color scale. All six settings and seven types are included. Paired uncertainty and expected-count and random controls are available in the tables. Visualization code was prepared with OpenAI Codex assistance. Underlying data are provided in Supplementary Tables S37 and S38.

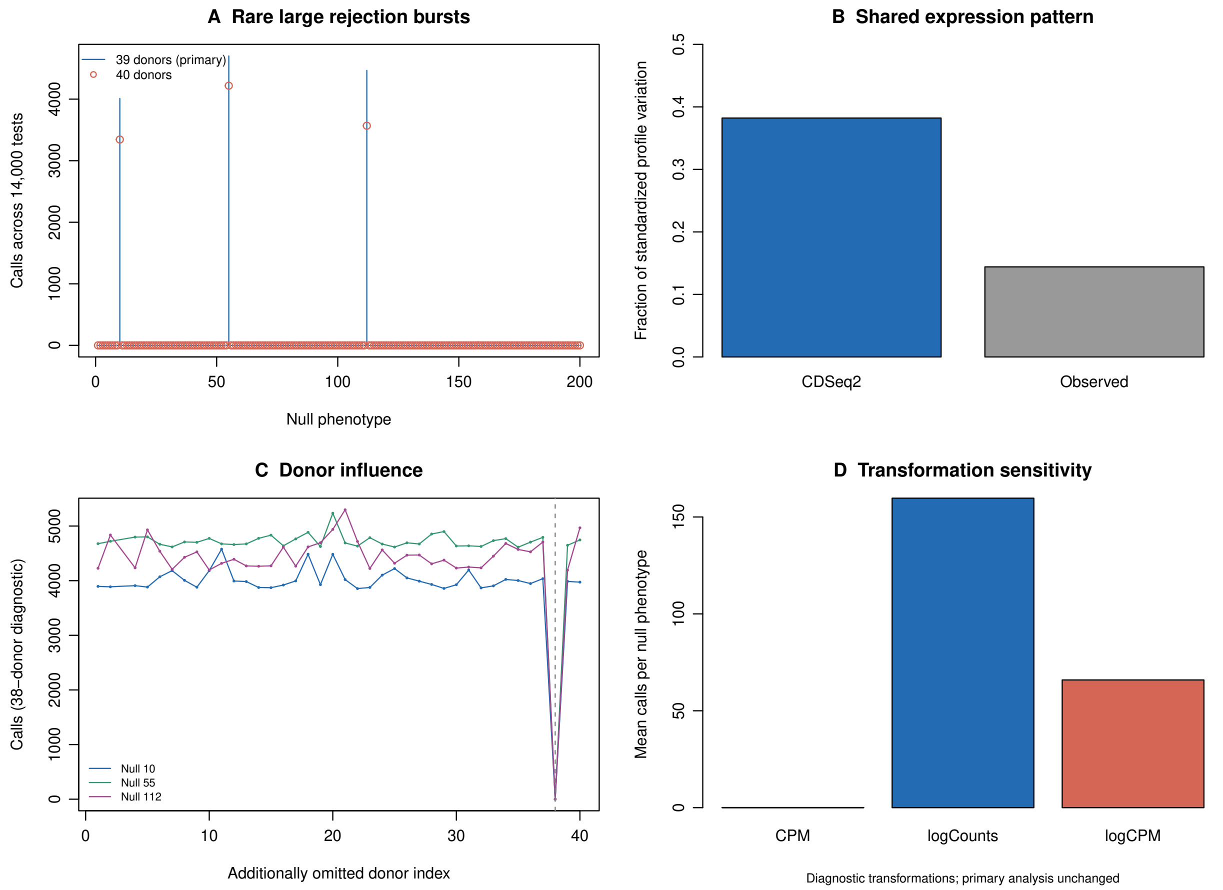


**Figure S18: Expression structure and continuous-null sensitivity.** (A) Calls across 200 null phenotypes under the primary 39-donor and full 40-donor log-CPM analyses. (B) The fraction of variation explained by the first principal component after row centering and unit-norm scaling. (C) Additional donor omissions for the three burst phenotypes; donor 38 eliminates their calls, but no additional omission is applied to the primary analysis. (D) Mean null calls under three response transformations, with the same 39 donors and fixed full testing family. These are post-analysis stability diagnostics, not outcome-selected preprocessing recommendations. Visualization code was prepared with OpenAI Codex assistance. Underlying data are provided in Supplementary Tables S39–S40 and S42.
