## Additional file 2: Supplementary Tables S1-S11 for "CDSeq2 integrates individual-level cell-type expression deconvolution with recoverability-guided prioritization": README.txt

Additional file 2: Profile recovery and association benchmark data
Tables S1--S11: profile scores, retention curves, continuous-phenotype summaries, nominated-eQTL results, design and recovery medians.

CSV contents are unchanged from the v16/0.1.0 archived release.
Table numbers and meanings are defined in Additional file 1.
Historical S6 labels in S23/S24 filenames refer to the current Figure S4C.
Software and complete reproduction bundle: https://doi.org/10.5281/zenodo.23489824

Table S1: S1_profile_predictions.csv
Table S2: S2_recovery_curves.csv
Table S3: S3_continuous_summary.csv
Table S4: S4_null_results.csv
Table S5: S5_eqtl_summary.csv
Table S6: S6_standard_comparison.csv
Table S7: S7_eqtl_leakage.csv
Table S8: S8_eqtl_tests.csv
Table S9: S9_continuous_replicates.csv
Table S10: S10_eqtl_design.csv
Table S11: S11_profile_medians.csv
