## Additional file 3: Supplementary Tables S12-S19 for "CDSeq2 integrates individual-level cell-type expression deconvolution with recoverability-guided prioritization": README.txt

Additional file 3: Ground-truth accuracy and displayed application data
Tables S12--S19: RNA-fraction and expression accuracy, continuous examples, eQTL plot data and BayesPrism preprocessing.

CSV contents are unchanged from the v16/0.1.0 archived release.
Table numbers and meanings are defined in Additional file 1.
Historical S6 labels in S23/S24 filenames refer to the current Figure S4C.
Software and complete reproduction bundle: https://doi.org/10.5281/zenodo.23489824

Table S12: S12_ground_truth_accuracy.csv
Table S13: S13_RNA_fraction_pairs.csv
Table S14: S14_expression_accuracy.csv
Table S15: S15_eqtl_observed_agreement.csv
Table S16: S16_continuous_slope_recovery.csv
Table S17: S17_continuous_donor_example.csv
Table S18: S18_eqtl_before_after.csv
Table S19: S19_BayesPrism_preprocessing.csv
