## Additional file 4: Supplementary Tables S20-S25 for "CDSeq2 integrates individual-level cell-type expression deconvolution with recoverability-guided prioritization": README.txt

Additional file 4: Precision, recall and prioritization sensitivity data
Tables S20--S25: precision--recall curves, F1, matched empirical FDP caps, selection membership and donor sensitivity.

CSV contents are unchanged from the v16/0.1.0 archived release.
Table numbers and meanings are defined in Additional file 1.
Historical S6 labels in S23/S24 filenames refer to the current Figure S4C.
Software and complete reproduction bundle: https://doi.org/10.5281/zenodo.23489824

Table S20: S20_precision_recall_curves.csv
Table S21: S21_precision_recall_F1.csv
Table S22: S22_matched_empirical_FDP.csv
Table S23: S23_S6_GAM6_membership.csv
Table S24: S24_S6_recovery_counts.csv
Table S25: S25_continuous_donor_sensitivity.csv
