## Additional file 5: Supplementary Tables S26-S35 for "CDSeq2 integrates individual-level cell-type expression deconvolution with recoverability-guided prioritization": README.txt

Additional file 5: Independent-cohort simulation benchmark data
Tables S26--S35: LD-aware design, haplotypes, tests, accuracy, null results, expression recovery, preprocessing and injection design.

CSV contents are unchanged from the v16/0.1.0 archived release.
Table numbers and meanings are defined in Additional file 1.
Historical S6 labels in S23/S24 filenames refer to the current Figure S4C.
Software and complete reproduction bundle: https://doi.org/10.5281/zenodo.23489824

Table S26: S26_LD_cohort_design.csv
Table S27: S27_LD_reference_windows.csv
Table S28: S28_LD_gene_type_tests.csv
Table S29: S29_LD_accuracy_coverage.csv
Table S30: S30_LD_precision_recall.csv
Table S31: S31_LD_global_null.csv
Table S32: S32_LD_expression_recovery.csv
Table S33: S33_LD_BayesPrism_preprocessing.csv
Table S34: S34_LD_retrospective_FDP_caps.csv
Table S35: S35_LD_injection_design.csv
