## Additional file 6: Supplementary Tables S36-S42 for "CDSeq2 integrates individual-level cell-type expression deconvolution with recoverability-guided prioritization": README.txt

Additional file 6: Prioritization controls and association stability data
Tables S36--S42: score groups, equal-budget controls, uncertainty, null sensitivity and donor expression support.

CSV contents are unchanged from the v16/0.1.0 archived release.
Table numbers and meanings are defined in Additional file 1.
Historical S6 labels in S23/S24 filenames refer to the current Figure S4C.
Software and complete reproduction bundle: https://doi.org/10.5281/zenodo.23489824

Table S36: S36_score_deciles.csv
Table S37: S37_matched_retention_controls.csv
Table S38: S38_control_uncertainty.csv
Table S39: S39_null_stability.csv
Table S40: S40_null_profile_diagnostics.csv
Table S41: S41_score_decile_uncertainty.csv
Table S42: S42_donor_expression_support.csv
